# Bandwidth-aware fusion of resting-state EEG–fMRI connectivity in cortical eigenmode space

**DOI:** 10.64898/2026.05.15.725408

**Authors:** Hyung G. Park

## Abstract

We introduce a *geometry-informed, bandwidth-aware* model for fusing resting-state EEG and fMRI connectivity by treating each modality as a masked, physics-limited observation of a shared latent cortical connectivity structure. Both modalities are represented in a shared cortical Laplace–Beltrami (LB) eigenmode basis that provides an anatomically grounded ordering of spatial scales. In this coordinate system, fMRI constrains connectivity across a broad range of spatial scales but only at slow timescales, whereas EEG provides frequency-resolved information primarily for coarse spatial modes. We formalize this complementarity through a latent spatio-spectral connectivity object and estimate it using a low-rank factorization with shared eigenmode-network factors and subject- and frequency-specific nonnegative strengths. Estimation is driven by the regions supported by each modality (EEG: coarse spatial modes across broad frequencies; fMRI: broad spatial modes at low frequencies), while unobserved bands are inferred through shared low-rank structure and spectral smoothness regularization. In the MPI–LEMON cohort, the fused representation yields compact, interpretable subject features (fMRI network strengths and EEG oscillatory summaries) that support out-of-sample age prediction under nested cross-validation, and coherent multi-view interpretation through cortical maps, spectral profiles, and atlas-referenced system composition summaries. These results demonstrate that explicitly modeling modality-specific bandwidth gaps enables principled EEG–fMRI connectivity fusion and provides a practical route to multimodal network biomarkers for individual-differences research.

## 1. Introduction

Resting-state electroencephalography (EEG) and functional MRI (fMRI) offer complementary views of large-scale brain connectivity: fMRI provides whole-cortex, spatially detailed network organization, whereas EEG provides frequency-resolved coupling and fast neural dynamics (Biswal et al., 1995; Smith et al., 2009; Yeo et al., 2011; Buzsáki and Draguhn, 2004; Fries, 2005; Hipp et al., 2012). A unified representation of these signals could provide a more complete account of large-scale brain organization than either modality alone. Despite extensive interest in relating these modalities (Mantini et al., 2007; Sadaghiani and Wirsich, 2020; Sadaghiani et al., 2022), principled *connectivity-level* fusion remains challenging because the natural connectivity objects differ (low-frequency rs-fMRI connectomes versus frequency-resolved EEG connectivity) and each modality is fundamentally bandwidth-limited by measurement physics (Boynton et al., 1996; Logothetis, 2008; Cordes et al., 2001; Nunez and Srinivasan, 2006; Baillet, 2017; Michel and Brunet, 2019; Colclough et al., 2015). This work addresses that gap by introducing a geometry-informed, bandwidth-aware model for integrating resting-state EEG and fMRI connectivity in a shared cortical coordinate system (Pang et al., 2023; Park, 2025).

No single noninvasive modality simultaneously provides whole-cortex coverage, high spatial specificity, and direct access to fast neural dynamics. In rs-fMRI, signals are temporally filtered by hemodynamics and neurovascular coupling and are therefore dominated by slow fluctuations (Boynton et al., 1996; Logothetis, 2008; Cordes et al., 2001). EEG, in contrast, provides millisecond-scale neural dynamics and frequency-resolved oscillatory structure that support large-scale coordination (Buzsáki and Draguhn, 2004; Fries, 2005; Siegel et al., 2012; Hipp et al., 2012). Yet EEG connectivity is inherently bandwidth-limited in space: volume conduction and the ill-posed inverse problem restrict the reliably identifiable spatial degrees of freedom, and residual spatial mixing can bias connectivity even with standard leakage-mitigation approaches (Nunez and Srinivasan, 2006; Baillet, 2017; Michel and Brunet, 2019; Colclough et al., 2015). These modality-specific limits induce *structured missing bandwidth*—regions of a joint spatio-temporal description that are unobserved due to measurement physics rather than missing at random—that the other modality can help constrain.

This complementarity motivates a core question: *Can we build a principled and interpretable fusion model that represents resting-state connectivity from fMRI and EEG in a shared cortical coordinate system, while explicitly accounting for modality-specific bandwidth limits and leveraging complementary strengths of fMRI (spatial coverage/specificity) and EEG (spectral resolution)?*

Cross-modal studies have established reproducible links between electrophysiological signatures and fMRI-derived resting-state networks, including correlations between band-limited EEG power (or network-level electrophysiological features) and fMRI network structure/dynamics (Mantini et al., 2007; Sadaghiani and Wirsich, 2020; Sadaghiani et al., 2022). However, practical fusion for individual-differences analysis remains challenging because EEG–fMRI integration often proceeds either by collapsing EEG into a small set of canonical-band summaries or by expanding to multiband/multitimescale feature sets (including connectome-level decompositions), which can complicate interpretability and statistical stability in moderate sample sizes (Deligianni et al., 2014; Wirsich et al., 2020a, 2021, 2020a). Connectivity estimates are also high-dimensional, and their reliability can vary across edges, preprocessing choices, and spatial scales, motivating representations that are both statistically stable and interpretable for downstream individual-differences analyses (Finn et al., 2015; Cole and Franke, 2017; Noble et al., 2019; Schaefer et al., 2018).

Here we introduce a low-rank resting-state connectivity fusion framework built on a common coordinate system indexed by: (i) a spatial-scale axis defined by cortical eigenmodes (Laplace–Beltrami eigenfunctions) ordered by eigenvalue (coarse-to-fine spatial frequency) (Reuter et al., 2009; Lévy, 2006; Pang et al., 2023), and (ii) a temporal frequency (spectral) axis *ω*. Let *k* ∈ {1, …, *K*} denote the leading *K* eigenmodes (low *k* = coarse spatial scales; high *k* = fine scales). In the resulting (*k, ω*) (spatial–spectral) plane, EEG and fMRI provide complementary sampling windows: EEG yields frequency-resolved connectivity over a broad *ω* range but is informative primarily at coarse spatial scales (low *k*), whereas fMRI constrains connectivity over a broad spatial bandwidth (many *k*) but effectively only at very low temporal frequencies due to hemodynamic low-pass filtering (Boynton et al., 1996; Logothetis, 2008). Consequently, the high-*k* EEG spatial region and the high-*ω* fMRI spectral region are not merely noisy—they are *structurally unobserved* given the physics of each measurement modality. This multiscale coordinate-system view is also consistent with broader efforts to relate brain structure and function in low-dimensional descriptions of macroscale brain organization (Suárez et al., 2020).

### Latent-field fusion in eigenmode coordinates

This perspective suggests a direct modeling principle: represent resting-state connectivity as a latent spatio-spectral field and view each modality as a noisy, masked observation of that common object. Specifically, let Σ_*i*_(*k, k*^′^; *ω*) denote subject *i*’s latent, frequency-resolved mode-by-mode coupling matrix in cortical eigenmode space. The key challenge is that neither modality identifies the latent field Σ_*i*_(*k, k*^′^; *ω*) everywhere in (*k, ω*). Fusion then becomes a constrained inference problem: the observed EEG and fMRI bands jointly inform the spatio-spectral field Σ_*i*_ on their respective supports, while the structurally unobserved regions must be inferred through a structural model that couples the observed bands and regularizes the structurally unobserved regions.

### A low-rank, positive semi-definite (PSD), frequency-resolved network factorization

We impose a parameterization that is both parsimonious and PSD-by-construction:

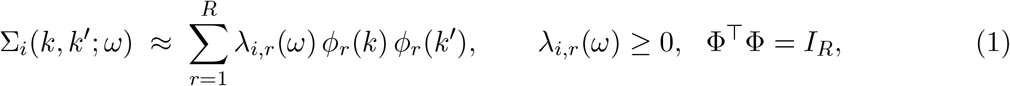

where 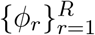 are shared spatial network factors in eigenmode coordinates and 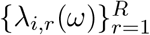 are subject- and frequency-specific nonnegative network strengths. This factorization acts as a *compressor* : it replaces the high-dimensional object Σ_*i*_(*k, k*^′^; *ω*) with a small set of interpretable subject-level summaries while coupling modalities through a shared network subspace span(Φ). Model estimation is driven by entries supported by each modality; structurally unobserved bands are inferred indirectly through the shared low-rank structure together with smoothness regularization over *ω*.

### Relation to existing multimodal fusion

A large literature integrates multimodal neuroimaging via feature-level fusion, regression-based integration, and multi-view predictive modeling (Sui et al., 2012; Calhoun and Sui, 2016; Jorge et al., 2014). Within EEG–fMRI connectomics, common strategies relate rs-fMRI connectivity to band-specific EEG summaries or to multiband/multi-timescale EEG connectomes (Deligianni et al., 2014; Wirsich et al., 2021, 2020a). Latent-variable approaches (e.g., linked/joint decompositions) can learn shared components across modalities and accommodate missingness (Groves et al., 2011; Sadaghiani and Wirsich, 2020; Wirsich et al., 2020b), but typically operate in ROI/voxel feature spaces and do not explicitly encode a geometry-defined, coarse-to-fine *spatial-scale axis* or represent *frequency-resolved* EEG connectivity within a single shared connectivity object.

Our formulation differs in that it places both modalities in spatial-frequency-ordered cortical eigenmode coordinates and treats multimodal fusion as inference on a shared spatio-spectral connectivity field. The resulting low-rank representation yields a single network coordinate system with subject- and frequency-specific strengths, making *scale–frequency organization* an explicit estimand while producing low-dimensional features that are interpretable and stable for downstream prediction and association analyses.

### Contributions

Our approach makes three contributions (Fig. 2). First, it introduces a geometry-defined spatial-scale axis for multimodal connectomics by expressing EEG and fMRI connectivity in a shared Laplace–Beltrami eigenmode basis, where spatial bandwidth is explicit and ordered. Second, it formulates EEG–fMRI fusion as inference under *structured missing bandwidth*: each modality provides a physics-limited sampling window in the (*k, ω*) plane, and model fitting is driven by modality-supported regions while structurally unobserved bands are inferred via shared low-rank structure and smoothness regularization. Third, it yields a single, PSD-by-construction, frequency-resolved network representation with shared eigenmode-network factors and subject- and frequency-specific nonnegative strengths, enabling interpretable scale–frequency signatures and low-dimensional subject features for individual-differences analyses.

### Application to MPI–LEMON

We demonstrate the framework in the MPI–LEMON (Max Planck Institute Leipzig Mind-Body-Brain Interactions) cohort, which provides resting fMRI and resting EEG from the same participants across young and older adults (Babayan et al., 2019). Age serves as a useful validation target because it is associated with well-replicated changes in both rs-fMRI connectivity (e.g., altered network segregation and default-mode vulnerability) and resting EEG spectral organization (e.g., reduced alpha-band activity and shifts in periodic/aperiodic structure) (Scally et al., 2018; Donoghue et al., 2020; Yang et al., 2023; Merkin et al., 2023). In MPI–LEMON, we show that fused network features yield strong out-of-sample prediction of age, robust age associations in compact spatial and spectral summaries, and coherent scale–frequency structure across the learned networks.

The remainder of the paper is organized as follows. Section 2 presents the model and estimation procedure. Section 3 evaluates estimation behavior and effective-rank recovery in simulations. Section 4 applies the method to MPI–LEMON. Section 5 discusses implications, limitations, and future directions.

## 2. Methods

### 2.1. Cortical eigenmodes as a shared spatial basis

All analyses are performed in a common cortical coordinate system defined by the Laplace–Beltrami (LB) eigenmodes of a template cortical surface. Let 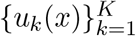 denote the first *K* LB eigenfunctions on the template cortical surface *x* ∈ ℳ, ordered by increasing LB eigenvalue *κ*_*k*_ (increasing spatial frequency; i.e., decreasing spatial scale) (Reuter et al., 2006; Pang et al., 2023). Any sufficiently smooth cortical field *f* (*x, t*) at time *t* can be approximated by the truncated expansion

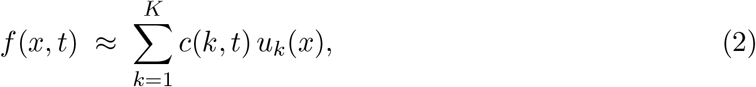

where *c*(*k, t*) is the coefficient of the *k*th eigenmode at time *t*.

For rs-fMRI, we project each subject’s preprocessed blood-oxygenation-level-dependent (BOLD) time series onto the LB basis to obtain LB-domain coefficient time-series *b*_*i*_(*k, t*) For EEG, we perform source reconstruction and represent the resulting cortical activity in the same LB basis, yielding LB-domain coefficient time-series *a*_*i*_(*k, t*). Throughout, low LB indices (small *k*) correspond to coarse spatial patterns that are comparatively well-resolved by EEG, whereas higher LB indices (large *k*) correspond to finer spatial patterns that are less reliably identifiable from EEG due to sensor-space mixing and inverse ill-posedness.

Figure 1 provides an illustrative example of this shared multiscale indexing for one participant. Selected eigenmodes *u*_*k*_(*x*) (*x* ∈ ℳ) provide a common multiscale spatial coordinate (here *k* ∈ {1, 10, 20} for illustration), while the effective temporal bandwidths differ by modality: fMRI LB coefficients *b*_*i*_(*k, t*) reflect slow, hemodynamically filtered fluctuations (middle panel), whereas EEG LB coefficients *a*_*i*_(*k, t*) exhibit fast, oscillatory dynamics (right panel).

**Figure 1:**
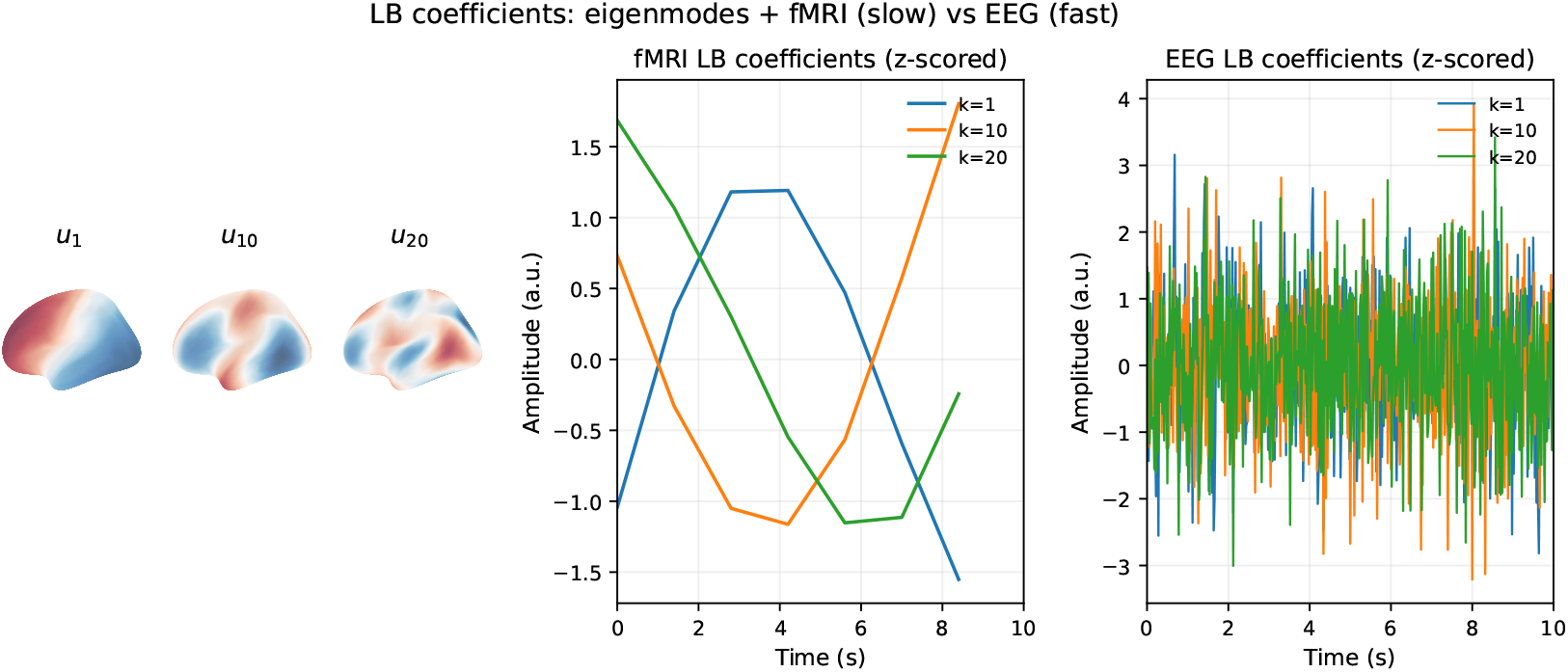
Illustration of a shared cortical eigenmode basis across modalities. Left: selected Laplace–Beltrami eigenmodes *u*_*k*_ (ordered by increasing spatial frequency) visualized on the `fsaverage5` left hemisphere (lateral view) (here *k* ∈ {1, 10, 20} for illustration). Middle: the corresponding rs-fMRI LB coefficient time series *b*_*i*_(*k, t*) (*k* = 1, 10, 20) for the same participant over a short segment, showing slow, hemodynamically filtered fluctuations. Right: the corresponding EEG LB coefficient time series *a*_*i*_(*k, t*) (*k* = 1, 10, 20) over a representative short segment, showing faster oscillatory dynamics. Because fMRI provides higher effective spatial bandwidth than EEG, higher-*k* (finer-scale) eigenmode coefficients are typically more reliably constrained by fMRI, whereas EEG predominantly constrains lower-*k* (coarser-scale) components; the shared eigenmode indexing enables direct cross-modal comparison within a common multiscale coordinate system.

### 2.2. Latent spatio-spectral connectivity field

We model resting-state connectivity via a latent spatio-spectral connectivity field in LB space. For subject *i*, let *X*_*i*_(*k, t*) denote an unobserved latent activity process in the LB domain. Our target is a *frequency-indexed, mode-by-mode coupling field* Σ_*i*_(*k, k*^′^; *ω*) defined over LB mode indices (*k, k*^′^) and temporal frequency *ω*. When *X*_*i*_ is approximately second-order stationary, Σ_*i*_ coincides with the cross-spectral density (CSD),

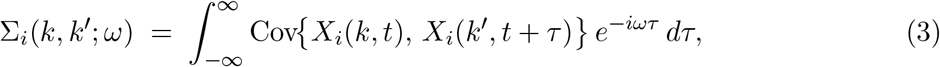

for *k, k*^′^ ∈ {1, …, *K*} and angular frequency *ω*, which provides the canonical second-order spectral target in the stationary case.

In practice, the observed connectivity objects are band-limited or aggregated covariance-type summaries. After source reconstruction, EEG connectivity is constructed from band-limited, leakage-robust amplitude-envelope covariance (AECov), whereas rs-fMRI connectivity is summarized by the low-frequency covariance of LB-projected BOLD coefficients. We work with covariance rather than correlation because scale-dependent variance is itself informative in the LB domain. These observed connectivities are treated as modality-specific readouts of the underlying latent field Σ_*i*_(*k, k*^′^; *ω*). Accordingly, Σ_*i*_(*k, k*^′^; *ω*) serves as a latent frequency-indexed coupling field whose observations may be band-limited, aggregated, and real-valued.

### 2.3. Low-rank eigen-network factorization

To obtain a parsimonious connectivity representation and ensure positive semi-definiteness, we impose a low-rank factorization (introduced in Eq. (1)) of the latent connectivity field:

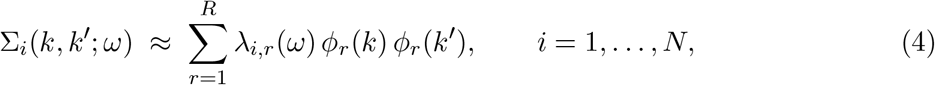

where *ϕ*_*r*_(*k*) is the *r*th spatial network factor in LB space, shared across subjects and modalities; *λ*_*i,r*_(*ω*) ≥ 0 is the subject- and frequency-specific strength of network *r*; and *R* ≪ *K* is the number of network factors.

We constrain the spatial factors *ϕ*_*r*_ to be orthonormal, 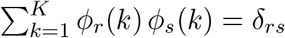, equivalently Φ^⊤^Φ = *I*_*R*_ with Φ = [*ϕ*_1_, …, *ϕ*_*R*_] ∈ ℝ^*K*×*R*^. For numerical implementation, we represent *λ*_*i,r*_(*ω*) on a discrete frequency grid 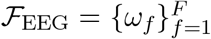 and regularize the spectral profile to be smooth in *ω* (Section 2.5).

Under (4), each Σ_*i*_(*ω*) is positive semi-definite (PSD) for every *ω*, and the full spatio-spectral structure is captured by the low-dimensional factors {*ϕ*_*r*_, *λ*_*i,r*_(·)}.

#### 2.3.1. fMRI connectivity (full spatial bandwidth; low-frequency summary)

For subject *i*, we compute LB-domain BOLD coefficients *b*_*i*_(*k, t*) and define the empirical rs-fMRI connectivity as the sample covariance across time,

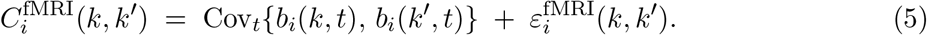

Because BOLD fluctuations primarily reflect slow temporal components after standard denoising and band-limited preprocessing, 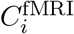 can be interpreted as a low-*ω* summary of the latent spatio-spectral connectivity field. Under the low-rank factorization (4), this summary is parametrized by

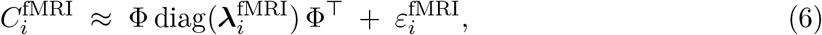

where 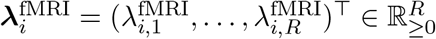 is the subject–network strength vector and Φ ∈ ℝ^*K*×*R*^ is the shared spatial factor matrix coupling fMRI and EEG. Equivalently, 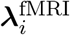 may be viewed as a low-frequency summary of the latent frequency-resolved strengths {*λ*_*i,r*_(*ω*)} in (4), but we do not require an explicit discretization of the fMRI band for estimation.

#### 2.3.2. EEG connectivity (restricted spatial bandwidth; frequency-resolved)

For EEG, we compute frequency-resolved functional connectivity matrices in LB space using a leakage-robust covariance-type summary. To maximize comparability with the covariance-based rs-fMRI connectome in (5), we use band-limited amplitude-envelope covariance (AECov) as the EEG connectivity object (Supplementary Methods Section S5.4). Briefly, source-localized LB coefficient time series are band-pass filtered into *F* frequency bins, and symmetric orthogonalization is applied to the band-limited signals to reduce instantaneous linear mixing (Colclough et al., 2016). Amplitude envelopes are then obtained via the Hilbert transform, and connectivity is defined as the covariance of these envelopes, averaged across time windows to yield a single subject-level matrix per frequency bin. Let 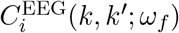 denote the empirical EEG connectivity at frequency bin *ω*_*f*_.

EEG connectivity is informative primarily for coarse spatial modes (low *k*), which we encode via a mode-wise reliability weight, *η*_EEG_(*k*) ∈ [0, 1], and the induced edge mask

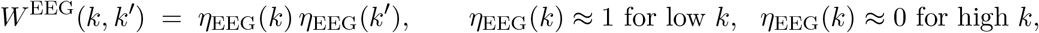

implemented either as a hard cutoff 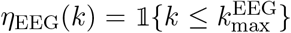 or as a smooth taper. We model EEG connectivity as a noisy masked readout of the latent field:

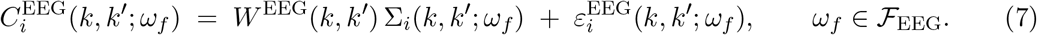

Then, under the factorization (4), the EEG observations depend on the shared spatial factors through their low-*k masked* projections: 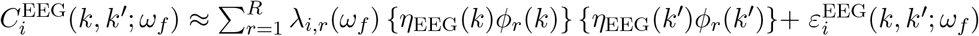. Thus, EEG primarily constrains the frequency dependence *λ*_*i,r*_(*ω*) over ℱ_EEG_ = {*ω*_*f*_} and the coarse-scale (low-*k*) components of the shared spatial network factors.

### 2.4. Missing bandwidths as structured latent connectivity

The fMRI and EEG observation models imply that the two modalities probe complementary subsets of the latent spatio-spectral connectivity field. Specifically, rs-fMRI provides a noisy low-*ω* summary of Σ_*i*_(*k, k*^′^; *ω*) across the full spatial bandwidth *k, k*^′^ ∈ {1, …, *K*} (model (5)), whereas EEG provides frequency-resolved observations over *ω* ∈ ℱ_EEG_ but primarily on edges supported by coarse spatial modes (low *k*), as encoded by the mask *W* ^EEG^(*k, k*^′^) (model (7)).

We formalize this complementarity via modality-specific support masks *W* ^fMRI^(*k, k*^′^; *ω*) and *W* ^EEG^(*k, k*^′^; *ω*) that are nonzero only where the corresponding modality provides meaningful information. For any (*k, k*^′^, *ω*) where both masks are zero, Σ_*i*_(*k, k*^′^; *ω*) is *structurally unobserved* —missing due to the modalities’ measurement physics rather than missing at random. Rather than imputing such entries independently, we treat them as latent and infer them indirectly through the shared low-rank structure (4), smoothness regularization on the spectral weights *λ*_*i,r*_(*ω*), and the joint constraints imposed by the observed bands from both modalities. Figure 2 provides a schematic overview.

**Figure 2:**
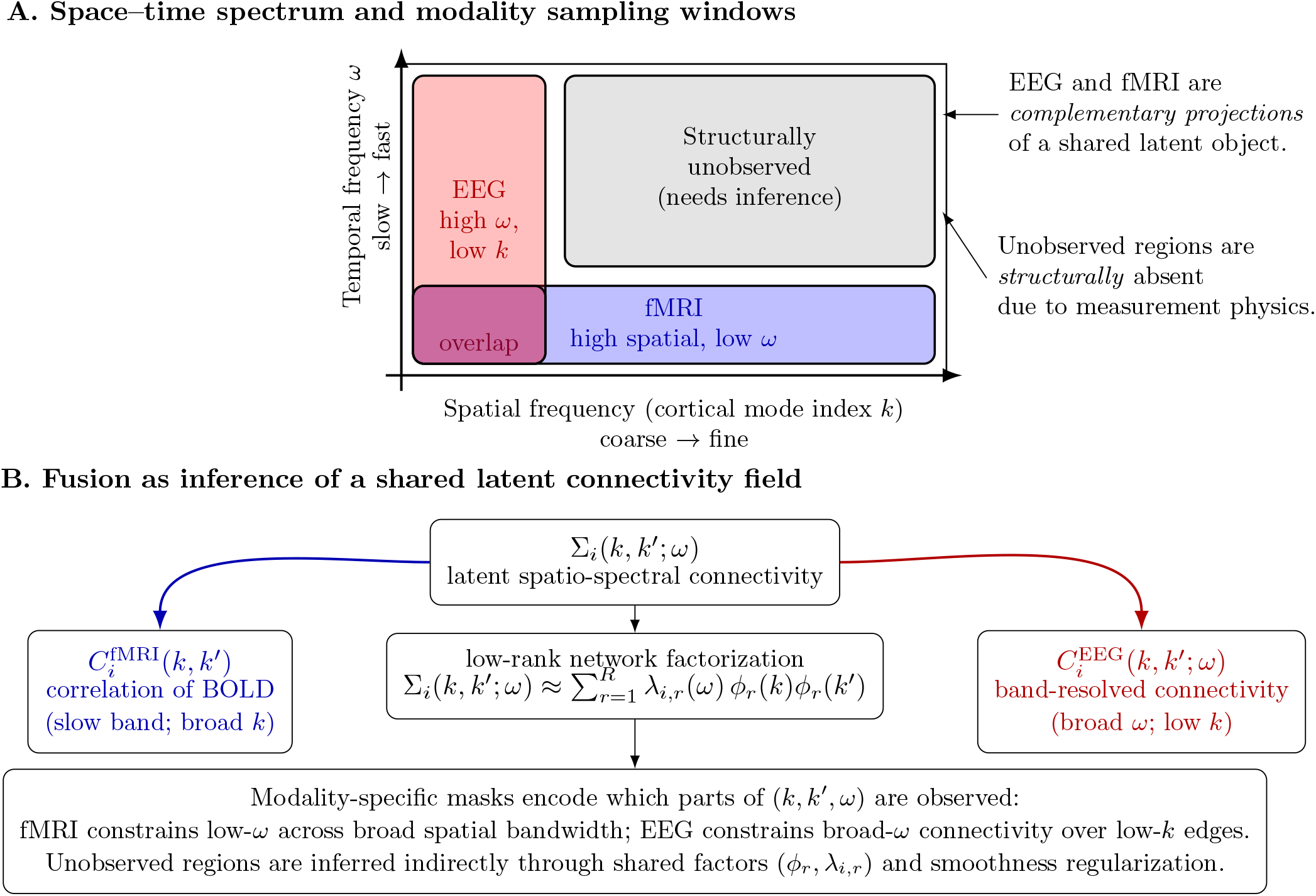
Conceptual diagram of EEG–fMRI connectivity fusion. (A) Brain dynamics can be viewed in a joint space–time-frequency spectrum. fMRI and EEG provide complementary, band-limited views, leaving structurally unobserved regions that cannot be directly measured by either modality. (B) Fusion is posed as inference of a shared latent spatio-spectral connectivity field, constrained jointly by both modalities via a low-rank network factorization and modality-specific observation masks.

### 2.5. Penalized maximum a posteriori (MAP) estimation

Let Θ = (Φ, *λ, λ*^fMRI^) collect all model parameters, where Φ = [*ϕ*_1_, …, *ϕ*_*R*_] ∈ ℝ^*K*×*R*^ satisfies Φ^⊤^ Φ = *I*_*R*_, 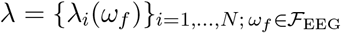 with 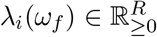, and 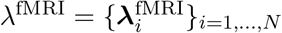 with 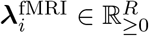. We assume Gaussian observation models for the empirical connectivity matrices on their observed support, as encoded by modality-specific masks.

#### EEG likelihood

For each subject *i* and frequency bin *ω*_*f*_ ∈ ℱ_EEG_, we model

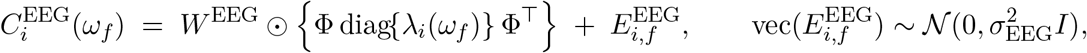

where Φ diag{*λ*_*i*_(*ω*_*f*_)} Φ^⊤^ is the model-implied frequency-indexed connectivity matrix in LB space in (4).

#### fMRI likelihood

We model rs-fMRI connectivity as a low-frequency summary that constrains the same shared spatial factors Φ through a nonnegative subject–network strength vector 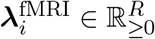:

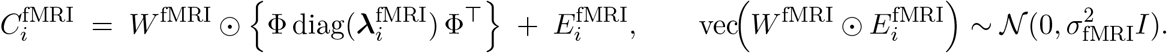

Conceptually, 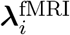 can be interpreted as a low-*ω* aggregation of {*λ*_*i*_(*ω*)} over a small set of BOLD-relevant frequencies, but in practice, to avoid imposing a literal correspondence between EEG frequency bins and the BOLD band, we treat 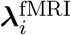 as free nonnegative parameters.

#### Penalties (priors)

We regularize the spectral weights to encourage smooth frequency profiles. For each subject *i* and factor *r*, we impose a second-order difference penalty on *λ*_*i,r*_(*ω*_*f*_) across *ω*_*f*_ ∈ ℱ_EEG_ together with the nonnegativity constraint *λ*_*i,r*_(*ω*_*f*_) ≥ 0:

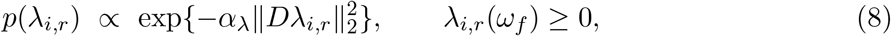

where *D* is the (*F* − 2) × *F* second-difference matrix on the frequency grid, (*Dλ*)_*f*_ = *λ*_*f*+2_ − 2*λ*_*f*+1_ + *λ*_*f*_.

#### Objective (penalized MAP)

We estimate 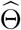 by maximizing the posterior 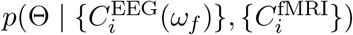, equivalently by minimizing the negative log-posterior (up to constants):

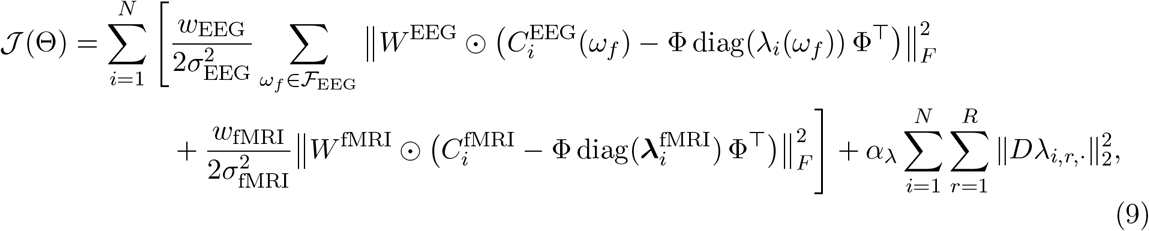

where *λ*_*i,r*,·_ = (*λ*_*i,r*_(*ω*_1_), …, *λ*_*i,r*_(*ω*_*F*_))^⊤^ stacks the spectral weights over the EEG frequency grid. We set the modality-specific weights *w*_fMRI_ = 1 and *w*_EEG_ = 1*/F*, so that the EEG contribution is averaged over frequency bins and is comparable in scale to the single fMRI term.

### 2.6. ARD shrinkage and effective rank

To enable automatic effective-rank selection, we use an automatic relevance determination (ARD) regularization strategy (Mørup and Hansen, 2009; Tipping, 2001). We fit the model with a moderately large upper bound *R*_max_ and allow the data to deactivate unnecessary factors through factor-specific shrinkage on the spectral profiles {*λ*_*i,r*_(*ω*_*f*_)}.

#### ARD penalty

When ARD is enabled, we add to the objective (9) the factor-wise quadratic penalty

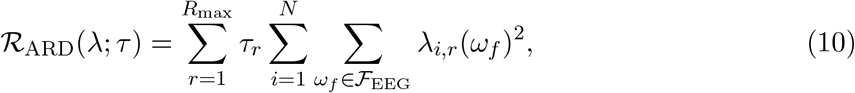

where *τ*_*r*_ *>* 0 controls shrinkage of the component *r* across subjects and frequencies. Intuitively, the components with little support in the data are driven toward *λ*_*i,r*_(*ω*_*f*_) ≈ 0.

#### Empirical-Bayes update of τ_r_

The ARD penalty depends on component-wise precision parameters *τ*_*r*_. Motivated by a Gaussian prior on *λ*_*i,r*_(*ω*_*f*_) with a Gamma hyperprior on *τ*_*r*_, we can update

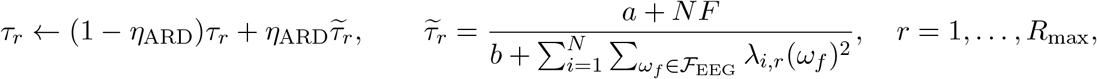

with small hyperparameters (e.g., *a* = *b* = 10^−6^) and truncation to [*τ*_min_, *τ*_max_]. This update increases *τ*_*r*_ when factor *r* has low energy (shrinking it more strongly) and decreases *τ*_*r*_ when the factor is used substantially. This update is not required for fixed-rank estimation; it is used only when we want the model to adaptively deactivate factors and determine an effective rank (see simulations).

#### Effective rank

After fitting with *R*_max_, we summarize factor usage by the average energy 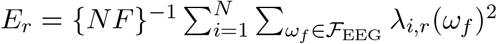 and define the effective rank as 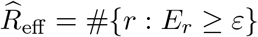. In this paper we set *ε* using a relative rule, *E*_*r*_*/* max_*r*_*′ E*_*r*_*′* ≥ 10^−2^, and confirm that results are robust across a small range of thresholds. In real data, we found the empirical-Bayes *τ* updates can be unstable; therefore we use a stability-first approach by freezing *τ*_*r*_ (i.e., setting *η*_ARD_ = 0) and computing 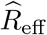 from the fitted energy curve.

### 2.7. Optimization algorithm

We minimize the objective (9) (optionally with the ARD penalty (10)) by alternating minimization over (i) subject-specific spectral weights and (ii) shared spatial factors. Specifically, we iterate: (A) update the nonnegative spectral profiles {*λ*_*i*_(*ω*_*f*_)} given Φ, (B) update the orthonormal spatial networks Φ given {*λ*_*i*_(*ω*_*f*_)} via a Stiefel-constrained gradient step, and when ARD is enabled, (C) update the ARD precision parameters *τ* (Section 2.6).

#### 2.7.1. (A) Update λ given Φ

Fixing Φ, the objective decouples by subject and is convex in the stacked vector of spectral weights 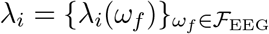, with nonnegativity constraints and a quadratic smoothness penalty across frequency (and optionally ARD shrinkage). For each subject *i*, we solve a non-negative penalized least-squares problem of the form

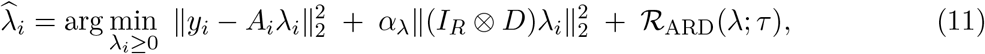

where *y*_*i*_ and *A*_*i*_ are obtained by vectorizing the masked empirical connectivities (EEG across frequencies and fMRI once) and expressing the model-implied matrices under the current Φ. We solve (11) using bounded-variable least squares (nonnegative least squares on an augmented system); implementation details are provided in Supplementary Methods (Section S1.2).

#### 2.7.2. (B) Update Φ given λ (Stiefel-constrained descent)

Fixing {*λ*_*i*_(*ω*_*f*_)}, we update Φ under the orthonormality constraint Φ^⊤^Φ = *I*_*R*_ by performing a gradient step on the Stiefel manifold followed by a QR retraction. Concretely, we compute the Euclidean gradient *G* = ∇_Φ_𝒥, project it onto the tangent space at the current iterate, take a step with backtracking line search, and retract back to the feasible set via QR factorization. This procedure enforces Φ^⊤^Φ = *I*_*R*_ at every iteration and yields stable optimization in practice; full formulas are given in Supplementary Methods (Section S1.3).

#### 2.7.3. Convergence and outputs

We iterate (A)–(B) until the relative decrease in 𝒥 falls below a tolerance (e.g., 10^−5^) (or a maximum number of iterations is reached). The final estimates define the fitted latent-field matrices 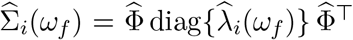 and fMRI network strength vector 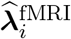. For downstream analyses we compute compact subject-level features, including (i) fMRI network strengths 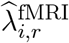, (ii) EEG band-mass summaries per network, and (iii) an EEG spectral centroid per network (Hz-weighted center of mass across the modeling frequency grid).

## 3. Simulation study

### 3.1. Simulation design and data generation

We conducted simulations to assess whether the masked low-rank latent connectivity model can recover (i) the shared LB network subspace and (ii) subject-specific frequency profiles under conditions resembling MPI–LEMON (*n* ≈ 200). Throughout, we set the LB dimension to *K* = 50 and discretized the frequency axis on a grid 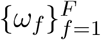 with *F* = 20 bins spanning the EEG frequency range Ω_EEG_. For simulation convenience, we also index the latent field on this same grid, while using a small index set ℱ_fMRI_ ⊂ {1, …, *F*} to represent *abstract* “slow” bins contributing to the fMRI summary; these indices are not interpreted as literal EEG frequencies.

#### Latent truth

In each Monte Carlo replicate we generated subject- and frequency-specific latent connectivity matrices using the same PSD low-rank form as the fitted model: 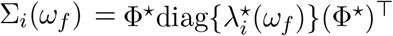, for *i* = 1, …, *n* and *f* = 1, …, *F*, with orthonormal Φ^⋆^ ∈ ℝ^*K*×*R*^ and nonnegative, smooth weights 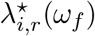. We generated Φ^⋆^ by drawing a *K* × *R* matrix with i.i.d. 𝒩(0, 1) entries followed by QR orthonormalization. For each factor *r*, we specified a smooth group-level spectral template *µ*_*r*_(*ω*) (with unimodal peaks distributed across *θ/α/β*-like ranges) and generated subject-specific log-weights using a low-dimensional spline basis:

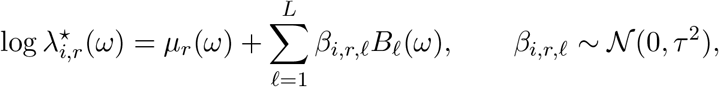

with *L* = 4 cubic B-splines and 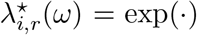 to enforce nonnegativity. This yields smooth variation over frequency while keeping the effective degrees of freedom modest.

#### Masking (missing bandwidths)

We encoded complementary modality bandwidth limits as follows.

1. **EEG spatial cutoff:** EEG observations were restricted to coarse spatial scales by masking to low LB modes *k*, 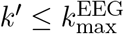, via an edge mask

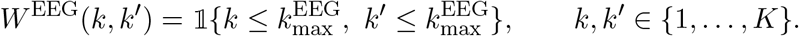
2. **EEG frequency coverage:** EEG observations were available for all bins *ω*_*f*_ (*f* = 1, …, *F*).
3. **fMRI spatial coverage and slow-band summary:** fMRI provided one full *K* × *K* connectivity matrix per subject (full spatial bandwidth), intended to represent a low-*ω* summary of the latent field. For simulation convenience we implemented this as an average over a small set of “slow” indices ℱ_fMRI_ ⊂ {1, …, *F*}: 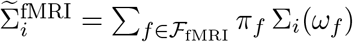, with |ℱ_fMRI_| = 4 and uniform weights *π*_*f*_ = 1*/*|ℱ_fMRI_|.

#### Observed data generation

Given the above masks, we generated observed connectivities by adding symmetric Gaussian noise:

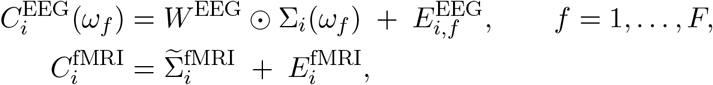

where 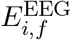 and 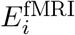 have i.i.d. 𝒩(0, *σ*^2^) entries on the upper triangle and are mirrored to ensure symmetry. We set *W* ^fMRI^ to the all-ones matrix (full spatial bandwidth).

### 3.2. Simulation scenarios and evaluation metrics

#### Simulation grid

We fixed *n* = 200. We varied the true rank *R*^*^ ∈ {5, 10}, the EEG spatial cutoff 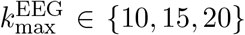, and the signal-to-noise ratio (SNR). For the oracle-rank accuracy study we used SNR ∈ {0.5, 1, 2, 4}, and for ARD rank recovery we used SNR ∈ {1, 2, 4}. We define SNR as the ratio of mean *masked* signal energy to mean *masked* noise energy for each modality, e.g.,

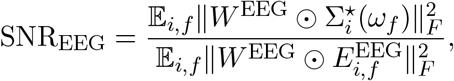

and analogously for fMRI. We set *σ*_EEG_ and *σ*_fMRI_ to achieve a common target SNR across modalities, so that SNR remains comparable as 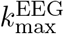 varies. We used *N* = 100 Monte Carlo replicates per grid cell for oracle-rank accuracy and *N* = 50 replicates per grid cell for ARD rank recovery.

For each simulated dataset we fit the penalized MAP estimator using the alternating minimization procedure described in Section 2.7. ARD rank-selection fits used *R*_max_ = 20 and defined an effective rank by thresholding the factor-energy curve (Section 2.6). Oracle-rank fits fixed the rank to the true value *R* = *R*^*^.

#### Evaluation metrics

We quantified recovery of (i) the shared spatial network subspace span(Φ^∗^), (ii) subject-specific spectral weights, and (iii) reconstruction accuracy on observed and structurally unobserved regions.

i. *Spatial subspace recovery*. Because Φ is identifiable only up to rotation within the rank-*R* subspace, we evaluate recovery at the subspace level using:
ii. **Projection-matrix discrepancy:** 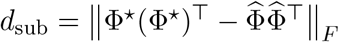.
  - **Principal angles:** letting 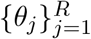 be the principal angles between span(Φ^⋆^) and span 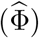, we report max_*j*_ *θ*_*j*_ and 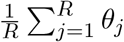 (degrees).
iii. *Spectral-weight recovery*. We report the root-mean-squared error (RMSE) of the frequency-resolved weights:

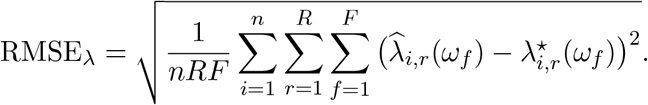
iv. *Reconstruction and extrapolation to structurally-missing regions*. We report relative error on: (1) the observed region ℛ_obs_ = {(*k, k*^′^, *f*) : *W* ^EEG^(*k, k*^′^) = 0 or *f* ∈ ℱ_fMRI_}; and on two structurally-missing regions: (2) EEG-high-*k* entries ℛ_EEG-miss_ = {(*k, k*^′^, *f*) : 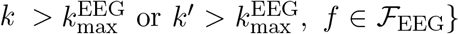; and (3) fMRI-high-frequency entries ℛ_fMRI-miss_ = {(*k, k*^′^, *f*) : *f* ∈*/* ℱ_fMRI_, (*k, 𝒦*^′^) ∈ 𝒦 ^2^}, where 𝒦 = {1, …, *K*} denotes the LB mode indices. For each subject *i* and region ℛ we compute

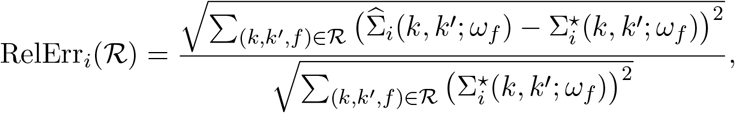

and summarize RelErr_*i*_(ℛ) across subjects by mean±SD.

### 3.3. Simulation results

We report two complementary evaluations. First, we assess rank recovery using an ARD-regularized fit with a generous working rank *R*_max_ = 20 and define an effective rank 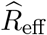 by thresholding the factor-energy curve at *E*_*r*_*/E*_max_ ≥ 0.01 (see Figures S6 and S7 in Section S8.1 for the results under the thresholds 0.005 and 0.02, respectively). Second, under the oracle setting *R* = *R*^*^, we evaluate estimation accuracy for the shared network subspace and reconstruction error on observed and structurally-missing regions.

#### 3.3.1. ARD effective-rank recovery

Figure 3 summarizes the distribution of 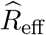 across Monte Carlo replicates for each scenario, stratified by true rank *R*^*^ and (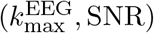. Table 1 provides the corresponding numerical summaries (median, mean (SD), exact recovery, and within-one recovery).

**Figure 3:**
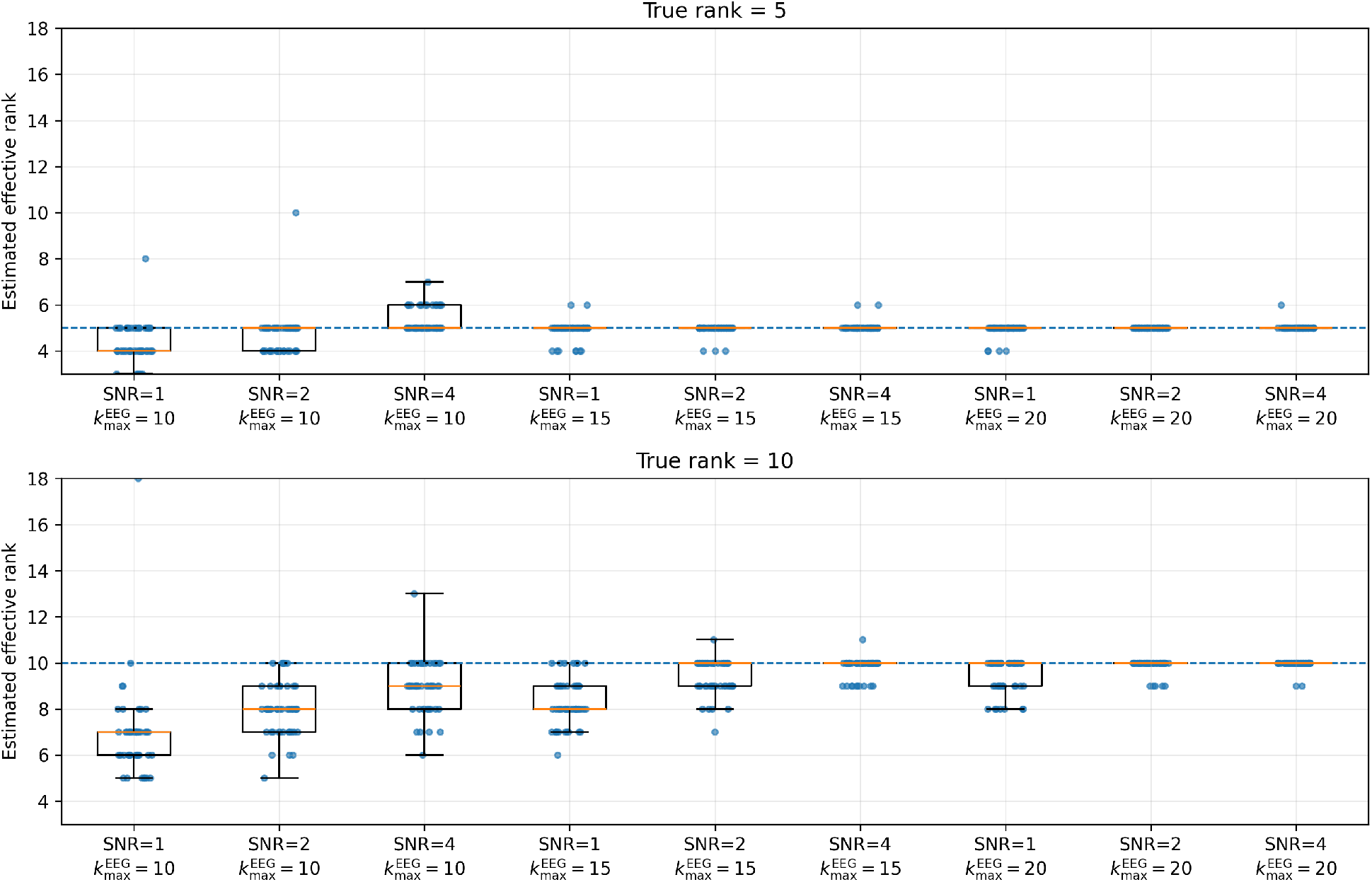
ARD effective-rank recovery. Boxplots show the distribution of the estimated effective rank 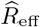 across Monte Carlo replicates for each (SNR, 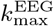) condition, stratified by true rank *R*^*\**^ ∈ {5, 10}. 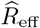 is defined by thresholding the factor-energy curve at *E*_*r*_*/E*_max_ ≥ 0.01 using an ARD-regularized fit with *R*_max_ = 20.

**Table 1:** ARD-based effective-rank recovery in the resting EEG–fMRI fusion simulation. For each scenario (true rank *R*^*\**^, EEG cutoff 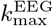, SNR), we report the distribution of the estimated effective rank 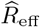 across *N* = 50 Monte Carlo replicates. 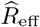 is defined by thresholding the factor-energy curve at 0.01 relative to the maximum energy (i.e., *E*_*r*_*/E*_max_ ≥ 0.01), using *R*_max_ = 20 and ARD regularization. Shown are median and mean (SD) of 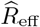, plus the exact recovery rate 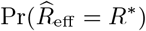 and the within-one rate 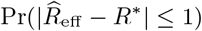.

| True rank $R^*$ | $(k_{\max}^{\text{EEG}}, \text{SNR})$ | Median | Mean (SD) | Exact | $\leq 1$ |
| --- | --- | --- | --- | --- | --- |
| 5 | (10, 1) | 4 | 4.34 (0.85) | 0.38 | 0.86 |
|  | (10, 2) | 5 | 4.74 (0.90) | 0.62 | 0.98 |
|  | (10, 4) | 5 | 5.30 (0.51) | 0.72 | 0.98 |
|  | (15, 1) | 5 | 4.90 (0.42) | 0.82 | 1.00 |
|  | (15, 2) | 5 | 4.94 (0.24) | 0.94 | 1.00 |
|  | (15, 4) | 5 | 5.04 (0.20) | 0.96 | 1.00 |
|  | (20, 1) | 5 | 4.92 (0.27) | 0.92 | 1.00 |
|  | (20, 2) | 5 | 5.00 (0.00) | 1.00 | 1.00 |
|  | (20, 4) | 5 | 5.02 (0.14) | 0.98 | 1.00 |
| 10 | (10, 1) | 7 | 6.90 (1.94) | 0.02 | 0.06 |
|  | (10, 2) | 8 | 8.02 (1.15) | 0.12 | 0.30 |
|  | (10, 4) | 9 | 8.96 (1.14) | 0.28 | 0.70 |
|  | (15, 1) | 8 | 8.28 (1.01) | 0.14 | 0.38 |
|  | (15, 2) | 10 | 9.42 (0.76) | 0.50 | 0.90 |
|  | (15, 4) | 10 | 9.82 (0.44) | 0.78 | 1.00 |
|  | (20, 1) | 10 | 9.44 (0.73) | 0.58 | 0.86 |
|  | (20, 2) | 10 | 9.90 (0.30) | 0.90 | 1.00 |
|  | (20, 4) | 10 | 9.96 (0.20) | 0.96 | 1.00 |
| 15 | (10, 1) | 8 | 8.52 (1.34) | 0.00 | 0.00 |
|  | (10, 2) | 11 | 10.68 (1.52) | 0.00 | 0.00 |
|  | (10, 4) | 12 | 11.96 (1.38) | 0.00 | 0.02 |
|  | (15, 1) | 11 | 10.78 (1.17) | 0.00 | 0.00 |
|  | (15, 2) | 13 | 12.90 (0.91) | 0.04 | 0.22 |
|  | (15, 4) | 14 | 14.08 (0.83) | 0.34 | 0.78 |
|  | (20, 1) | 13 | 12.76 (0.87) | 0.00 | 0.14 |
|  | (20, 2) | 14 | 14.18 (0.75) | 0.38 | 0.80 |
|  | (20, 4) | 15 | 14.78 (0.42) | 0.78 | 1.00 |

Across *R*^*^ ∈ {5, 10}, rank recovery improves monotonically as either EEG spatial support expands (larger 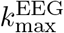) or SNR increases. For example, when *R*^*^ = 10, moving from (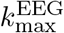, SNR) = (10, 1) to (20, 4) increases the exact recovery rate from 0.02 to 0.96 and the within-one rate from 0.06 to 1.00 (Table 1). When EEG covers at least the first 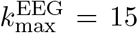 LB modes and SNR is moderate to high, ARD yields high exact recovery for *R*^*^ = 5 and strong within-one recovery for *R*^*^ = 10. As expected, the most challenging regime is low spatial coverage 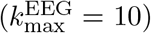 and low SNR, where 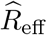 is typically underestimated due to limited information to support higher-rank structure.

The higher-rank *R*^*^ = 15 condition is more challenging under severe missingness, but follows the same monotone pattern: recovery improves substantially as 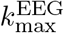 and SNR increase, reaching exact recovery 0.78 and within-one recovery 1.00 in the highest-information setting (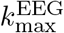, SNR) = (20, 4) (Table 1).

Overall, ARD-based rank adaptivity provides an efficient practical rank-selection strategy, avoiding repeated refitting across candidate ranks while yielding stable estimates in the moderate-to-high information regimes most relevant to EEG–fMRI fusion.

#### 3.3.2. Estimation accuracy in the true-rank setting

We next evaluate estimation performance when the rank is fixed to the true value (*R* = *R*^*^), which we refer to as the *true-rank* (oracle-rank) setting. Figure 4 summarizes recovery of the shared network subspace using the projection-matrix discrepancy 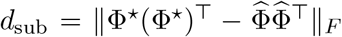 and the mean principal angle between span(Φ^⋆^) and span 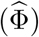 in degrees (∘). Subspace recovery improves systematically with higher SNR and with larger EEG spatial support 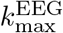, consistent with improved identifiability of the shared Φ as the missing-bandwidth constraints become less severe. Notably, when SNR ≥ 1, the mean principal angles are typically ≤ 5° across scenarios, indicating near-alignment of the estimated and true subspaces on an interpretable angular scale.

**Figure 4:**
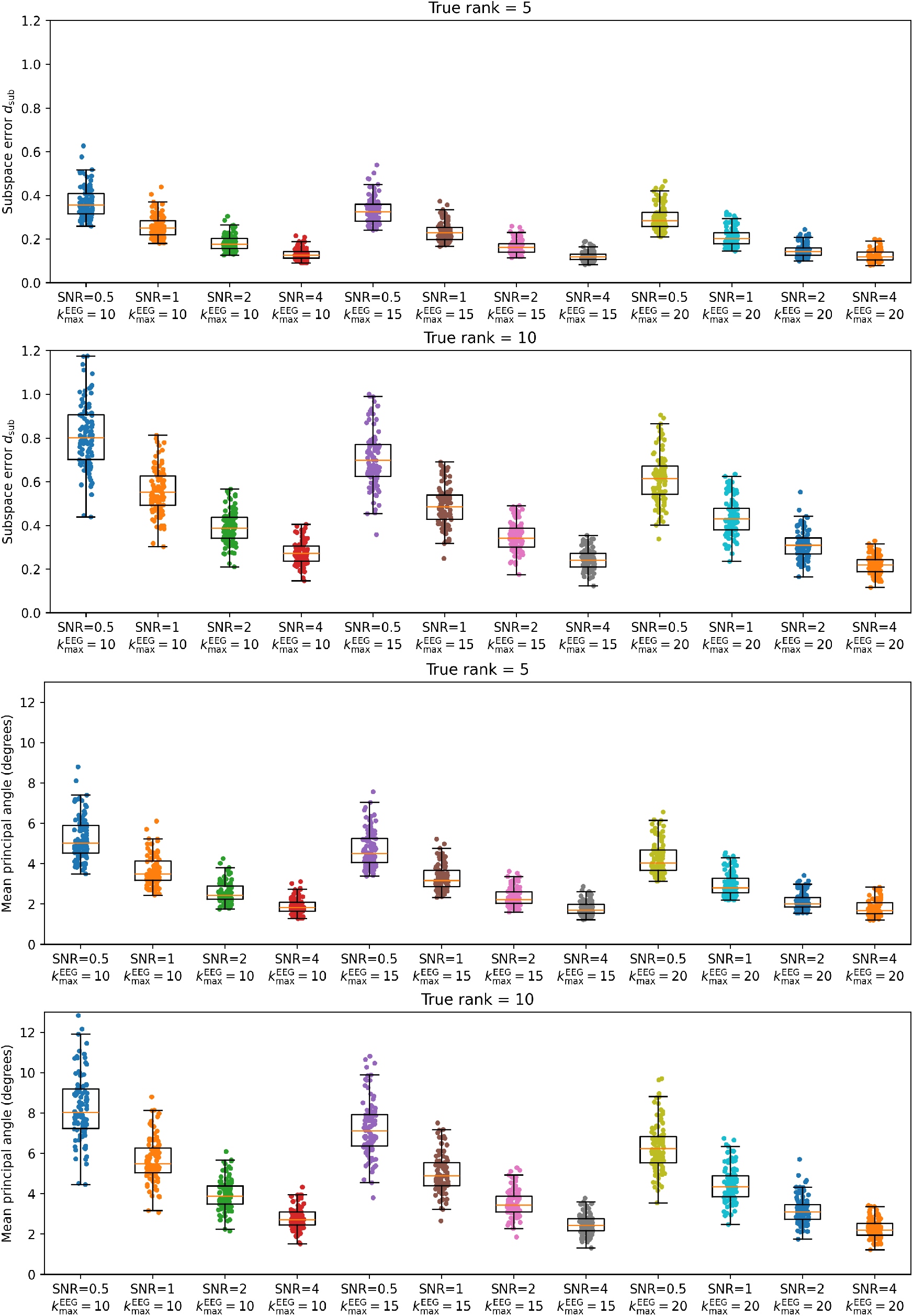
Subspace recovery performance. Top two (1st and 2nd) panels: distribution of *d*_sub_ (projection matrix Φ discrepancy) across 100 replicates by scenario (SNR ∈ {0.5, 1, 2, 4}, 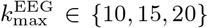) and true rank *R*^*\**^ (*R* = 5 for the first panel and *R*^*\**^ = 10 for the second). Bottom two (3rd and 4th) panels: distribution of the mean principal angle (degrees) between span(Φ^⋆^) and span(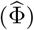) for the same scenarios. Recovery improves with higher SNR and larger EEG spatial support.

Table 2 summarizes spectral-weight recovery (RMSE_*λ*_) and reconstruction accuracy on the observed support (RelErr[ℛ_obs_]) and on the two *structurally unobserved* regions (RelErr[ℛ_EEG-miss_] and RelErr[ℛ_fMRI-miss_]) of the latent connectivity field. Across scenarios, RMSE_*λ*_ decreases monotonically with increasing SNR and with larger EEG spatial support 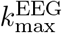, and is larger for the higher-rank setting (*R*^*^ = 10) than for *R*^*^ = 5 at comparable SNR, reflecting the increased degrees of freedom in the spectral-weight profiles. For example, at 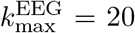 and SNR = 4, RMSE_*λ*_ is 0.041 for *R*^*^ = 5 and 0.055 for *R*^*^ = 10 (Table 2).

**Table 2:** True-rank simulation (MAP fit with *R* = *R*^*\**^): recovery and reconstruction metrics reported as mean (SD) across 100 Monte Carlo replicates per scenario. RelErr[ℛ_obs_] is reconstruction error on observed-support entries; RelErr[ℛ_EEG-miss_] and RelErr[ℛ_fMRI-miss_] quantify *extrapolation* error on structurally unobserved EEG-high-*k* and fMRI-high-frequency regions, respectively.

| True rank $R^*$ | $(k_{\text{max}}^{\text{EEG}}, \text{SNR})$ | $\text{RMSE}_\lambda$ | $\text{RelErr}[\mathcal{R}_{\text{obs}}]$ | $\text{RelErr}[\mathcal{R}_{\text{EEG-miss}}]$ | $\text{RelErr}[\mathcal{R}_{\text{fMRI-miss}}]$ |
| --- | --- | --- | --- | --- | --- |
| 5 | (10, 0.5) | 0.153 (0.018) | 0.192 (0.020) | 0.280 (0.028) | 0.277 (0.027) |
|  | (10, 1) | 0.118 (0.014) | 0.148 (0.017) | 0.209 (0.022) | 0.207 (0.022) |
|  | (10, 2) | 0.092 (0.012) | 0.115 (0.013) | 0.158 (0.017) | 0.156 (0.017) |
|  | (10, 4) | 0.072 (0.010) | 0.091 (0.011) | 0.121 (0.013) | 0.119 (0.013) |
|  | (15, 0.5) | 0.111 (0.014) | 0.147 (0.009) | 0.230 (0.027) | 0.222 (0.026) |
|  | (15, 1) | 0.085 (0.011) | 0.113 (0.008) | 0.170 (0.020) | 0.164 (0.019) |
|  | (15, 2) | 0.066 (0.009) | 0.087 (0.006) | 0.127 (0.015) | 0.122 (0.015) |
|  | (15, 4) | 0.051 (0.008) | 0.072 (0.007) | 0.096 (0.011) | 0.093 (0.011) |
|  | (20, 0.5) | 0.089 (0.009) | 0.122 (0.007) | 0.203 (0.025) | 0.190 (0.023) |
|  | (20, 1) | 0.069 (0.007) | 0.093 (0.005) | 0.149 (0.018) | 0.140 (0.017) |
|  | (20, 2) | 0.053 (0.006) | 0.072 (0.005) | 0.110 (0.014) | 0.104 (0.013) |
|  | (20, 4) | 0.041 (0.006) | 0.069 (0.010) | 0.086 (0.011) | 0.084 (0.011) |
| 10 | (10, 0.5) | 0.215 (0.028) | 0.224 (0.014) | 0.394 (0.045) | 0.385 (0.044) |
|  | (10, 1) | 0.171 (0.023) | 0.173 (0.012) | 0.299 (0.036) | 0.292 (0.036) |
|  | (10, 2) | 0.135 (0.019) | 0.135 (0.011) | 0.226 (0.029) | 0.221 (0.029) |
|  | (10, 4) | 0.105 (0.016) | 0.106 (0.009) | 0.171 (0.024) | 0.167 (0.024) |
|  | (15, 0.5) | 0.148 (0.015) | 0.175 (0.010) | 0.317 (0.037) | 0.300 (0.035) |
|  | (15, 1) | 0.116 (0.010) | 0.134 (0.008) | 0.235 (0.027) | 0.223 (0.026) |
|  | (15, 2) | 0.090 (0.008) | 0.103 (0.007) | 0.175 (0.021) | 0.166 (0.021) |
|  | (15, 4) | 0.070 (0.008) | 0.080 (0.006) | 0.131 (0.017) | 0.124 (0.017) |
|  | (20, 0.5) | 0.115 (0.010) | 0.145 (0.009) | 0.278 (0.036) | 0.252 (0.033) |
|  | (20, 1) | 0.090 (0.008) | 0.112 (0.007) | 0.206 (0.027) | 0.187 (0.025) |
|  | (20, 2) | 0.070 (0.007) | 0.086 (0.006) | 0.154 (0.022) | 0.140 (0.021) |
|  | (20, 4) | 0.055 (0.007) | 0.067 (0.006) | 0.114 (0.017) | 0.104 (0.016) |

Observed-support reconstruction error RelErr[ℛ_obs_] decreases with both SNR and 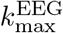; for example, at 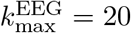 and SNR = 4 we obtain RelErr[ℛ_obs_] = 0.069 for *R*^*^ = 5 and 0.067 for *R* = 10, indicating accurate reconstruction of the latent connectivity on the modality-observed support when information is sufficient (moderate-to-high SNR and non-restrictive EEG spatial coverage). In the same high-information regime (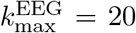, SNR = 4), extrapolation errors on structurally unobserved regions are also small and comparable in magnitude to observed-support error: for *R*^*^ = 5, RelErr[ℛ_EEG-miss_] = 0.086 and RelErr[ℛ_fMRI-miss_] = 0.084, and for *R* = 10 the corresponding errors are 0.114 and 0.104 (Table 2), indicating that the shared low-rank structure can meaningfully extrapolate into structurally unobserved “missing bandwidth” regions rather than merely fitting the observed entries. Extrapolation errors show the same monotone improvements with increasing SNR and EEG spatial support 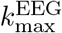, supporting that the shared low-rank structure enables principled inference in missing bandwidths, particularly when the EEG spatial cutoff is not too restrictive and noise is moderate.

## 4. Application to MPI–LEMON

### 4.1. Data and LB-domain connectivity objects

We demonstrate the proposed fusion model using the publicly available MPI–LEMON cohort. We used the eyes-open resting condition for EEG to align with the rs-fMRI instruction (eyes open with fixation). After requiring both usable rs-fMRI and rs-EEG, the primary fusion sample comprised *n* = 189 participants spanning young and older adults (Supplementary Methods: sample definition in Section S3; preprocessing details in Sections S4–S5).

All analyses are performed in a common cortical eigenmode coordinate system (see Section 2.1 and Supp. Section S2). For each subject *i*, the model takes as inputs: (i) an rs-fMRI connectivity matrix 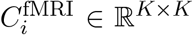 computed from LB-projected BOLD coefficients over low temporal frequencies, and (ii) frequency-resolved EEG connectivity matrices 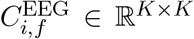 over a discrete grid 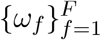 spanning [1, 45] Hz, computed from LB-domain EEG coefficients using leakage-robust envelope covariance. EEG connectivity is treated as informative primarily on coarse spatial modes (low *k*), which we encode using the fixed mask *W* ^EEG^ with 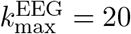 and *K* = 50 (Supplementary Methods Section S1).

### 4.2. Real-data model specification and selection workflow

#### Cross-modality scaling

Because raw EEG and fMRI connectivity magnitudes can differ substantially, we rescale each modality by a robust global factor prior to fitting baed on the median Frobenius norms (Supplementary Methods Section S6).

#### Rank selection and energy screening

We selected the working rank *R* using a two-stage procedure. First, we fit an over-parameterized model with *R*_max_ = 20 under ARD-style regularization and computed an effective rank 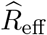 by thresholding the factor-energy curve at *E*_*r*_*/E*_max_ ≥ 0.01, yielding 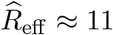. Second, we constructed elbow plots of reconstruction error across candidate ranks *R* ∈ {1, 3, 5, 8, 9, 10, 11, 12, 13, 14, 15}, summarizing (i) median subject-level relative error for fMRI connectivity, (ii) median subject-level relative error for EEG connectivity, and (iii) a combined diagnostic for the fMRI and EEG relative errors (Supplementary Fig. S1; Section S6). Based on the elbow behavior across these panels, we selected *R* = 12 for the main analysis, consistent with 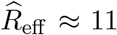 up to ±1. Finally, for interpretability and downstream analyses we rank networks by decreasing median total factor energy and use this ordering for Top-3,-5,-7, and -10 energy screening from the fitted rank-12 model to demonstrate robustness to using smaller, high-energy subsets.

### 4.3. Synergy of fused EEG–fMRI LB-network features for age prediction

To quantify the practical value of multimodal fusion in MPI–LEMON, we tested whether subject-level LB-network features from the fused model improve out-of-sample prediction of the participants’ age (midpoint of the recorded age bin) beyond what is achievable from either modality alone.

#### Fused features and energy screening

From the fitted fused model (*R* = 12), each subject *i* is summarized by (i) an fMRI network-strength vector 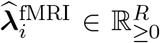 (one scalar per network; *R* = 12) and (ii) the corresponding EEG-derived spectral summaries per network computed from the fitted nonnegative weights 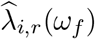: a *spectral centroid* 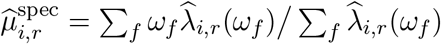 and *band-mass summaries* in *θ/α/β/γ* (bin-sums of 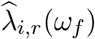 within each frequency band). To reduce dimensionality and assess stability with respect to network selection, we also perform energy-based screening. After fitting the rank-*R* = 12 model, networks are ranked by median total factor energy, and downstream analyses are repeated using only the Top-*K* networks, with *K* ∈ {3, 5, 7, 10, 12}.

#### Nested-CV prediction and synergy metrics

We performed nested cross-validation (CV) with five outer folds using ridge regression, selecting the regularization parameter within each training fold by an inner holdout split. All models adjust for sex (male). We compare covariates only (*M*_0_), covariates+fMRI features (*M*_fMRI_), covariates+EEG features (*M*_EEG_), and covariates+fMRI–EEG fused features (*M*_fused_). The fMRI–EEG synergy is quantified by the incremental gain of adding one modality given the other modality:

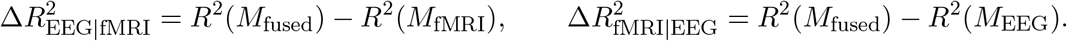

#### Concatenation baseline (feature-level early fusion)

We also evaluate a concatenation model *M*_concat_ that stacks modality-specific features extracted from separate single-modality low-rank fits (fit independently for EEG-only *M*_EEG_ and fMRI-only *M*_fMRI_) as an early-fusion baseline, and then applies the same nested-CV ridge protocol.

#### Results and interpretation

Table 3 shows that using all 12 networks (“All (12)”), the fused fMRI–EEG model achieved substantial out-of-sample performance for age prediction (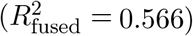), clearly exceeding the single-modality models (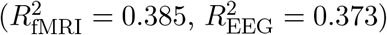) and yielding positive synergy in both directions 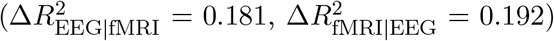. Notably, restricting to the Top-10 and Top-7 networks preserved (and slightly improved) predictive performance, suggesting that the age-relevant information is concentrated in a subset of high-energy networks. Even a compact representation using only the Top-5 networks maintained strong out-of-sample prediction 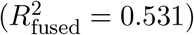, whereas Top-3 networks did not fully capture the signal. The positive incremental gains 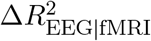 and 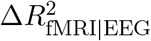 for *M*_fused_ indicate that the fused representation leverages complementary information: frequency-resolved spectral summaries add value beyond spatial strength alone, and conversely spatial strength adds value beyond spectral summaries alone. Relative to the fusion model, the concatenation baseline (*M*_concat_) performs competitively, which is expected because it uses the same low-dimensional feature types; the key difference is that these features are extracted from separate unimodal fits (*M*_EEG_ and *M*_fMRI_) rather than from a jointly learned network basis. The fused model *M*_fused_ is nevertheless comparable to or modestly better than concatenation across settings (notably for Top-7), and it yields larger relative improvements in more parsimonious regimes (e.g., Top-3), supporting the value of learning a shared network basis. Crucially, the fused model preserves a *single shared* network coordinate system that supports the joint interpretability analyses below (surface maps, spectral profiles, and the scale–frequency signature; Figures 6, 7, and 9).

**Table 3:** Nested-CV synergy analysis for predicting age from the fused network features in MPI–LEMON, adjusting for sex (male indicator). We report mean (SD) out-of-sample *R*^2^ across 5 outer folds. “Top-*K* networks” retains the *K* highest-energy networks (ranked by median total factor energy), yielding *K* fMRI-strength features and 5*K* EEG-derived features (one spectral centroid and four band-mass summaries per retained network). We report the coefficient of determination *R*^2^ for the fused model, and for single-modality models (fMRI-only and EEG-only). Synergy is quantified by 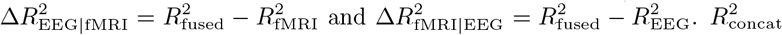 denotes an early-fusion baseline obtained by concatenating unimodal features from separate unimodal fits.

| Top-$K$ | $R_{\text{fused}}^2$ (SD) | $R_{\text{fMRI}}^2$ (SD) | $R_{\text{EEG}}^2$ (SD) | $R_{\text{concat}}^2$ (SD) | $\Delta R_{\text{EEG\vert fMRI}}^2$ | $\Delta R_{\text{fMRI\vert EEG}}^2$ |
| --- | --- | --- | --- | --- | --- | --- |
| All (12) | 0.566 (0.129) | 0.385 (0.109) | 0.373 (0.161) | 0.546 (0.171) | 0.181 | 0.192 |
| Top-10 | 0.567 (0.130) | 0.383 (0.114) | 0.388 (0.161) | 0.566 (0.111) | 0.184 | 0.179 |
| Top-7 | 0.582 (0.106) | 0.349 (0.085) | 0.442 (0.101) | 0.529 (0.098) | 0.233 | 0.140 |
| Top-5 | 0.532 (0.082) | 0.259 (0.101) | 0.441 (0.082) | 0.509 (0.148) | 0.273 | 0.091 |
| Top-3 | 0.340 (0.138) | 0.203 (0.112) | 0.234 (0.110) | 0.289 (0.119) | 0.136 | 0.106 |

### 4.4. Results II: Age associations with fused network features

We next examine which fused features drive the age-prediction signal by testing their marginal associations with age:

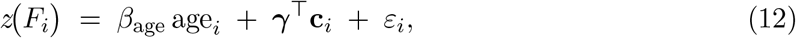

where *F*_*i*_ denotes a subject-level feature (network strength, spectral centroid, and *θ/α/β/γ* band-mass summaries; Section 4.3), *z*(·) denotes across-subject standardization (z-scoring), and **c**_*i*_ denotes covariates (primary: sex; sensitivity: sex+education). We fit (12) by ordinary least squares and report heteroskedasticity-consistent (HC) standard errors using the HC3 estimator (MacKinnon and White, 1985; Long and Ervin, 2000). Multiple testing across features was controlled using Benjamini–Hochberg false discovery rate (BH-FDR) across all tested features, and we report *q*-values. For interpretability, we report effects per decade (10-year change).

We focus primary interpretation on features derived from the Top-5 energy-ranked networks. For completeness, Supplementary Results (Section S8.2) report (i) the full fused-feature association analysis results for all *R*=12 networks, (ii) the single-modality association analysis results using fMRI-only strength features and EEG-only spectral-summary features extracted from unimodal fits, and (iii) a sensitivity analysis that additionally adjusts for education (sex+education), which yields qualitatively similar conclusions.

#### Primary association results and interpretation

Table 4 summarizes age associations for fused features from the Top-5 networks, reporting per-decade standardized effects *β*_age_, standard errors (SE), and BH-FDR *q*-values. Across Networks 1–5, fMRI-derived network strengths 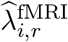 show consistently negative age associations, while network spectral centroids 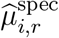 (see Section 4.3 for definition) show consistently positive age associations. Among EEG band-mass summaries, *θ* and *α* band masses tend to decrease with age across multiple networks, whereas *β* band mass effects are weaker and more variable; in contrast, *γ* band mass shows a consistent positive association with age across the Top-5 networks. Figure 5 visualizes the strongest associations by absolute effect magnitude, grouped by feature type. Together, these results show coordinated age dependence within the same fused components: fMRI network strength tends to decrease with age, while EEG summaries shift toward higher effective frequency (higher spectral centroid, increased *γ* mass, and reduced *θ/α* mass), all expressed in a single shared network coordinate system.

**Table 4:** Age associations for fused features derived from the Top-5 energy-screened networks, adjusting for sex. Each row corresponds to a univariate regression of a *z*-scored feature on age and sex, using robust standard errors with BH-FDR correction across tests. Effects are reported per decade (10-year change) in SD units of the feature. Rows are ordered by network ID (energy order) and feature type.

| Network | Feature summary | $\beta$ (per decade) | SE | $q$ |
| --- | --- | --- | --- | --- |
| Network 1 | Network strength ($\lambda^{\text{fMRI}}$) | -0.156 | 0.035 | 1.90e-05 |
| | Spectral centroid ($\mu^{\text{spec}}$) | 0.187 | 0.043 | 3.45e-05 |
| | Band mass: $\theta$ | -0.126 | 0.033 | 2.48e-04 |
| | Band mass: $\alpha$ | -0.124 | 0.032 | 2.22e-04 |
| | Band mass: $\beta$ | -0.074 | 0.033 | 3.12e-02 |
| | Band mass: $\gamma$ | 0.150 | 0.045 | 1.42e-03 |
| Network 2 | Network strength ($\lambda^{\text{fMRI}}$) | -0.139 | 0.040 | 9.40e-04 |
| | Spectral centroid ($\mu^{\text{spec}}$) | 0.272 | 0.046 | 1.63e-08 |
| | Band mass: $\theta$ | -0.155 | 0.031 | 2.29e-06 |
| | Band mass: $\alpha$ | -0.112 | 0.033 | 1.08e-03 |
| | Band mass: $\beta$ | -0.032 | 0.035 | 4.03e-01 |
| | Band mass: $\gamma$ | 0.206 | 0.042 | 2.29e-06 |
| Network 3 | Network strength ($\lambda^{\text{fMRI}}$) | -0.244 | 0.032 | 4.52e-13 |
| | Spectral centroid ($\mu^{\text{spec}}$) | 0.241 | 0.046 | 6.26e-07 |
| | Band mass: $\theta$ | -0.140 | 0.032 | 3.72e-05 |
| | Band mass: $\alpha$ | -0.110 | 0.033 | 1.42e-03 |
| | Band mass: $\beta$ | -0.041 | 0.036 | 2.96e-01 |
| | Band mass: $\gamma$ | 0.164 | 0.050 | 1.44e-03 |
| Network 4 | Network strength ($\lambda^{\text{fMRI}}$) | -0.239 | 0.034 | 1.36e-11 |
| | Spectral centroid ($\mu^{\text{spec}}$) | 0.288 | 0.039 | 1.46e-12 |
| | Band mass: $\theta$ | -0.092 | 0.036 | 1.35e-02 |
| | Band mass: $\alpha$ | -0.056 | 0.037 | 1.54e-01 |
| | Band mass: $\beta$ | 0.030 | 0.039 | 4.80e-01 |
| | Band mass: $\gamma$ | 0.198 | 0.043 | 1.10e-05 |
| Network 5 | Network strength ($\lambda^{\text{fMRI}}$) | -0.132 | 0.036 | 3.87e-04 |
| | Spectral centroid ($\mu^{\text{spec}}$) | 0.296 | 0.034 | 2.41e-17 |
| | Band mass: $\theta$ | -0.137 | 0.033 | 6.92e-05 |
| | Band mass: $\alpha$ | -0.095 | 0.034 | 7.65e-03 |
| | Band mass: $\beta$ | 0.012 | 0.037 | 7.78e-01 |
| | Band mass: $\gamma$ | 0.211 | 0.044 | 5.02e-06 |

**Figure 5:**
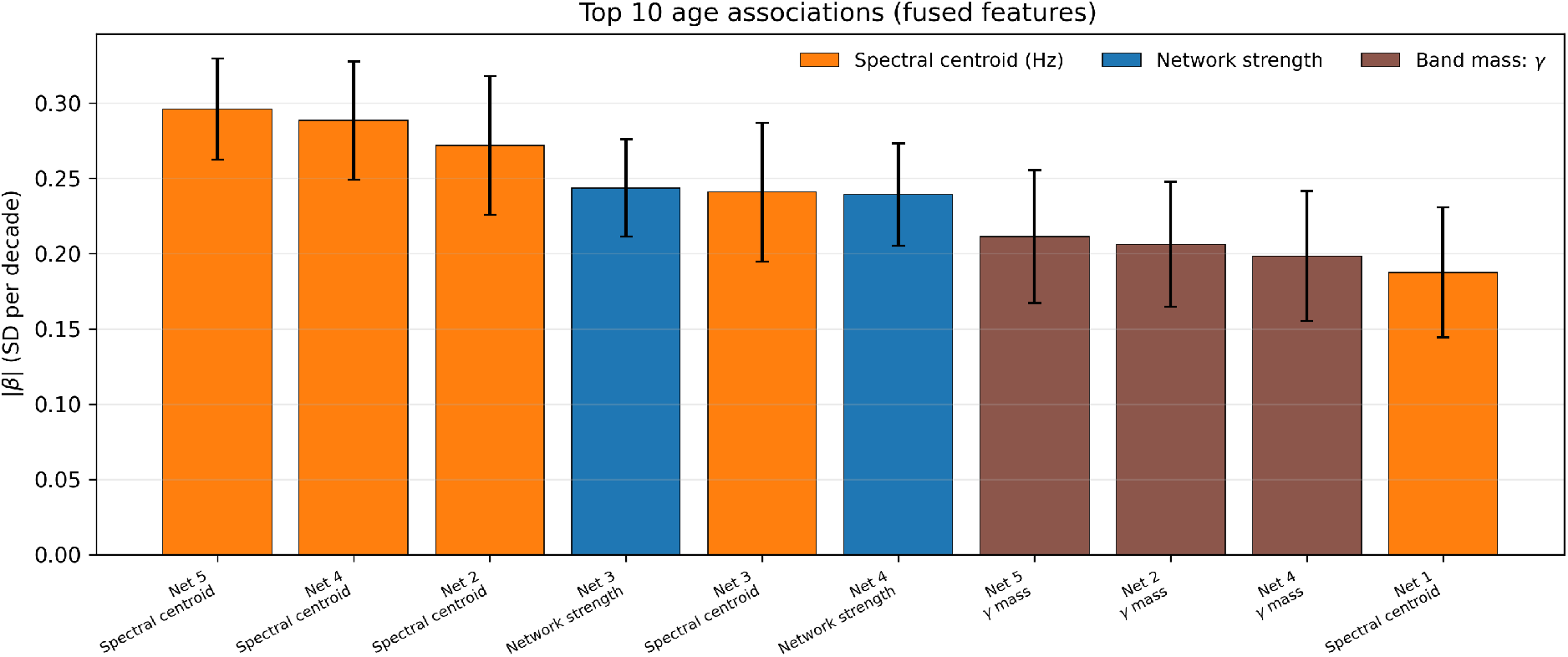
MPI–LEMON: top-10 age associations among fused features from the Top-*K* energy-screened networks (*K* = 5), adjusting for sex. Bars show the absolute standardized effect of age on each feature (SD change in the feature per decade), estimated from univariate regressions of the form *z*(*F*_*i*_) = *β*_age_age_*i*_ + *γ* male_*i*_ + *ε*_*i*_ with HC3 standard errors; error bars denote the corresponding robust SEs (per decade). Features are derived from the shared fused network coordinates and include network spatial strength (fMRI-derived) and EEG spectral summaries (spectral mean in Hz and band-mass summaries in *θ/α/β/γ*). Color indicates feature type.

### 4.5. Results III: Network maps and spectral summaries for the top networks

#### Surface maps and system-level composition

Figure 6 shows the Top-5 fused networks (ordered by median total factor energy) on the `fsaverage5` surface (lateral/medial views; both hemispheres). The highest-energy components exhibit structured, spatially smooth patterns spanning association, sensory, and attention-related territories, broadly consistent with canonical resting-state organization (Yeo et al., 2011; Schaefer et al., 2018; Babayan et al., 2019). Surface maps for networks *r* = 6, …, 12 are provided in Supplementary Results (Fig. S8, Section S8.3).

**Figure 6:**
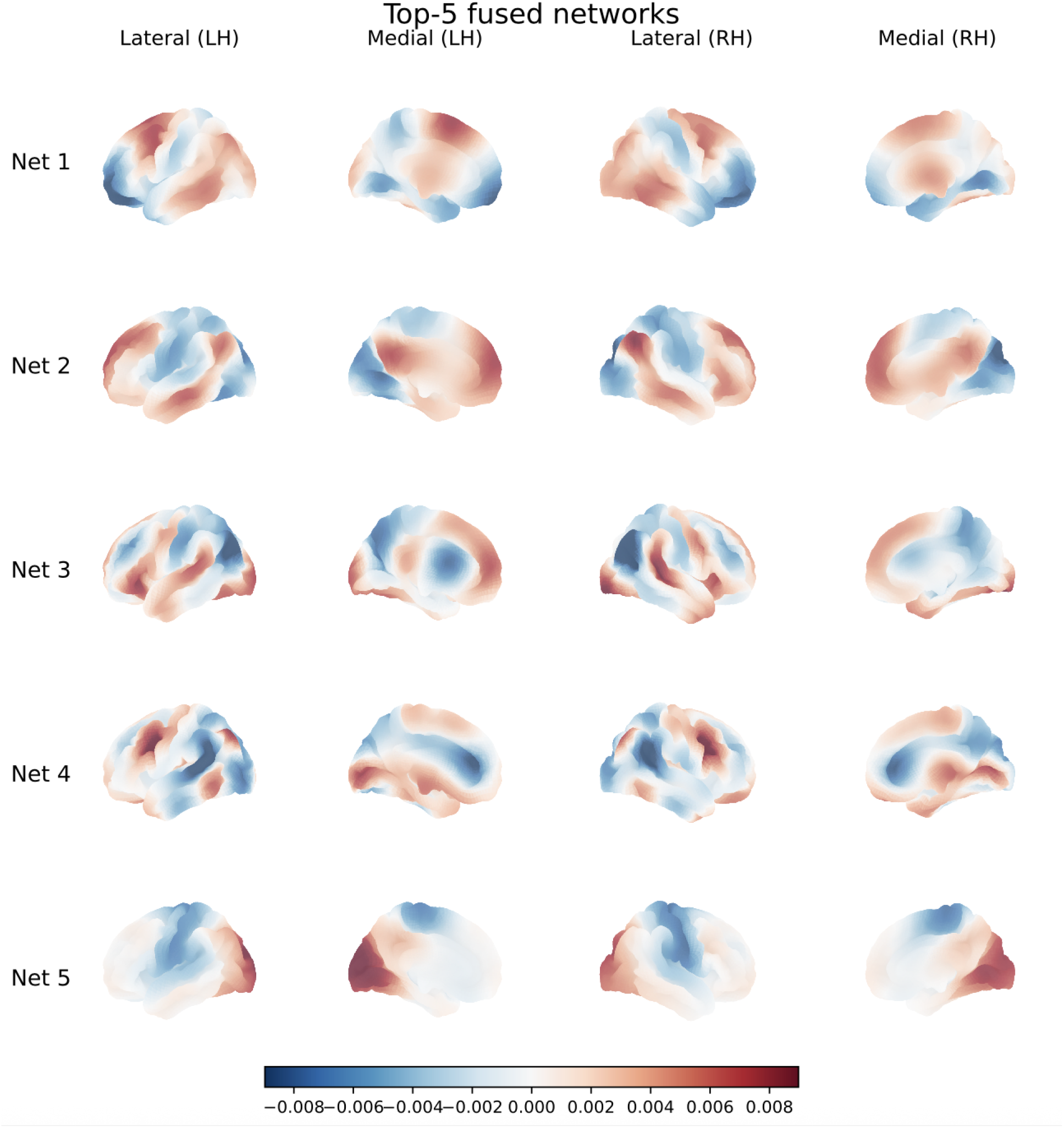
MPI–LEMON: surface visualization of the Top-5 fused spatial networks (rank-*R* = 12), displayed on `fsaverage5`. Each row shows one factor (Net *r* = 1, …, 5; ordered by median factor energy), with four columns corresponding to left-hemisphere (LH) lateral/medial views and right-hemisphere (RH) lateral/medial views.

Several networks show clear qualitative correspondence to familiar systems based on their dominant anatomical drivers (e.g., Net 5 is early-visual dominant; Net 8 shows a salience/cinguloopercular motif with frontal operculum/insula and cingulate contributions; Net 10 resembles a dorsal-attention pattern with frontal eye fields and superior parietal involvement; Net 11 is somatomotor-dominant; Table 5). Other networks capture cross-system structure expressed as smooth transitions along the macroscale functional hierarchy. Such hybrid components are expected in low-dimensional representations of resting-state organization and are compatible with gradient-like descriptions of macroscale cortical connectivity (Margulies et al., 2016; Huntenburg et al., 2018; Vos de Wael et al., 2020; Bernhardt et al., 2022). For example, Nets 1–4 are Default-enriched but show substantial secondary contributions from Control, Visual, and/or Dorsal-Attention territories.

**Table 5:** Canonical identities, Top-2 Schaefer-7 systems (percent mass), and primary anatomical drivers for the 12 energy-ordered fused networks. Visual identities are based on the signed surface loading maps (after the network-wise sign convention in Fig. 6) and complement the ROI/system summaries in Supplementary Table S2.

| Net $r$ | Identity (visual) | Top-2 systems (% mass) | Primary anatomical drivers (high-loading regions) |
| --- | --- | --- | --- |
| 1 | Default–Control hybrid | Default (22.6), Cont (16.7) | dlPFC/lateral PFC; OFC; temporal pole; angular/lateral parietal |
| 2 | Default–Visual hybrid | Default (28.6), Vis (19.8) | pCun/PCC; lateral parietal; occipital/visual cortex |
| 3 | Default–Dorsal Attention hybrid | Default (27.8), Vis (15.4) | posterior parietal/IPS; pCun/PCC; FEF patches |
| 4 | Default–Control gradient | Default (24.2), Vis (17.8) | lateral PFC; parietal association cortex; posterior midline |
| 5 | Visual-dominant (early) | SomMot (33.8), Vis (29.4) | calcarine sulcus; occipital pole; cuneus |
| 6 | Visual–Somatomotor hybrid | SomMot (23.2), Default (23.1) | occipital/visual; pre/postcentral (somatomotor) strip |
| 7 | Frontoparietal / Executive | Default (22.4), Cont (19.1) | dlPFC/lateral PFC; posterior parietal cortex |
| 8 | Salience / Cingulo-opercular | SomMot (20.5), Vis (19.2) | insula; frontal operculum; mid/anterior cingulate |
| 9 | Temporal–Limbic / DMN | Default (28.4), SomMot (19.8) | lateral temporal cortex; temporal pole |
| 10 | Dorsal Attention (DAN) | Default (22.2), SomMot (18.2) | FEF; superior parietal lobule |
| 11 | Somatomotor-dominant | SomMot (28.9), Default (21.6) | bilateral precentral and postcentral gyri |
| 12 | Limbic / Orbitofrontal | Default (22.2), SomMot (17.7) | OFC; anterior temporal pole |

#### Network identities from signed surface maps

To facilitate interpretation, Table 5 assigns each component a visual identity based on its surface-loading pattern (spatial distribution in Figures 6 and S8). The table also summarizes (i) the Top-2 Schaefer-7 systems by magnitude-based percent (%) mass (computed as in Supplementary Methods Section S7) and (ii) primary anatomical drivers (high-loading regions).

#### ROI-based system composition of fused networks

To complement the visual identities and anatomical drivers in Table 5, we quantified each network’s full system composition with respect to the Schaefer2018 parcellation (200 parcels; 7-network labeling) on `fsaverage5` using magnitude-based loadings (% parcel mass; Supplementary Methods Section S7). Because these compositions are magnitude-based (absolute loadings), they summarize overall mass within systems rather than the signed opposition patterns visible in the surface maps. Supplementary Figure S5 and Supplementary Table S2 show that most networks have a dominant system together with nontrivial secondary contributions. Among the Top-5 networks, Nets 1–4 are Default-enriched (Default ≈ 23–29% by mass) with substantial secondary Control and Visual contributions (e.g., Net 1: Default 22.6%, Cont 16.7%; Net 4: Default 24.2%, Vis 17.8%). Net 5 illustrates how a visually salient driver can coexist with broader mixing: although its signed surface map is early-visual dominant, its magnitude-based atlas composition also includes substantial SomMot mass in addition to Vis (Table 5).

#### Spectral profiles and subject-level summaries

To summarize each network’s oscillatory signature, Figure 7 (left) plots the distribution of subject-specific spectral weight curves 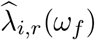 across the modeling frequency grid. Across networks, the median profiles generally decrease with frequency (consistent with an aperiodic 1*/f* -like background), but differ in their relative low-vs high-frequency emphasis, yielding network-specific spectral summaries (Donoghue et al., 2020). Figure 7 (right) summarizes inter-individual variability via compact subject-level scores: 1) the fMRI-derived network strength; and 2) EEG-derived spectral summaries (spectral centroid; Section 4.4). Together, these summaries provide a readout of (i) how strongly each spatial network is expressed (through 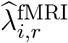) and (ii) how its expression is distributed across oscillatory frequencies (through 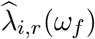, summarized via 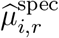 and band mass summaries).

**Figure 7:**
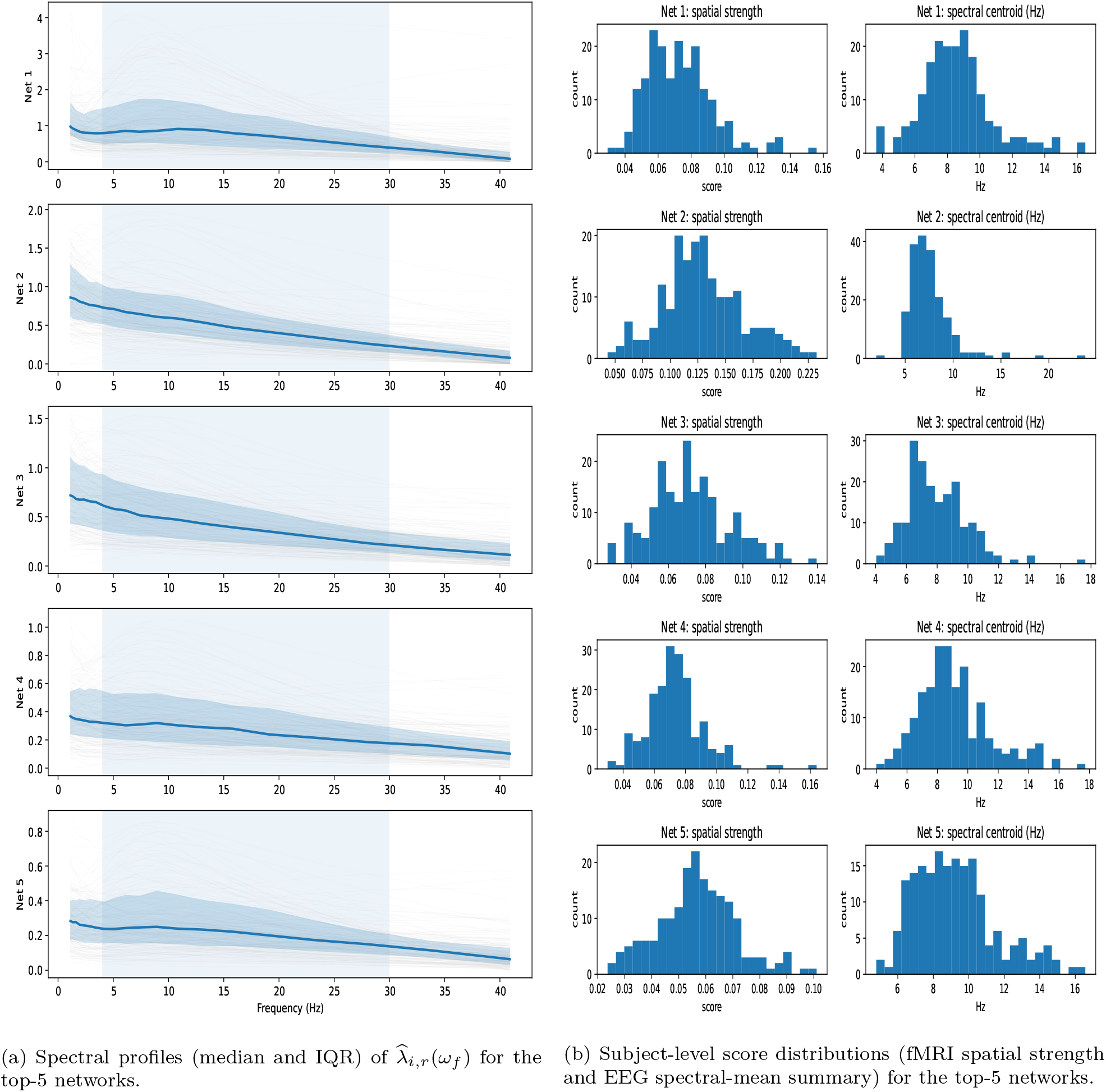
MPI–LEMON: spectral and subject-level summaries for the fused networks (Top-5; rank-*R* = 12). **Left:** spectral profiles of subject-specific EEG weights 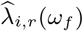 (light gray curves) across the modeling frequency grid, and the median across subjects with an interquartile range (25th–75th percentiles) band. **Right:** histograms of subject-level network scores, including the fMRI-derived spatial-strength score 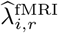 (one scalar per subject and network) and the EEG-derived spectral-mean summary 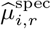 (Hz-weighted center of mass of 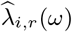).

Analogous spectral/score summaries and age association plots for networks *r* = 6, …, 12 are provided in Supplementary Results (Figs. S9 and S10). Figure 8 provides a visual complement to the association screen (Section 4.4) by plotting age versus (i) spatial network strength and (ii) spectral centroid for the Top-5 networks. These plots visualize the direction and variability underlying the regression results in Table 4.

**Figure 8:**
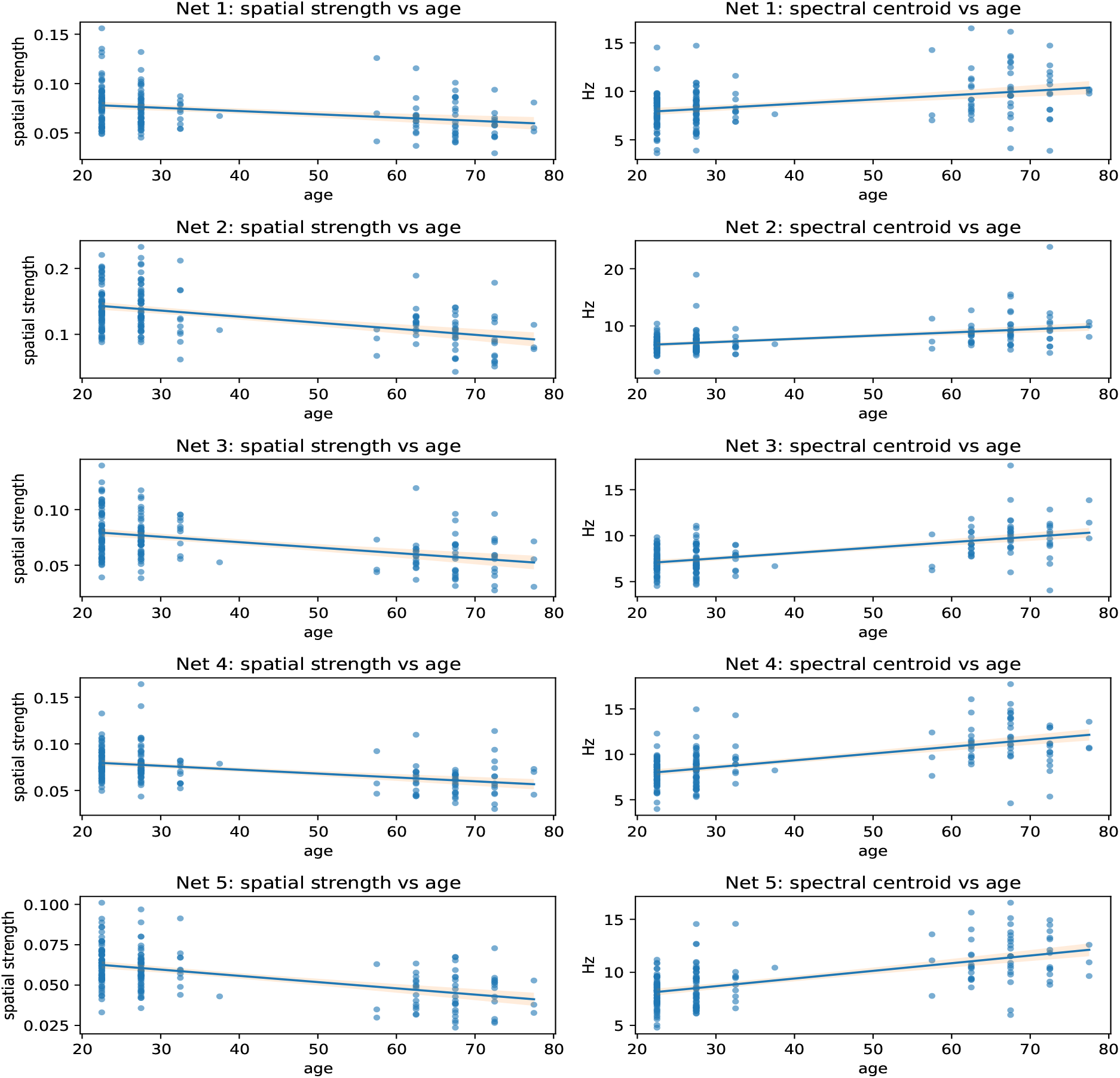
MPI–LEMON: age relationships for Top-5 fused network summaries (rank-*R* = 12). For each network, we show scatter plots of subject age versus (left) fMRI-derived spatial-strength score 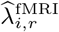; and (right) EEG-derived spectral-mean summary 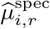. Overlaid lines denote linear regression fits with confidence bands.

#### Anatomical distribution of age-related variation

Age associations in the fused feature set are summarized in Table 4. Across the Top-5 energy-screened networks, the fMRI-derived network-strength features 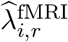 show consistently negative age associations, with the largest effects in Net 3 (*β* = −0.244 per decade) and Net 4 (*β* = −0.239 per decade). Based on the signed surface-map identities (Table 5) and atlas-referenced compositions (Figure S5, Table S2), the strongest effects concentrate in Default-/association-enriched components with systematic contributions from control/attention and posterior visual–parietal territories, consistent with prior rs-fMRI reports that association-system connectivity (including default-mode and control-related circuits) is particularly sensitive to healthy aging (Babayan et al., 2019; Hedden et al., 2009; Staffaroni et al., 2018). These spatial patterns are also compatible with gradient-based accounts of adult lifespan reorganization that describe age-related redistribution of functional communities in low-dimensional connectivity space (Bethlehem et al., 2020).

On the spectral side, we observe systematic age dependence within the same components: spectral centroids tend to increase across the Top-5 networks, while low-frequency band-mass summaries (especially *θ* and *α*) show predominantly negative associations in several networks (Table 4). These trends are compatible with reported age-related changes in resting EEG, including reduced alpha-band activity and shifts in spectral organization that can reflect both oscillatory and aperiodic components (Scally et al., 2018; Merkin et al., 2023; Donoghue et al., 2020).

#### Interpretability payoff of a shared network coordinate system

Together, the surface maps (Figure 6), atlas-referenced compositions (Table 5; Supplementary Table S2), and feature-level age associations (Figure 7, Table 4) provide a coherent multi-view interpretation: each fused component is anchored to (i) a cortical topography, (ii) a standard system-level reference profile, and (iii) coupled spatial-strength and oscillatory summaries that exhibit systematic age dependence. This unified readout is difficult to obtain from parallel unimodal analyses and clarifies, within the same network coordinates, both *where* (cortex) and *how* (oscillatory distribution) age-related variability manifests.

### 4.6. Results IV: Scale–frequency organization across networks

An advantage of the cortical eigenmode representation is that it provides an explicit *spatial-frequency ordering* through the LB eigenvalues. To quantify the spatial scale of each learned network *r*, we compute an effective spatial-frequency index by eigenvalue-weighting the squared LB-factor coefficients:

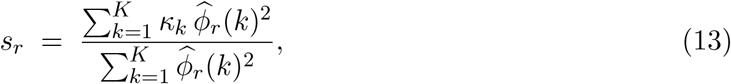

where 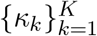 are the cortical template LB eigenvalues (in increasing order) and 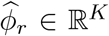 is the estimated LB-domain spatial factor for network *r*. Larger *s*_*r*_ indicates relatively greater weight on higher spatial-frequency modes (finer-scale structure). We summarize each network’s oscillatory signature by the across-subject median spectral centroid, 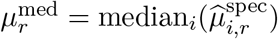.

Figure 9 plots 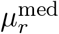 against *s*_*r*_ for all *R* = 12 networks. Most networks (except Nets 3–4) follow an overall positive scale–frequency trend, consistent with systematic organization linking finer spatial structure (larger *s*_*r*_) to higher-frequency oscillatory expression in the fused representation. Nets 3–4 deviate from this pattern by combining relatively high *s*_*r*_ with comparatively low spectral centroids. Both are Default-enriched cross-system components (Table 5), compatable with the possibility that slower oscillatory signatures can accompany finer-scale spatial structure in association-dominant patterns.

**Figure 9:**
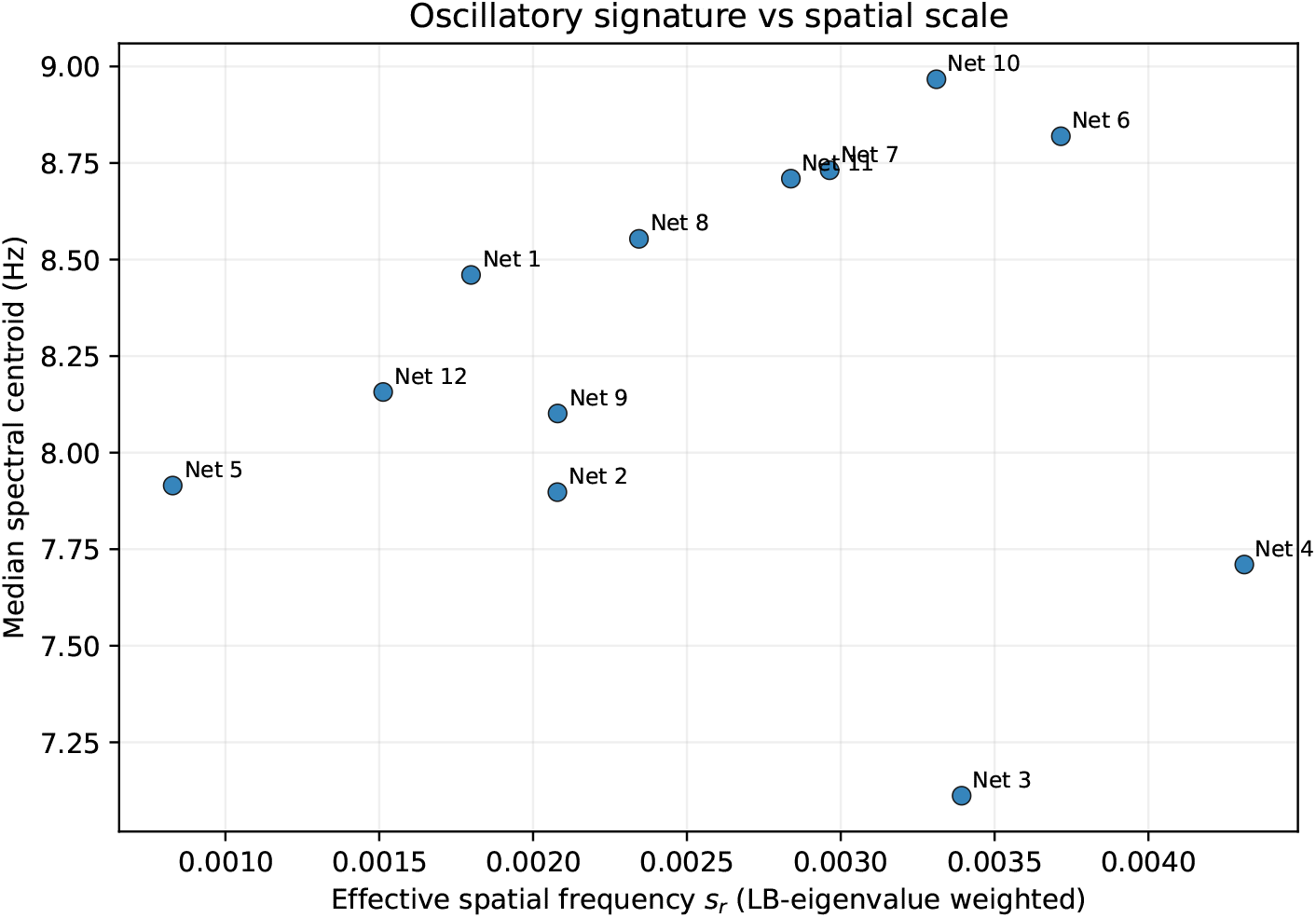
MPI–LEMON: oscillatory signature as a function of effective spatial frequency across fused networks (*R* = 12). Each point represents one network *r* learned by the fused LB EEG–fMRI model. The x-axis shows the network’s effective spatial frequency *s*_*r*_, computed from the LB eigenvalues as a weighted summary (Eq. 13) of how strongly the network loads onto higher spatial-frequency modes (higher *s*_*r*_ indicates a finer spatial scale). The y-axis shows the network’s spectral centroid (median across subjects), i.e., the Hz-weighted center of mass of the estimated EEG spectral weights 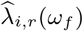. Points are labeled by network index.

#### Representative latent spatio-spectral connectivity field

To visualize the estimated latent object itself, we formed a representative latent field

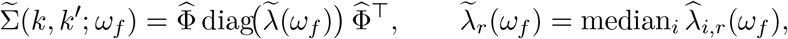

using the median subject-level factor strengths at each frequency. Its diagonal summary, 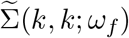, shows that latent variance is concentrated primarily in the first ∼ 10–12 LB modes, with the largest contributions at lower frequencies and a general decay in magnitude as frequency increases in Figure 10 (left). Thus, the fitted field is dominated by coarse, large-scale cortical structure rather than being diffusely distributed across all spatial modes. In Figure 10 (right), representative *K* × *K* slices (at *α* and high *β*) show that the dominant mode-by-mode covariance is concentrated in a low-*k* block (roughly the first ∼ 12 modes, and more weakly through about mode 20), whereas higher modes exhibit weaker and less coherent structure. Across frequencies, the broad coupling pattern is similar, while lower-frequency slices primarily differ by stronger overall magnitude. Together, these summaries show that the estimated latent field is structured rather than diffuse, with variance and coupling concentrated in coarse spatial modes and strongest at lower frequencies.

**Figure 10:**
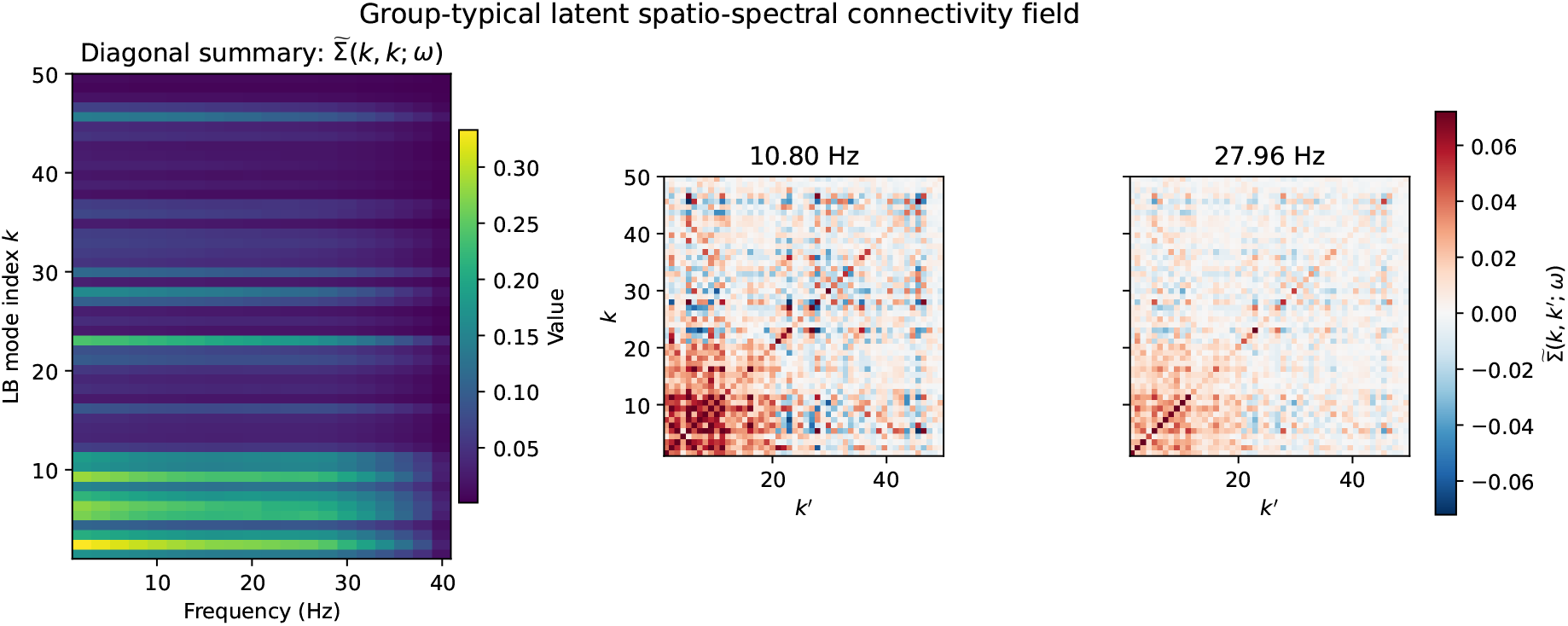
Representative latent spatio-spectral connectivity field in LB mode space. The field is defined as 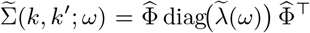, where 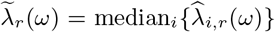 is the median subject-level strength of network *r* at frequency *ω*. **Left:** diagonal summary 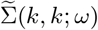 shown as a heatmap over LB mode index *k* and frequency *ω*, indicating how variance is distributed across spatial scale *k* and spectral regime. **Right:** representative full *K* × *K* slices at selected frequencies (10.80 Hz in *α* and 27.96 Hz in high *β*), showing the mode-by-mode coupling structure of the latent field. The field is dominated by low-*k* structure and is strongest at lower frequencies. (The left (diagonal summary) and right panels use separate color scales to preserve visibility of off-diagonal structure.)

#### Coherence across results

Taken together, the synergy analysis (Table 3), feature-level age associations (Table 4), and the scale–frequency summary (Figure 9) support a consistent interpretation: the fused LB-network representation yields compact subject-level features that capture complementary spatial (fMRI) and oscillatory (EEG) organization relevant to aging, while remaining interpretable via cortical surface maps and spectral summaries.

## 5. Discussion

We presented a framework for fusing resting-state EEG and fMRI connectivity in a shared cortical eigenmode coordinate system. By modeling a latent LB-domain spatio-spectral connectivity field with a PSD low-rank factorization, the method yields a compact set of spatial networks together with subject-specific oscillatory strength profiles. A key design choice is to treat modality-specific unobserved bandwidths as *structurally missing* and to infer them through shared low-rank structure and spectral regularization, rather than estimating high-dimensional connectivity objects independently within each modality. This produces interpretable subject-level features that leverage the complementary spatial specificity of fMRI and the spectral resolution of EEG while respecting modality-specific bandwidth limits.

### Main empirical findings in MPI–LEMON

In the MPI–LEMON application, fused LB-network features supported strong out-of-sample prediction of age under nested cross-validation, with consistent incremental gains when combining EEG and fMRI features relative to either modality alone (Table 3). Predictive performance remained substantial when restricting to a small subset of energy-screened networks (e.g., Top-5), suggesting that age-relevant information is concentrated in a compact set of high-energy factors. Complementing prediction, feature-on-age association screening with robust standard errors and BH-FDR control identified systematic age-related shifts in both fMRI strength and EEG spectral summaries, providing a characterization of how the fused representation varies across the adult lifespan.

### Aging signatures across fused spatial and spectral summaries

Age-related variation was most evident in high-energy, Default-enriched components that also incorporated attention/control and posterior visual–parietal territories (e.g., Nets 3–4; Tables 5). From a connectome-gradient perspective, such cross-system components are compatible with age-sensitive shifts in the embedding of functional communities reported in lifespan analyses of gradient dispersion (Bethlehem et al., 2020). In line with prior rs-fMRI work, prominent age associations in network strength *λ*^fMRI^ occurred in Default-enriched components with substantial secondary control/attention and posterior visual–parietal contributions (Table 4), echoing reports that association networks and default–control/attention balance are especially age sensitive (Hedden et al., 2009; Staffaroni et al., 2018). In the fused representation, these components are informative because the same network factor indexes coordinated variation in spatial expression (fMRI strength) and frequency-domain organization (EEG summaries) within a shared coordinate system. Within these same components, EEG-derived summaries also showed coordinated age dependence: spectral centroid increased with age across all Top-5 networks, *θ* and *α* band-mass summaries were predominantly negative (with some network-to-network heterogeneity), and *γ* mass effect was consistently positive, whereas *β* effects were weaker and more variable (Table 4). These patterns mirror well-replicated resting EEG aging signatures, including reduced alpha activity and age-related changes in aperiodic structure that can shift relative spectral weight toward higher frequencies (Scally et al., 2018; Donoghue et al., 2020; Merkin et al., 2023).

### Multi-view interpretation of learned networks

A central motivation for the eigenmode formulation is interpretability. Each learned component can be read out as (i) a cortical surface map (Figure 6), (ii) an atlas-referenced system profile and anatomical summary (Table 5), and (iii) paired fMRI-strength and EEG-spectral summaries indexed by the same component (Figures 7 and 8). Because atlas compositions use absolute loadings rather than the signed cortical maps (Figure 6), atlas-based summaries should be interpreted as coarse composition rather than as a one-to-one network labeling: many components naturally exhibit cross-system mixtures that reflect continuous macroscale organization (e.g., hybrid/gradient-like patterns) learned in a geometry- and bandwidth-constrained coordinate system.

### Methodological considerations and limitations

We emphasize that EEG and fMRI connectivity are distinct observables; the goal of fusion is to infer shared latent structure under modality-specific bandwidth limits, rather than to enforce edge-by-edge agreement between EEG and fMRI connectomes. Several limitations should be noted. First, we used age as the primary validation target because it is well powered in MPI–LEMON and has well-replicated associations with both rs-fMRI connectivity and resting EEG; however, the dataset includes additional phenotypes that could be used to test the generality of the fused features beyond aging. Second, EEG connectivity in source space depends on preprocessing and inverse-modeling choices.

Although we used a leakage-robust envelope-covariance measure and explicitly restricted EEG to coarse spatial modes, residual source mixing and model mismatch may remain and could influence fine-grained network estimates. Third, our estimation targets MAP point estimates; uncertainty quantification for networks and subject-level scores is not provided in the current implementation. Fourth, the LB basis is computed on a template cortical surface to provide a shared coordinate system; this choice necessarily abstracts away subject-specific geometry and may limit sensitivity to individualized cortical folding or spatial idiosyncrasies. Finally, the feature-on-age regressions are descriptive association analyses rather than causal effects, and should be interpreted as characterizing systematic age-related variation in the fused connectivity summaries.

### Why a common cortical eigenbasis is central to the latent-field EEG–fMRI fusion perspective

Our modeling perspective rests on three pillars: (i) a latent spatio-spectral connectivity object Σ_*i*_(*k, k*^′^; *ω*), (ii) a notion of spatial bandwidth indexed by *k*, and (iii) modality-specific partial observations over (*k, k*^′^, *ω*) (“structured holes” in the joint spatio-spectral space). While EEG–fMRI fusion can certainly be performed in ROI/parcellation space by fitting low-rank models to modality-specific connectomes, which can work well for prediction, they weaken the specific latent-field story emphasized here. All three become substantially cleaner when *k* indexes eigen-functions of a shared operator on the cortical manifold (e.g., Laplace–Beltrami eigenmodes), because *k* then corresponds to *spatial frequency* (coarse-to-fine scale) rather than an arbitrary feature index and forms a coordinate system that depends only on cortical shape, not on any particular modality, task, or parcellation. In parcel index space, “spatial bandwidth” is atlas-dependent and lacks a principled notion of wavelength, geometry-informed priors are harder to motivate (what is “high spatial frequency” on an arbitrary atlas graph?), and the structured-hole interpretation in (*k, k*^′^, *ω*) largely collapses towards the weaker statement that some edges are noisier than others. In our setting, fMRI provides broad spatial coverage but only a low-frequency temporal aggregate, while EEG provides frequency-resolved information but only for coarse spatial scales. This yields a geometrically meaningful missingness pattern in (*k, k*^′^, *ω*) space: roughly, “all *k* but very low *ω*” for fMRI, and “broad *ω* but 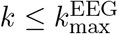 for EEG. Consequently, inferring unobserved bands via shared low-rank structure and regularization becomes constrained inference in a well-defined domain.

The same coordinate system also strengthens interpretability and directly underlies our main empirical payoffs. Under the low-rank factorization 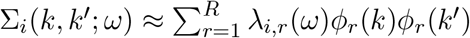, each factor couples (i) a *single spatial network ϕ*_*r*_ expressed in spatial-frequency coordinates and (ii) an *oscillatory signature λ*_*i,r*_(*ω*) for that same network. Because *k* is ordered by spatial frequency, we can define network-specific spatial-scale summaries (e.g., the effective spatial-frequency index in Section 4.6) and relate them directly to spectral profiles. Finally, although the LB basis is global and the learned networks are generally continuous rather than parcellated, they can be summarized in standard neuroanatomical terms via back-projection to the template surface: *ϕ*_*r*_ induces a vertex-wise map *ψ*_*r*_ = *U ϕ*_*r*_ (where *U* denotes the matrix of template LB eigenmodes), enabling post-hoc quantification of overlap with canonical atlases (e.g., Schaefer/Yeo) without altering the model’s native representation.

### Future directions

Several extensions follow naturally from the latent-field perspective. First, one can move beyond within-frequency connectivity by modeling cross-frequency coupling, for example through an object Σ_*i*_(*k, k*^′^; *ω, ω*^′^) and structured low-rank decompositions that incorporate frequency-domain kernels (Canolty and Knight, 2010; Aru et al., 2015; Nadalin et al., 2019). Second, the subject-specific spectral strengths *λ*_*i,r*_(*ω*) can be embedded in hierarchical models to support group-level inference and covariate effects directly on frequency-resolved network profiles, enabling richer hypothesis tests while retaining the same geometry-aligned, bandwidth-aware foundation. Hierarchical Bayesian variants could provide uncertainty for network maps and spectral profiles, and priors could encourage structured sparsity or system-specific organization when appropriate (Chen et al., 2016; Gorbach et al., 2020).

### Simultaneous EEG–fMRI and time-resolved coupling

The present work focuses on time-averaged connectivity, treating rs-fMRI and rs-EEG as complementary observations of a latent spatio-spectral connectivity field. An important next step is to extend the framework to *simultaneous* EEG–fMRI, where temporal alignment enables direct modeling of cross-modal dynamics rather than only static connectivity summaries. Such extensions would also connect naturally to work on connectivity-based identifiability and dynamic brain fingerprinting, which asks when time-varying connectome patterns are most individually informative (Amico and Goñi, 2018; Van De Ville et al., 2021). More broadly, these extensions would enable mechanistic tests of how fast electrophysiological coupling relates to slower BOLD co-fluctuation structure within a shared geometry-defined coordinate system, including questions about scale–frequency organization across modalities (Wang et al., 2012; Koller et al., 2024).

### Conclusion

Cortical eigenmode coordinates provide a principled geometry-defined scaffold for multimodal connectivity fusion. By combining a masked spatio-spectral latent-field formulation with PSD low-rank structure and smooth nonnegative spectral weights, the proposed approach yields compact and interpretable EEG–fMRI network features. In the MPI–LEMON application, these features show clear multimodal complementarity for age prediction and exhibit coherent age-associated variation in both spatial-strength and oscillatory summaries, illustrating their potential as practical multimodal network biomarkers for individual-differences research.

## Ethics

All analyses used de-identified human data from the publicly available MPI–LEMON dataset (Babayan et al., 2019). Ethical approval and informed consent procedures for participant recruitment and data collection were obtained by the original study investigators; the present work involves secondary analysis of these anonymized data.

## Data and Code Availability

The MPI–LEMON resting-state EEG and fMRI data analyzed in this study are publicly available (see Babayan et al., 2019 for access and dataset details).

## Acknowledgements

The author thanks the MPI–LEMON investigators for making the dataset publicly available. Computations were performed on NYU’s High Performance Computing (HPC) resources, including the BigPurple cluster.

## Supplementary Material

Supplementary material is included in this submission PDF.

## Supplementary Materials

### Overview

This supplement provides (i) additional details on fusion-model estimation, (ii) details on constructing LB-domain rs-fMRI and rs-EEG connectivity objects in the MPI–LEMON cohort, and (iii) expanded simulation and application results.

## S1. Model estimation details

This section provides implementation details for the alternating minimization procedure used to fit the fused connectivity model in the main text (Section 2.7). The optimization alternates between (A) updating the nonnegative EEG frequency-resolved weights 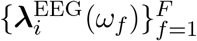 (and, in the separate-fMRI configuration, the nonnegative fMRI network-strength vector 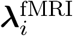) and (B) updating the shared orthonormal spatial factors Φ ∈ ℝ^*K*×*R*^ on the Stiefel manifold 𝕍_*K,R*_ = {Φ ∈ ℝ^*K*×*R*^ : Φ^⊤^ Φ = *I*_*R*_}. When ARD is enabled, an additional step updates the factor-wise precisions *τ* (Section 2.6).

### Discrete spectral weights

In our implementation, EEG spectral profiles are represented directly on a discrete frequency grid 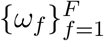 via the nonnegative weights 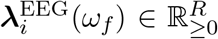. For each factor *r* and subject *i*, smoothness across frequency is enforced by a second-difference penalty over the index *f* (Section S1.2), rather than by an explicit spline basis.

### S1.1. Vectorization and design matrices

Let vech(·) denote half-vectorization that stacks the diagonal and upper-triangular entries of a symmetric matrix (including the diagonal). We further restrict vech(·) to those upper-triangular indices where the *corresponding mask entry is nonzero*, so that unobserved or unresolvable entries are excluded in the least-squares system. This constructs the index set from the upper triangle of each mask.

#### EEG blocks

For subject *i* and frequency bin *ω*_*f*_, define the masked observation vector

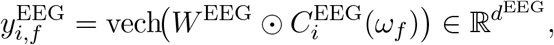

where *d*^EEG^ is the number of retained (upper-triangular) entries under the EEG mask. For each factor *r* = 1, …, *R*, define the corresponding design column

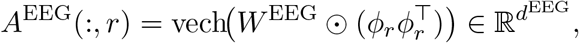

so that the model implies 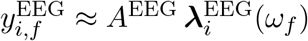.

#### fMRI block (separate-fMRI configuration)

Similarly, define

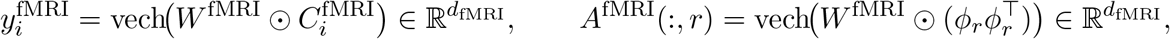

so that 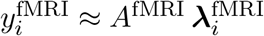. Although 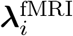 is an fMRI-strength vector, it is estimated *jointly* with 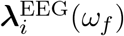 through the shared Φ.

#### Stacked system (per subject)

Let

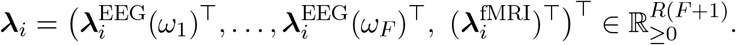

We form a weighted linear system ***y***_*i*_ ≈ *A*_*i*_***λ***_*i*_ by stacking (i) EEG blocks across frequencies and (ii) one fMRI block. In the implementation, the data-fit weights enter as multiplicative factors 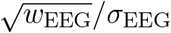 and 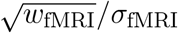. Concretely,

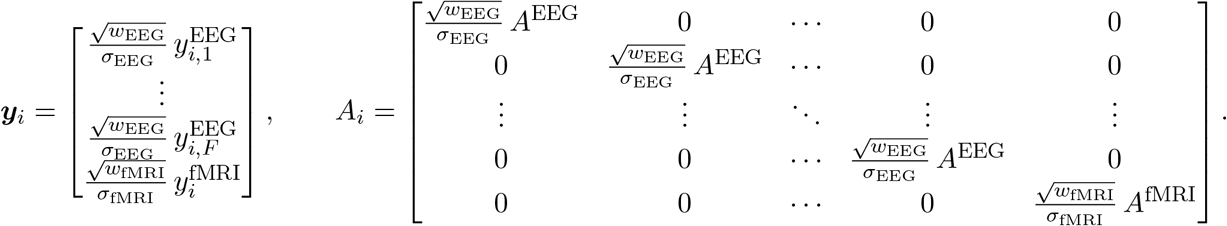

This representation makes explicit that EEG weights enter only their corresponding frequency-specific blocks, while the fMRI network-strength vector enters only the fMRI block.

### S1.2. (A) Update of nonnegative weights via NNLS

Fixing Φ, the update for each subject *i* is a convex penalized nonnegative least-squares (NNLS) problem over the stacked vector ***λ***_*i*_:

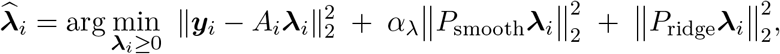

where *P*_smooth_ applies the second-difference penalty to the EEG portion of ***λ***_*i*_ only, and *P*_ridge_ denotes either (i) a small global ridge with weight *α*_0_ (when ARD is disabled) or (ii) an ARD factor-wise ridge determined by *τ* (when ARD is enabled). This NNLS update decouples across subjects and is parallelized across *i* in the implementation.

#### Augmented NNLS form

We solve the above by converting it to a standard NNLS problem on an augmented system:

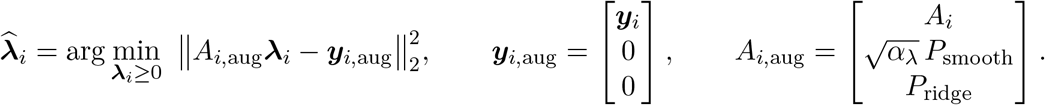

#### Smoothness operator

Let *D* ℝ^(*F* −2)×*F*^ denote the second-difference matrix over the frequency index *f*. Because 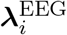 is stacked in *frequency-major* order 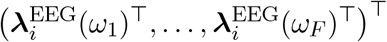, the corresponding block-smoothness operator is *D* ⊗ *I*_*R*_ ∈ ℝ^(*F* −2)*R*×*FR*^. We pad zeros for the last *R* entries (the fMRI block), yielding

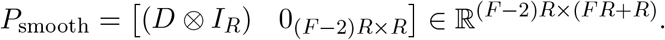

#### Ridge / ARD operator

When ARD is disabled, the implementation uses a small global ridge *α*_0_, corresponding to 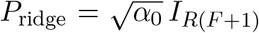. When ARD is enabled with factor-wise precisions *τ*_1_, …, *τ*_*R*_, the implementation uses a factor-wise ridge that is *repeated across frequencies* for EEG and (in the separate-fMRI configuration) also applied to the fMRI block:

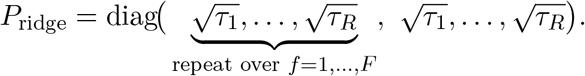

We solve the augmented NNLS problem using bounded-variable least squares with bounds (0, ∞).

### S1.3. (B) Update of Φ on the Stiefel manifold

Fixing 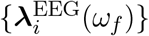 and 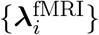, we update Φ under the orthonormality constraint Φ^⊤^ Φ = *I*_*R*_ using a projected gradient step on the Stiefel manifold followed by QR retraction.

Let *G* = ∇_Φ_ 𝒥(Φ, *λ*) denote the Euclidean gradient of the objective. We project to the tangent space *T*_Φ_𝕍_*K,R*_ = {*Z* ∈ ℝ^*K*×*R*^ : Φ^⊤^ *Z* + *Z*^⊤^Φ = 0} via

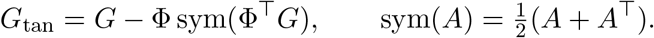

We then take a descent step and retract back to 𝕍_*K,R*_ using QR:

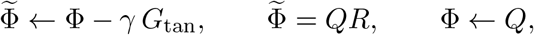

so that Φ ∈ 𝕍_*K,R*_. In the implementation, we use backtracking line search on *γ*: starting from *γ*_0_ (default 10^−2^), we repeatedly shrink *γ* ← *ργ* (we used *ρ* = 0.5) until the objective decreases or a maximum number of trials is reached.

### S1.4. Initialization, stopping, and practical settings

#### Initialization

By default, we initialize Φ from the top-*R* eigenvectors of the group-mean fMRI connectivity 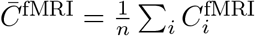 (followed by QR orthonormalization) if fMRI is used (*w*_fMRI_ *>* 0). If fMRI is not used, we initialize Φ from a random Gaussian matrix followed by QR. When a warm-start fit is provided, we initialize Φ by truncating (or expanding) the warm-start value Φ and then QR-orthonormalizing. All 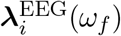 and 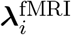 are initialized to a small positive constant (default 10^−2^).

#### Stopping criterion

We iterate (A)–(B) until the relative change in the objective satisfies |𝒥 ^(*t*)^ − 𝒥 ^(*t*−1)^ |*/*| 𝒥 ^(*t*−1)^ |*< ϵ* (default *ϵ* = 10^−6^), or until a maximum number of outer iterations is reached.

#### Modality weights and scaling

We use *w*_fMRI_ = 1 and *w*_EEG_ = 1*/F* so that the EEG contribution is averaged over frequency bins. In real data, we additionally apply robust global rescaling factors to each modality prior to fitting (Supplementary Section S6), to mitigate differences in raw connectivity magnitudes.

#### End-of-fit factor sorting

We sort factors after optimization by decreasing median total energy (median over subjects, and over frequency bins for EEG). This permutation is applied only at the end of the fit and does not affect estimation.

#### ARD settings

When ARD is enabled, we use the penalty described in Section 2.6 with an optional damped empirical-Bayes update for *τ* ; 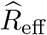 is computed by thresholding the fitted factor-energy curve.

##### Algorithm 1

Alternating minimization for fused EEG–fMRI connectivity factorization

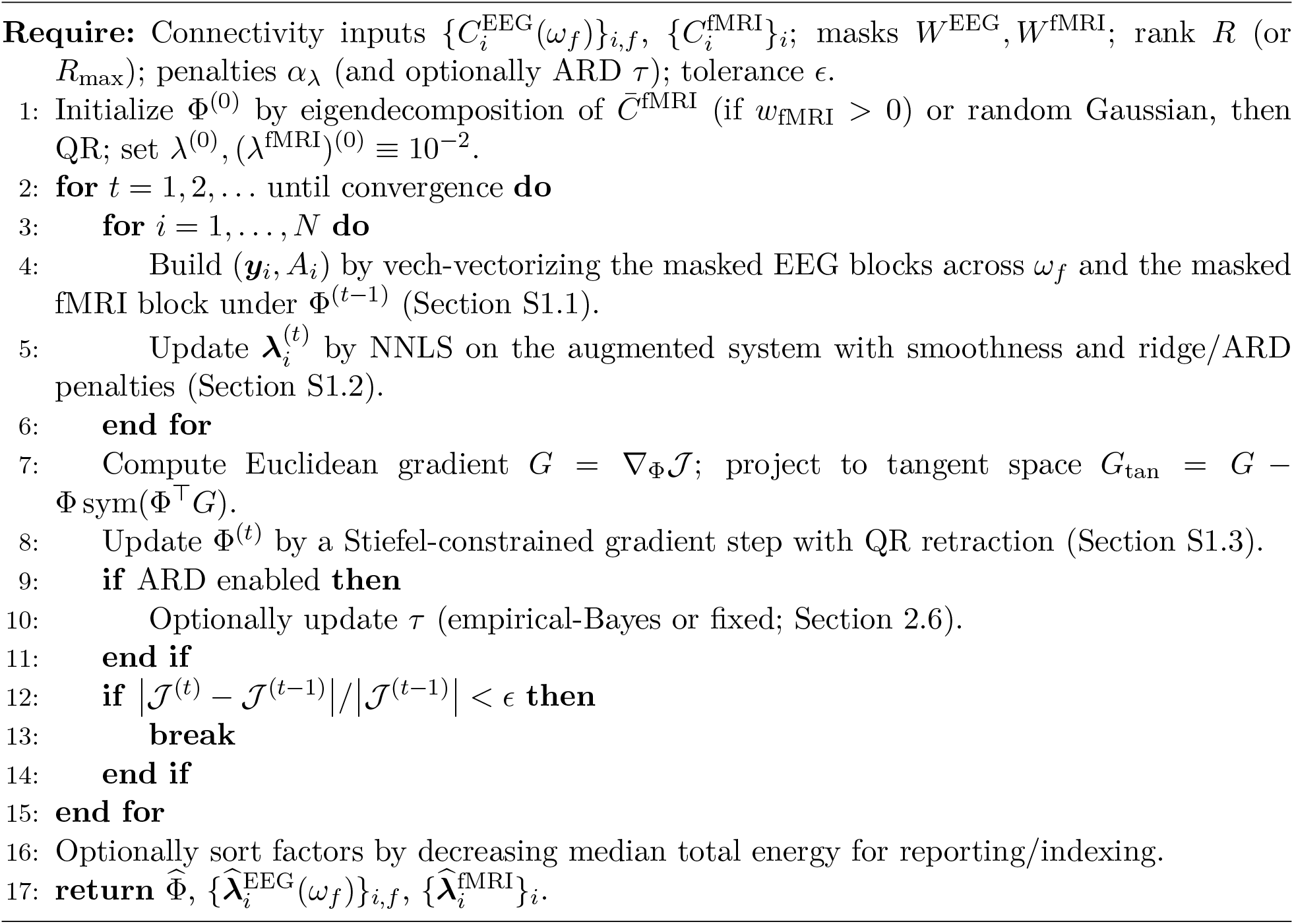

## S2. Template geometry and LB basis construction

### Shared LB representation

Both modalities were represented in the same template cortical eigen-mode coordinate system on `fsaverage5`. Let *K* denote the retained LB dimension (here *K* = 50 for model fitting). The model inputs (for each subject *i*) are: (i) an rs-fMRI LB-domain connectivity matrix 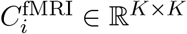, and (ii) frequency-resolved EEG LB-domain connectivity matrices 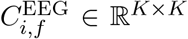 over a grid 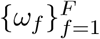 (here *F* = 20 bins over [1, 45] Hz). EEG unreliability at high spatial frequencies is encoded by a fixed mask *W* ^EEG^ on (*k, k*^′^).

### S2.1. Overview and shared notation

Let the cortical template mesh be `fsaverage5` with *V* vertices (left+right hemispheres; *V* = 20484 for `fsaverage5` ico-5). Let Φ ∈ ℝ^*V* ×*K*^ denote the first *K* Laplace–Beltrami (LB) eigenmodes on this template surface, ordered by increasing spatial frequency (eigenvalue). Let **a** ∈ ℝ^*V*^ be the vertex-wise area (mass) weights and define *A* = diag(**a**). For any vertex-domain signal **y**(*t*) ∈ ℝ^*V*^ (for time *t*), we define LB coefficients **b**(*t*) ∈ ℝ^*K*^ using the vertex-wise area *A*-weighted least-squares projection (Section S2.3).

#### Vertex ordering consistency between MNE source estimates and LB eigenmodes

To ensure that the LB eigenmode matrix Φ ∈ ℝ^*V* ×*K*^ is aligned with the vertex ordering used by MNE source estimates, we explicitly construct a template vertex index from the MNE `fsaverage5` source space. Let `src` denote the MNE source space created with `fsaverage5` ico-5. MNE stores the per-hemisphere vertex numbers as `src[0][`‘`vertno`‘] (left hemisphere) and `src[1][`‘`vertno`‘] (right hemisphere), and source time series are ordered as left-hemisphere vertices followed by right-hemisphere vertices. We therefore define the canonical vertex ordering: 𝒱_MNE_ = (`src[0][`‘`vertno`‘], `src[1][`‘`vertno`‘]), and let *V* = `|src[0][`‘`vertno`‘`]|`+`|src[1][`‘`vertno`‘`]|`. We store the LB eigenmodes Φ’s vertex dimension in this same concatenated order 𝒱_MNE_ before forming projections. With this alignment, for each epoch we stack MNE source time series as *s*_*i*_(*t*) = [*s*_*i*,lh_(*t*)^⊤^, *s*_*i*,rh_(*t*)^⊤^]^⊤^ ∈ ℝ^*V*^ and compute LB coefficients via *b*_*i*_(*t*) = (Φ^⊤^*A*Φ)^−1^Φ^⊤^ *A s*_*i*_(*t*) without hemispheric or vertex-index mismatch.

#### Why the mass/area matrix A is needed in LB projection

On a continuous surface, LB eigen-functions are orthonormal under the surface integral inner product:

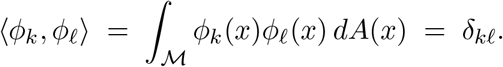

On a discrete mesh, the analogous inner product is represented by the finite-element *mass matrix A* (or a diagonal approximation using per-vertex areas). Using *A* ensures that (i) projections 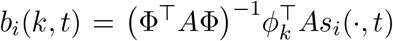 approximate the continuous inner product, (ii) mode energies are comparable across spatial scales, and (iii) orthonormality is respected numerically. In practice we use a diagonal area-weight approximation.

### S2.2. Template LB basis and area weights

We computed LB eigenmodes on the `fsaverage5` pial surface separately for left and right hemispheres using `LaPy` (`Solver(TriaMesh)`), removed the DC mode, and sign-aligned right-hemisphere modes to the left hemisphere via mirrored-vertex nearest-neighbor pairing. The final basis was stacked in template vertex order as

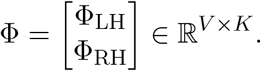

We also stored the vertex-wise area weights **a** ∈ ℝ^*V*^ for consistent *A*-weighted projections across fMRI and EEG.

### S2.3. A-weighted least squares projection

For each time point *t*, we project using weighted least squares:

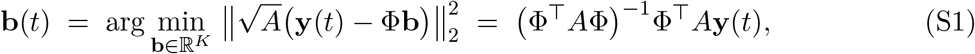

implemented via a numerically stable Cholesky factorization of the Gram matrix *G* = Φ^⊤^*A*Φ.

## S3. MPI–LEMON dataset and sample definition

**Table S1:** MPI–LEMON sample definition for the fusion analysis (eyes open). Counts reflect sequential inclusion criteria.

| Criterion | $n$ |
| --- | --- |
| Participants with usable rs-fMRI LB connectome | 212 |
| Among these, participants with usable EEG LB connectivity | 189 |
| Final fusion cohort (usable fMRI + usable EEG) | 189 |

We used the publicly available MPI–LEMON cohort and focused on the resting eyes-open condition to align with the resting-state fMRI instruction. We defined the rs-fMRI cohort as participants with successful fMRIPrep outputs and successful LB-domain connectome construction (Section S4), and the EEG cohort as participants with available consortium-preprocessed EEG files and successful LB-domain EEG connectivity construction (Section S5), yielding *n* = 189 participants with both modalities available after quality control (Table S1).

## S4. Resting-state fMRI preprocessing and LB-domain connectivity

### Overview

Resting-state fMRI (rs-fMRI) preprocessing and cortical surface sampling followed an `fMRIPrep`-based workflow. For each subject, preprocessed surface BOLD time series in `fsaverage5` space were denoised using a standardized nuisance-regression and temporal-filtering procedure. The denoised vertex time series were then projected to a Laplace–Beltrami (LB) eigenmode basis to obtain subject-specific LB coefficients, from which we formed an LB-domain covariance matrix per subject. This representation yields symmetric positive semidefinite (PSD) connectivity objects by construction, consistent with the PSD parameterization used in our modeling framework.

### S4.1. Resting-state fMRI preprocessing (fMRIPrep)

We processed MPI–LEMON rs-fMRI (`ds000221`, `ses-01`) using `fMRIPrep` (executed via Singularity on HPC cluster). We ran the `fMRIPrep` workflow at full processing level. Each subject was run as an independent task. Each task staged a per-subject BIDS directory containing the subject’s `ses-01` data. The `fMRIPrep` derivatives used were: (i) surface-sampled BOLD time series in `fsaverage5` space for each hemisphere (GIFTI functional time series) and (ii) the corresponding confound regressors.

#### Surface time series assembly

For each subject, we loaded the left- and right-hemisphere `fsaverage5` surface BOLD time series and concatenated hemispheres in fixed template vertex order (LH then RH) to form a matrix *Y* ∈ ℝ^*T* ×*V*^, where *T* is the number of time points and *V* is the total number of `fsaverage5` surface vertices across hemispheres.

#### Nuisance regression and filtering

For each subject, we performed nuisance regression using a standard confound set extracted from the fMRIPrep confounds table. The nuisance set comprised: (i) 6 rigid-body motion parameters; (ii) their first temporal derivatives and squared terms for the motion and derivative regressors; (iii) the first 10 aCompCor components; and fMRIPrep outlier regressors (motion and non-steady-state). An intercept and a linear time trend were included in the design matrix. For each vertex time series, confounds were regressed out via least squares to yield residual time series. The residual time series were then band-pass filtered to 0.01–0.10 Hz using an FIR filter and z-scored across time at each surface vertex prior to LB projection and downstream covariance estimation.

#### S4.1.1. LB projection and fMRI connectivity construction

For each subject, we projected the denoised vertex vector **y**(*t*) ∈ ℝ^*V*^ to LB coefficients via the *A*-weighted least squares solve (S1): *B*(*t*, :)^⊤^ = **b**(*t*) = *G*^−1^Φ^⊤^ *A* **y**(*t*). Let *B* ∈ ℝ^*T* ×*K*^ be the LB coefficient matrix (stacking across time) and 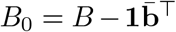 be column-centered coefficients. We computed the unbiased sample covariance of the (mean-centered) LB coefficient time series:

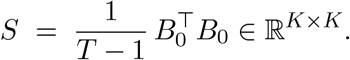

which is symmetric PSD by construction.

The primary fMRI LB cohort was defined as subjects for which project-level LB covariance outputs were successfully generated and passed basic structural checks (finite values, symmetry, and valid diagonals). In this work, this yielded *n* = 212 subjects.

## S5. Resting EEG preprocessing, source model, and LB-domain connectivity

### Overview

We constructed frequency-resolved EEG connectivity objects in the same `fsaverage5` LB coordinate system used for rs-fMRI. We started from consortium-distributed preprocessed EEG (EEGLAB), segmented continuous recordings into boundary-aware 10 s windows, estimated cortical source activity using a template-only linear inverse model, and projected source time series to LB coefficients. We then computed band-resolved LB-domain connectivity using a leakage-robust envelope-covariance measure across *F* = 20 frequency bins. The resulting per-frequency connectivity matrices are symmetric positive semidefinite (PSD) by construction and serve as the EEG inputs to the fusion model.

### S5.1. EEG data preprocessing

We used the MPI–LEMON consortium-provided *preprocessed* EEG distributed in EEGLAB `.set/.fdt` format (downloaded from the GWDG mirror). As provided by the consortium, recordings were downsampled to 250 Hz, band-pass filtered (approximately 1–45 Hz), and cleaned via PCA+ICA with ocular and cardiac components removed. We did not apply additional artifact rejection beyond boundary-aware windowing (Section S5.2).

### S5.2. Canonical channels and boundary-aware windowing

EEG was loaded in MNE-Python from EEGLAB files and mapped to the standard sensor montage `standard_1005` (`on_missing=`‘`ignore`‘). We used a fixed canonical set of 59 channels^1^; for each participant we retained the subset present in the recording (no interpolation) while maintaining canonical ordering. Average reference was applied as a projection operator.

Continuous EEG streams contain EEGLAB `boundary` annotations marking concatenation points. To avoid epoching and filtering across discontinuities, we segmented recordings into non-overlapping 10 s windows and excluded any window overlapping a boundary annotation (`reject_by_annotation=True`). Connectivity estimates were computed within windows and then averaged across windows to obtain a single subject-level matrix per frequency bin.

### S5.3. Template inverse and LB coefficient extraction

We performed template-only source modeling using MNE-Python on the `fsaverage` template: an ico-5 source space on the pial surface, a standard 3-layer BEM (brain/skull/scalp) with standard EEG conductivities, and the template transform `fsaverage-trans.fif`. Forward and inverse operators were constructed once and then restricted to each subject’s available canonical channels.

Source estimates were computed with a standard linear minimum-norm estimate (MNE) inverse with depth weighting and loose orientation, using `pick_ori=`‘`normal`’ to obtain a single time series per cortical vertex on the `fsaverage5` cortical template (with orientation normal to the surface). We use MNE as the primary inverse because it is linear, numerically stable in template-only settings, and yields source estimates that can be projected directly onto the same `fsaverage5` LB eigenmodes used for fMRI. For each 10 s window, let **s**(*t*) ∈ ℝ^*V*^ denote the template vertex-wise source time series stacked in fixed template vertex order (LH then RH). We projected **s**(*t*) to LB coefficients using the same *A*-weighted least-squares projection used for rs-fMRI:

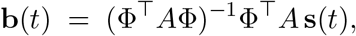

where Φ ∈ ℝ^*V* ×*K*^ denotes the template LB eigenmodes and *A* is the diagonal area-weight matrix (Section S2). To reflect limited EEG spatial bandwidth, we retained only low spatial modes for EEG connectivity, using 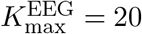.

### S5.4. Frequency-resolved EEG connectivity in LB space

For each subject, we computed frequency-resolved connectivity using *F* = 20 frequency bins spanning 1–45 Hz (log-spaced). Within each frequency bin, LB coefficient time series were band-pass filtered and connectivity was computed per 10 s window and then averaged across windows.

#### Leakage-robust envelope covariance

Let 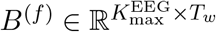 denote the LB coefficient time series within a 10 s window after band-pass filtering to frequency bin *f* (with *T*_*w*_ samples per window). To attenuate instantaneous linear mixing, we applied symmetric orthogonalization (regularized whitening):

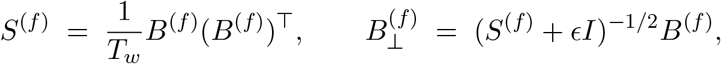

with a small ridge *ϵ* = 10^−4^ *>* 0 for numerical stability. This 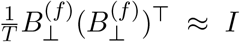, attenuating instantaneous linear mixing across modes. We then computed analytic amplitudes via the Hilbert transform, 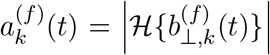, and defined the within-window connectivity as the envelope covariance 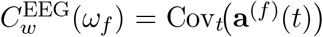. Finally, we averaged across windows to obtain the subject-level matrix 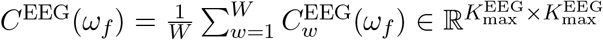. Because it is a covariance of envelopes, *C*^EEG^(*ω*_*f*_) is symmetric PSD by construction. We retained diagonal entries as mode-wise envelope variances.

## S6. Real-data model specification and tuning (MPI–LEMON)

### fMRI network strengths as a low-frequency summary

In real data, rs-fMRI connectivity is effectively a low-temporal-frequency summary and does not naturally admit a direct correspondence to the EEG frequency grid. We therefore parameterize rs-fMRI using its own nonnegative subject–network strengths 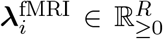, while EEG provides frequency-resolved nonnegative weights 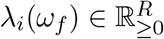 on *ω*_*f*_ ∈ ℱ_EEG_. Both modalities are coupled through the shared orthonormal spatial factors Φ.

### Cross-modality scaling (robust global normalization)

Due to modality-specific preprocessing, band averaging, and estimation noise, EEG and fMRI connectivity estimates can differ in overall magnitude. To place the modality residual terms on comparable numerical scales prior to fitting, we applied a robust global rescaling to each modality.

Let *W* ^EEG^ ∈ ℝ^*K*×*K*^ denote the fixed EEG spatial mask and *W* ^fMRI^ ∈ ℝ^*K*×*K*^ the fMRI mask (typically all ones for fMRI). For each subject *i*, define the masked Frobenius norms

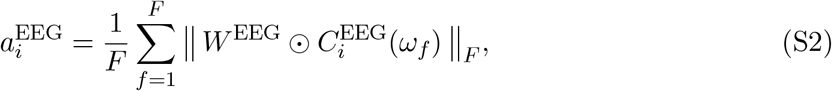

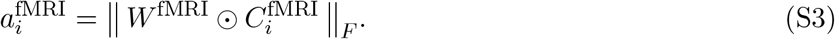

We then set robust modality scale factors as medians across subjects,

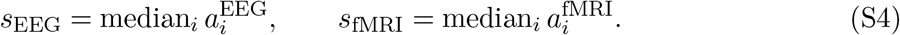

Finally, we rescaled the observed connectivity objects as

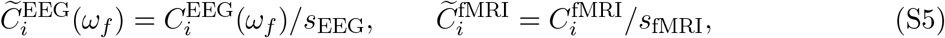

and fit the fusion model using 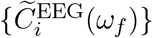 and 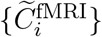. This normalization is robust to outliers (median-based), preserves the relative structure within each modality, and ensures that the two data-fit terms in the objective operate on comparable numerical scales.

**Figure S1:**
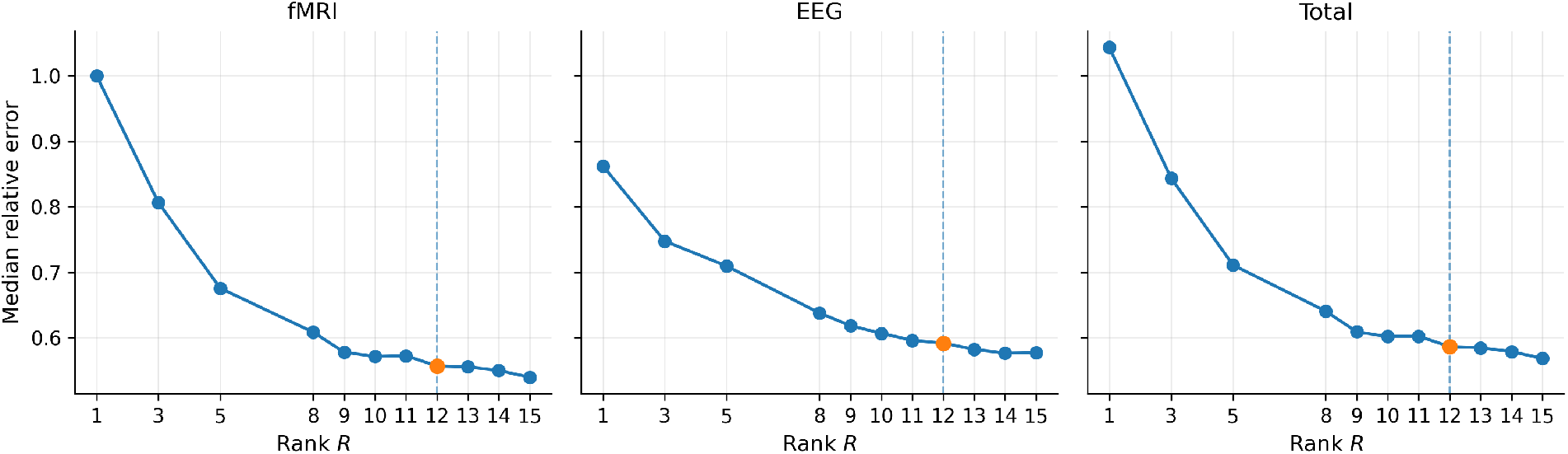
MPI–LEMON: rank-selection elbow plots from the rank sweep. Each point corresponds to a fixed-rank fit at *R* ∈ {1, 3, 5, 8, 9, 10, 11, 12, 13, 14, 15}. **Left:** median subject-level relative reconstruction error for fMRI connectivity, MedRelErr^fMRI^(*R*). **Middle:** median subject-level relative reconstruction error for EEG connectivity on the EEG-observed block, MedRelErr^EEG^(*R*). **Right:** combined diagnostic MedRelErr^Total^(*R*) = *w*_fMRI_MedRelErr^fMRI^(*R*) + *w*_EEG_MedRelErr^EEG^(*R*) using the same fixed weights as in the fitting objective (*w*_fMRI_ = 1, *w*_EEG_ = 1*/F*). The dashed vertical line marks the selected rank *R* = 12.

### Rank selection

We used two complementary rank diagnostics: (i) an over-parameterized ARD-based fit with *R*_max_ = 20 to obtain an effective-rank estimate 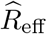 by thresholding the factor-energy curve at *E*_*r*_*/E*_max_ ≥ 0.01, and (ii) a rank sweep to construct elbow curves over candidate ranks. Specifically, for (ii), we fit an anchor model at *R* = 15, sorted factors by median total energy, and refit *R* ∈ {1, 3, 5, 8, 9, 10, 11, 12, 13, 14, 15} using truncated-anchor initialization. Elbow curves were then computed for fMRI, EEG, and a combined diagnostic as described below.

### Elbow-plot construction (rank sweep)

To assess rank-dependent reconstruction behavior, we performed a sweep over *R* ∈ {1, 3, 5, 8, 9, 10, 11, 12, 13, 14, 15} and extracted summary measures from each fitted model. For a given rank *R*, let 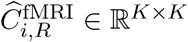 denote the model-predicted fMRI connectivity for subject *i* and let 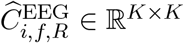 denote the model-predicted EEG connectivity at frequency bin *ω*_*f*_. Using the modality-specific masks *W* ^fMRI^ and *W* ^EEG^, we define subject-level relative errors

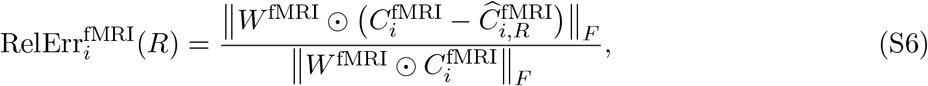

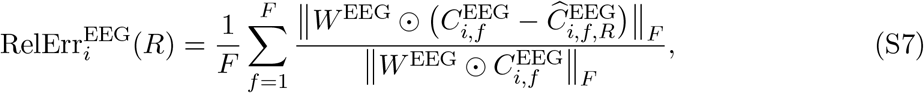

where *F* is the number of EEG frequency bins used in the model. We then summarize the above for each rank by the median across subjects,

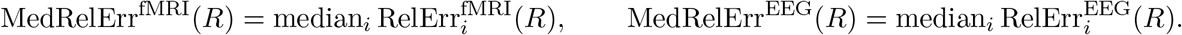

Finally, the elbow plot’s third panel (Figure S1) reports a combined diagnostic that matches the fitting objective weights,

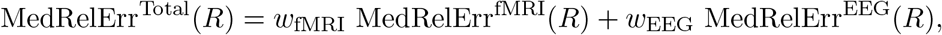

with *w*_fMRI_ = 1 and *w*_EEG_ = 1*/F*.

### Factor-energy ordering and typical spectral profiles

After selecting the working rank *R* = 12 (Fig. S1), we summarize the fitted networks using (i) a factor-energy ordering used throughout the application (Figs. S2 and S3) and (ii) typical EEG spectral profiles per network (Fig. S4).

**Figure S2:**
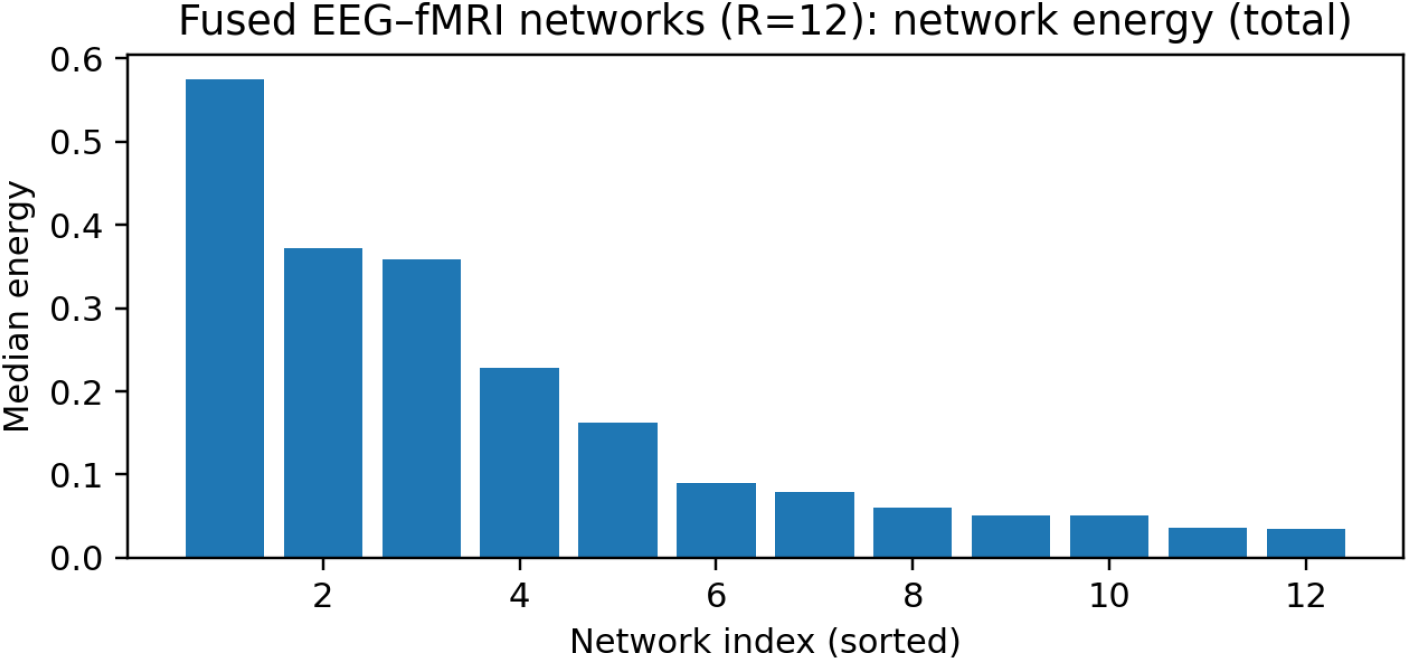
MPI–LEMON: per-network median total energy for the fitted rank-*R* = 12 model. Let 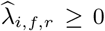 denote the fitted EEG weights (from 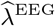) and 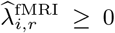 denote the fitted fMRI strengths (from 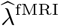). We define the EEG energy of network *r* as the median of squared EEG weights across subjects and frequency bins, 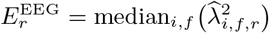, and the fMRI energy as the median of squared fMRI strengths across subjects, 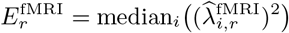. The plotted total energy is the sum of these two medians, 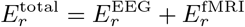. Networks are ordered by decreasing 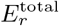 (the reporting order used throughout the application).

Specifically, we define per-network EEG and fMRI energies from the fitted nonnegative strengths and use their sum to order networks for reporting, downstream feature extraction, and the “Top-rank” energy screening procedure in the main text. Let 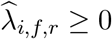 denote the fitted EEG weights (from 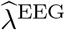) and 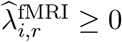 denote the fitted fMRI strengths (from 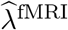). We define the EEG energy of network *r* as the median of squared EEG weights across subjects and frequency bins, 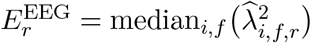, and the fMRI energy as the median of squared fMRI strengths across subjects, 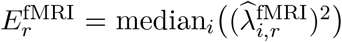. The plotted total energy in Figs. S2 and S3 is the sum of these two medians, 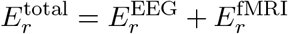.

We additionally visualize the across-subject median EEG spectral weight curve for each network to summarize its typical frequency-domain expression (Fig. S4).

## S7. ROI/system interpretation summaries (Schaefer-7)

We summarized the learned cortical network maps against a standard resting-state reference atlas using the Schaefer2018 parcellation (200 parcels; 7-network labeling) on `fsaverage5`. Let *ψ*_*r*_(*v*) denote the vertex-wise loading map of network *r* on `fsaverage5`, and let *V*_*p*_ be the set of surface vertices belonging to parcel *p* in the Schaefer2018 atlas. We summarize loadings using |*ψ*_*r*_(*v*)|.

### Parcel magnitude summaries

For each parcel *p*, we define the total absolute loading (*mass*)

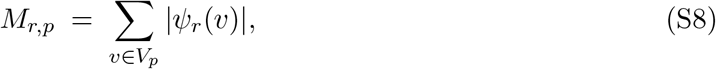

which is the parcel *p*-restricted *ℓ*_1_ magnitude of the map.

### System composition (percent mass)

System labels follow the Schaefer-7 nomenclature: **Default** (default-mode), **Cont** (frontoparietal control), **Vis** (visual), **SomMot** (somatomotor), **DorsAttn** (dorsal attention), **SalVentAttn** (salience/ventral attention), and **Limbic** (limbic). Let sys(*p*) ∈ {Default, Cont, Vis, SomMot, DorsAttn, SalVentAttn, Limbic} denote the Schaefer-7 system label of parcel *p*. We aggregate parcel masses over parcels within each system:

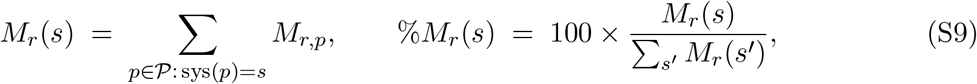

**Figure S3:**
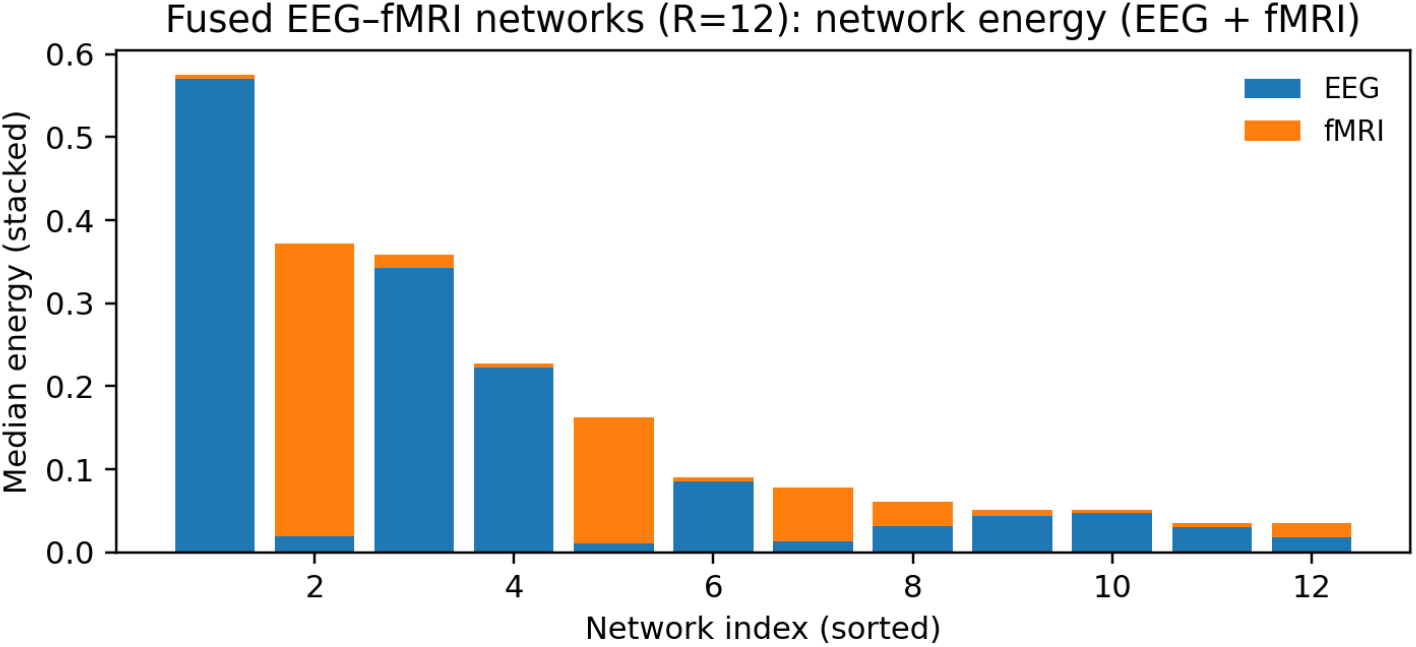
MPI–LEMON: stacked decomposition of per-network median energy into EEG and fMRI contributions (rank-*R* = 12). For each network *r*, the EEG component is 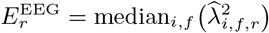 and the fMRI component is 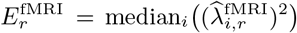. Bars are stacked as 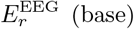 plus 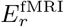 (top), so the bar height equals 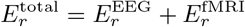 (sum of medians). Networks are ordered by decreasing 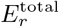.

where the % normalization yields a within-network composition (distribution) of the systems. The percent-mass values %*M*_*r*_(*s*) populate Figure S5 and the “Top systems” column of Table S2.

### Dominant systems/anatomy summaries

The “Dominant systems / anatomy” column in Table S2 is computed from a subset of parcels as follows to emphasize the network’s most informative regions. For each network *r*, we first rank parcels by *M*_*r,p*_ (bilateral pooling) in (S8) and retain the top *P* parcels (here *P* = 15). Within this top-*P* set, we (i) re-compute system totals ∑_*p*∈ top-*P* : sys(*p*)=*s*_ *M*_*r,p*_ and report the top *S* systems (here *S* = 3), and (ii) within each of those systems, list the most frequent/high-mass coarse anatomical tags derived from the Schaefer parcel-name components (top regions per system, here 3). This is different from the “Top systems (percent mass)” column, which uses all parcels and reports the global system composition of the full map.

The coarse tags correspond to standard abbreviations: PFC (prefrontal cortex), PFCl (lateral prefrontal cortex), PFCv (ventral prefrontal cortex), OFC (orbitofrontal cortex), Par (parietal cortex), Temp (temporal cortex), pCun/PCC (precuneus/posterior cingulate cortex; often labeled `pCunPCC`), FEF (frontal eye fields), FrOper/Ins (frontal operculum/insula; often labeled `FrOperIns`), PrC (precentral gyrus), Post (posterior cortical subdivision labels used in Schaefer naming, e.g., dorsal-attention “Post”), and Med (medial cortical subdivision labels used in Schaefer naming, e.g., salience/ventral-attention “Med”).

**Figure S4:**
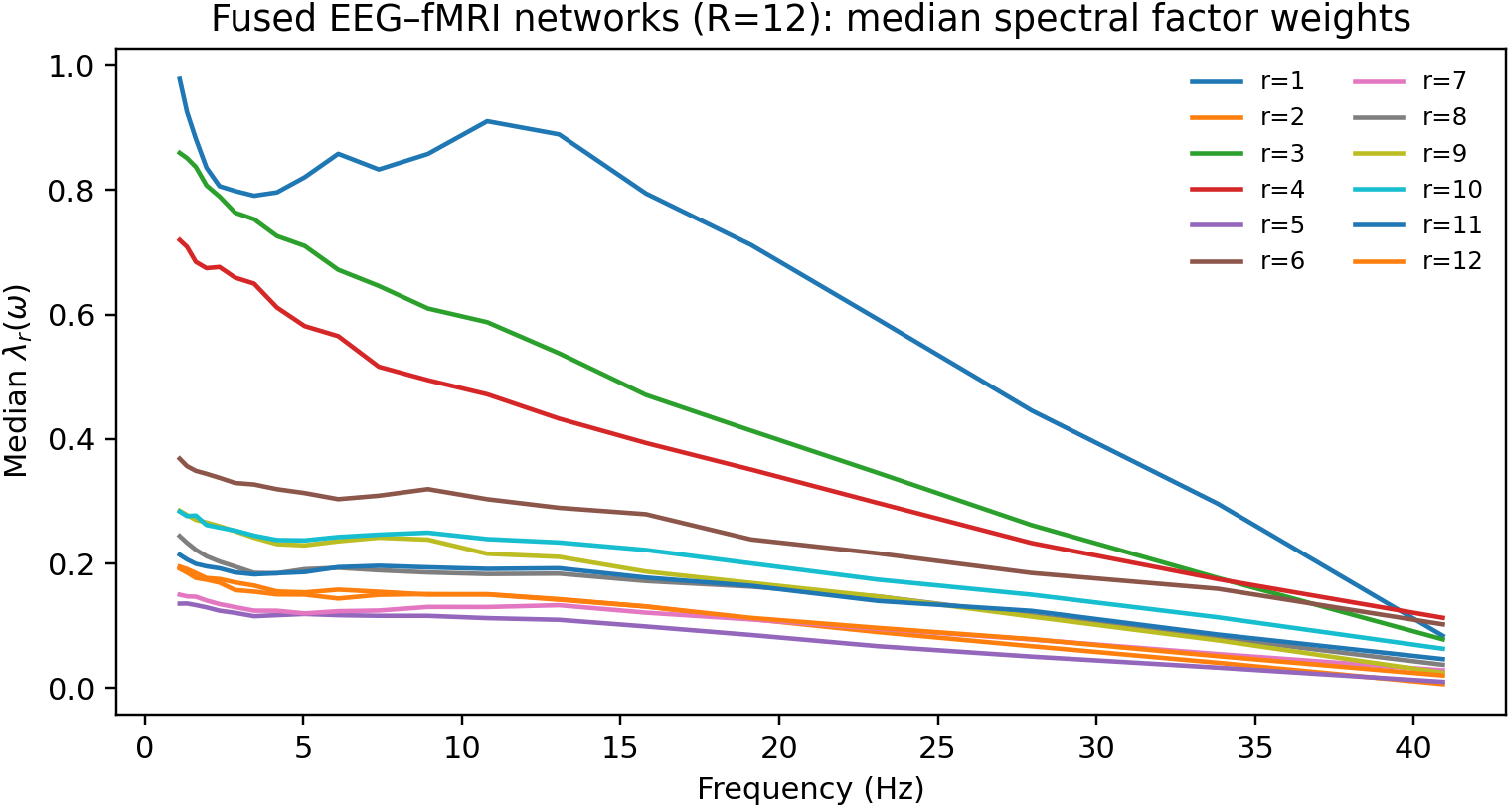
MPI–LEMON: median EEG spectral weight profiles for the fitted rank-*R* = 12 networks. For each network *r*, we plot the across-subject median of the fitted nonnegative EEG weights as a function of frequency, median_*i*_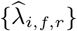 over the modeling frequency grid 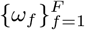. Networks are shown in the post-fit reporting order (decreasing median total energy; Fig. S2). This diagnostic summarizes typical spectral expression of each learned network in the fused representation.

**Figure S5:**
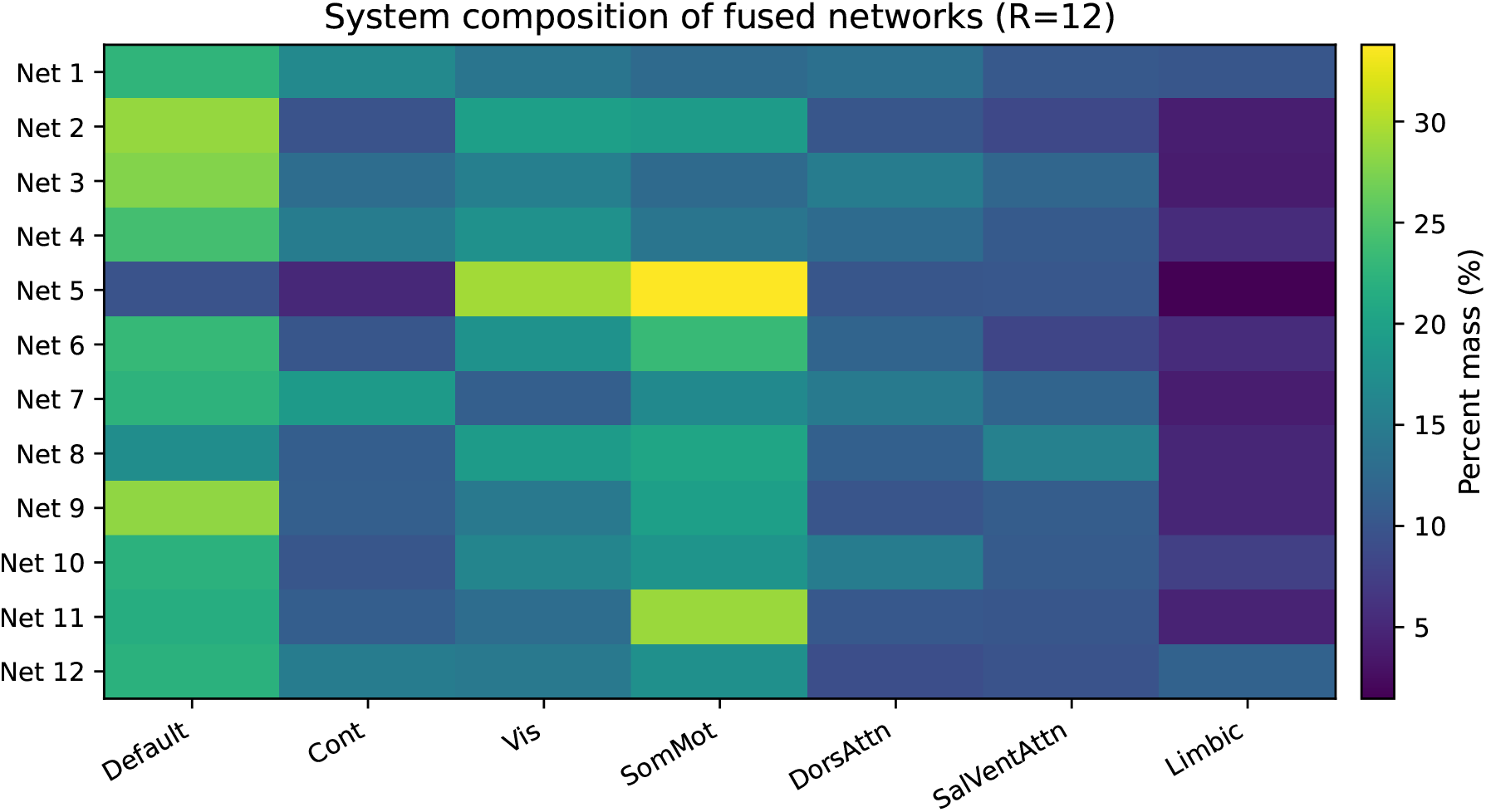
MPI–LEMON: Schaefer-7 system composition of the *R* = 12 fused networks. Each row is one learned network (Net *r*), and each column is a Schaefer2018 7-network system. Cell values are the *percent parcel mass* assigned to that system, computed by pooling left and right hemispheres and aggregating parcel-level magnitudes (absolute mean loading) with parcel size weighting (see Supplementary Methods Section S7). The heatmap provides a compact overview of how each learned network aligns with (and mixes across) canonical resting-state systems.

**Table S2:** Schaefer-7 system-level composition and dominant anatomy summaries (coarse anatomical labels) for each fused network (magnitude-based).

| Net $r$ | Top systems (percent mass) | Dominant systems / anatomy (bilateral) |
| --- | --- | --- |
| 1 | Default (22.6%), Cont (16.7%), Vis (14.0%) | **Cont** (PFCI), **Limbic** (OFC, TempPole), **Default** (Temp, PFC) |
| 2 | Default (28.6%), Vis (19.8%), SomMot (19.2%) | **Vis** (Vis), **Default** (pCunPCC, Temp, Par), **SomMot** (SomMot) |
| 3 | Default (27.8%), Vis (15.4%), DorsAttn (15.1%) | **Default** (Par, Temp, pCunPCC), **DorsAttn** (Post, PrCv), **Vis** (Vis) |
| 4 | Default (24.2%), Vis (17.8%), Cont (15.0%) | **Default** (Par, Temp, pCunPCC), **Vis** (Vis), **Cont** (PFCI, Par) |
| 5 | SomMot (33.8%), Vis (29.4%), SalVentAttn (10.3%) | **Vis** (Vis), **SomMot** (SomMot) |
| 6 | SomMot (23.2%), Default (23.1%), Vis (18.0%) | **Vis** (Vis), **SomMot** (SomMot), **Default** (pCunPCC, Temp) |
| 7 | Default (22.4%), Cont (19.1%), SomMot (16.8%) | **Cont** (PFCI, Par), **DorsAttn** (Post, PrCv), **Default** (pCunPCC, Temp) |
| 8 | SomMot (20.5%), Vis (19.2%), Default (17.3%) | **SalVentAttn** (FrOperIns, Med), **SomMot** (SomMot), **Vis** (Vis) |
| 9 | Default (28.4%), SomMot (19.8%), Vis (14.5%) | **Default** (Temp), **SomMot** (SomMot), **SalVentAttn** (FrOperIns, Med) |
| 10 | Default (22.2%), SomMot (18.2%), Vis (16.2%) | **DorsAttn** (Post, FEF), **SomMot** (SomMot), **Vis** (Vis) |
| 11 | SomMot (28.9%), Default (21.6%), Vis (13.0%) | **SomMot** (SomMot), **Vis** (Vis), **DorsAttn** (Post) |
| 12 | Default (22.2%), SomMot (17.7%), Cont (15.0%) | **Limbic** (TempPole, OFC), **Cont** (PFCI), **SomMot** (SomMot) |

### ROI/system composition summary (Schaefer-7)

Table S2 provides the resulting compact atlas-based summary of each fused network using the Schaefer2018 (200-parcel) atlas with 7-network labels. The “Top systems” column reports the global Schaefer-7 composition (%*M*_*r*_(*s*)) of each network across the full cortical map, while the “Dominant systems / anatomy” column focuses on the top-*P* parcels (by *M*_*r,p*_) to highlight the main contributing systems and the most informative coarse anatomical labels. Across networks, Default and Control contributions are prominent—especially among higher-energy components—while visual and somatomotor contributions are concentrated in specific networks (e.g., the visual-dominant and visual–somatomotor patterns). A consistent pattern is that many networks show substantial secondary system contributions rather than mapping one-to-one onto a single atlas label, which is consistent with the mixed-system/gradient-like appearance of several surface maps and supports interpreting the fused factors as distributed large-scale patterns bridging canonical systems. The two atlas summaries in Table S2 are complementary. The percent-mass composition (second column) and the heatmap (Fig. S5) reflect global distribution across systems, whereas the “dominant systems/anatomy” column (third column) highlights the most informative high-loading anatomical regions (based on the top-*P* parcels by mass).

### S7.1. Further interpretation of the representative latent spatio-spectral connectivity field

To summarize the fitted latent field at the cohort level, we formed a representative latent connectivity object

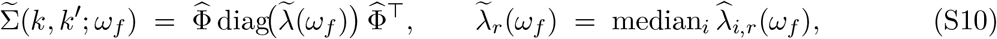

where 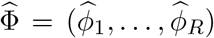 is the estimated shared LB-network basis and 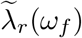 is the median subject-level strength of factor *r* at frequency *ω*_*f*_. This construction yields a representative positive semidefinite latent field in LB mode space (Fig. 10 of the main manuscript).

The diagonal summary

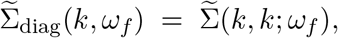

shows that latent variance is concentrated primarily in the first ∼ 10–12 LB modes, with the largest contributions at lower frequencies and a general decay in magnitude as frequency increases. Thus, the fitted field is dominated by coarse, large-scale cortical structure rather than being diffusely distributed across all spatial modes. In our data, one of the lowest modes is especially prominent in the 1–5 Hz range, whereas several intermediate modes are only weakly expressed across the spectrum. This pattern is best interpreted as uneven usage of the LB basis by the learned factor subspace, rather than as evidence that any single eigenmode is intrinsically “important” or “unimportant.”

The full *k* × *k* slices show that the dominant positive mode-by-mode covariance is concentrated in a low-*k* block (roughly the first ∼ 12 modes, and more weakly through about mode 20), whereas higher modes exhibit weaker and less coherent structure. At both 10 Hz and 30 Hz, the broad spatial pattern is similar, with the lower-frequency slice differing mainly in its larger overall magnitude. This is consistent with the model parameterization: because the spatial factors 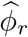 are shared across frequencies, frequency modulates the latent field primarily through the nonnegative strengths 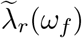 rather than by introducing entirely new spatial patterns at each frequency.

Diagonal entries are larger than most off-diagonal entries because they represent variances,

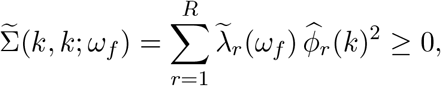

whereas off-diagonal entries,

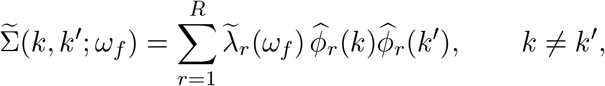

are signed covariances and can partially cancel across factors. Taken together, these diagnostics indicate that the fitted model produces a coherent latent spatio-spectral connectivity field that is dominated by coarse spatial scales, strongest at lower frequencies, and concentrated in a structured low-*k* coupling block.

## S8. Supplementary Results

### S8.1. Additional simulation results

#### Sensitivity of ARD effective-rank recovery to the energy threshold

We summarize ARD effective-rank recovery under two choices of the energy cutoff used to define 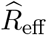 from the factor-energy curve *E*_*r*_*/E*_max_ (ARD-regularized fit with *R*_max_ = 20 in all conditions). Figure S6 uses a permissive threshold *E*_*r*_*/E*_max_ ≥ 0.005, while Fig. S7 uses a more conservative threshold *E*_*r*_*/E*_max_ ≥ 0.02. In both figures, boxplots show the distribution of 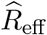 across Monte Carlo replicates for each (SNR, 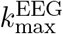) condition, stratified by true rank *R*^*^.

**Figure S6:**
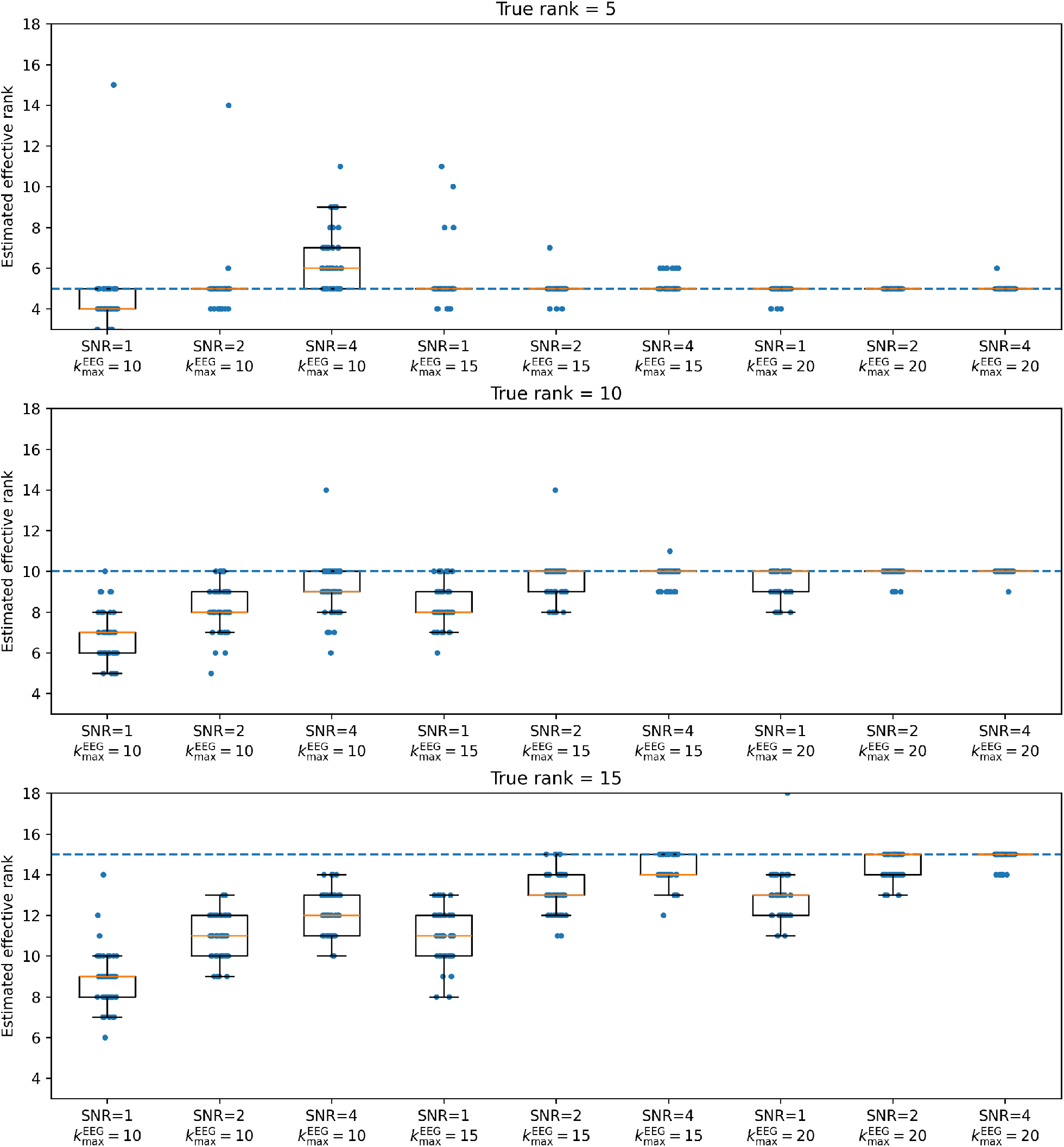
ARD effective-rank recovery. Boxplots show the distribution of the estimated effective rank 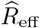 across Monte Carlo replicates for each (SNR, 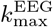) condition, stratified by true rank *R*^*\**^. 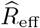 is defined by thresholding the factor-energy curve at *E*_*r*_*/E*_max_ ≥ 0.005 using an ARD-regularized fit with *R*_max_ = 20.

**Figure S7:**
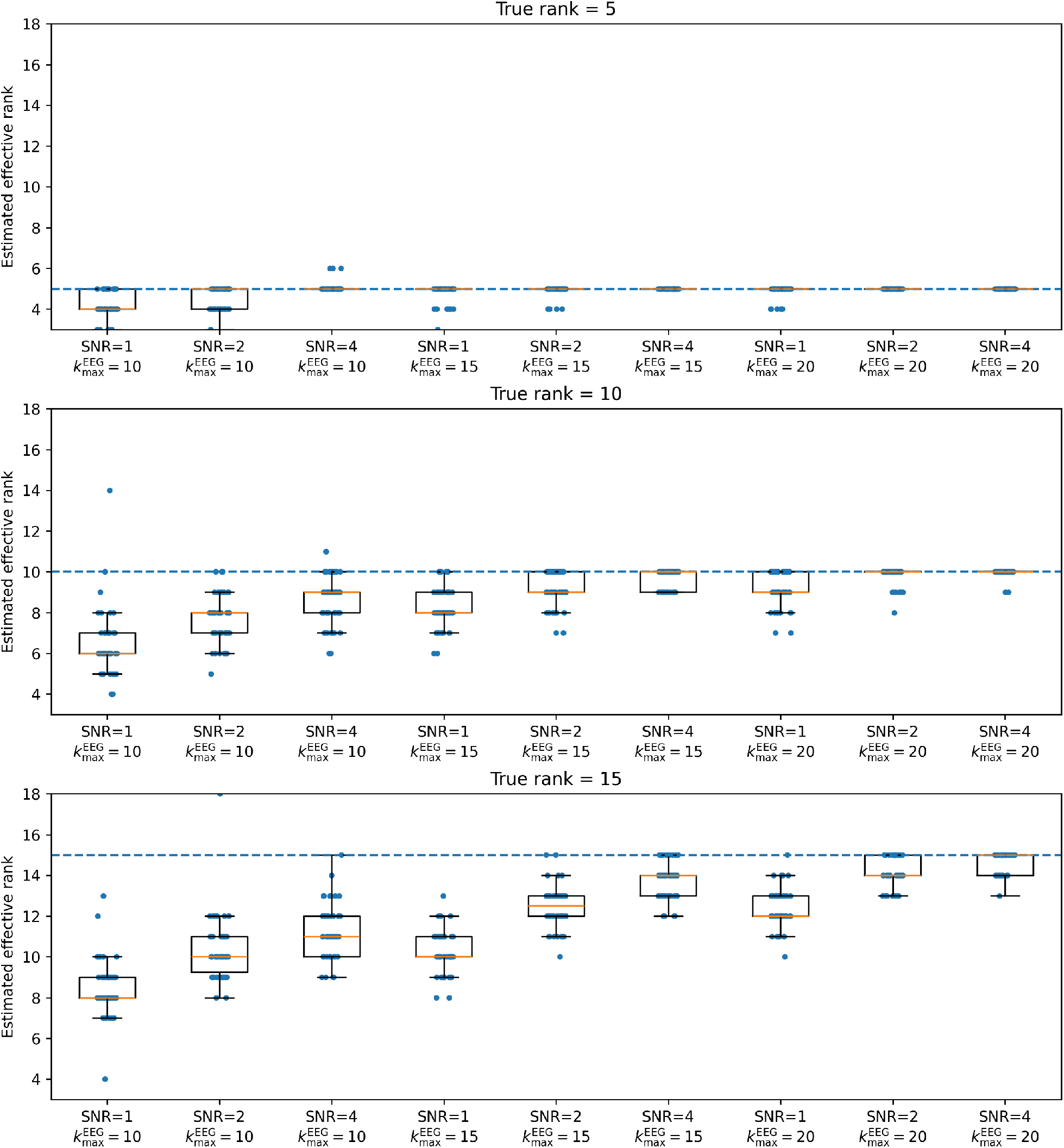
ARD effective-rank recovery. Boxplots show the distribution of the estimated effective rank 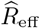 across Monte Carlo replicates for each (SNR, 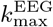) condition, stratified by true rank *R*^*\**^. 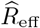 is defined by thresholding the factor-energy curve at *E*_*r*_*/E*_max_ ≥ 0.02 using an ARD-regularized fit with *R*_max_ = 20.

### S8.2. Age association screening and baseline comparisons

This section provides full results supporting the main-text age association screening in Section 4.4. We use the same feature-as-outcome regression direction (12) and report per-decade effects (10-year change) in SD units of each feature. Within each feature set, multiplicity is controlled using Benjamini–Hochberg FDR, and we report *q*-values.

#### Full fused-feature results (all networks)

Table S3 reports age associations for the complete fused feature set (*R* = 12). Each network contributes one spatial-strength feature and five spectral summaries (spectral centroid plus *θ/α/β/γ* band-mass summaries), yielding 6*R* fused tests.

#### Baseline feature sets

To contextualize the fused associations, Tables S4 and S5 report the corresponding EEG-only and fMRI-only baseline results. The EEG-only baseline includes spectral summaries for each network; the fMRI-only baseline includes the spatial-strength summary for each network. For comparability, tables are ordered by network ID (energy order) and feature type.

#### Sensitivity analysis: education adjustment

Finally, S6 repeats the primary Top-*K* = 5 fused analysis after adding education as an additional covariate (sex+education). The main patterns are robust, with modest attenuation for some effects as expected under additional adjustment.

**Table S3:** Age associations for all fused features across all networks (*K* = 12; *R* = 12), adjusting for sex (male). Each row corresponds to a univariate regression of z-scored feature on age (and covariates), with HC3 standard errors and BH-FDR correction within the fused feature set. Effects are reported per decade (10-year change) in SD units of the feature. Rows are ordered by network ID (energy order) and feature type.

| Network | Feature summary | $\beta$ (per decade) | SE | $q$ |
| --- | --- | --- | --- | --- |
| Network 1 | Network strength ($\lambda^{\text{fMRI}}$) | -0.156 | 0.035 | 1.90e-05 |
| Network 1 | Spectral centroid ($\mu^{\text{spec}}$) | 0.187 | 0.043 | 3.45e-05 |
| Network 1 | Band mass: $\theta$ | -0.126 | 0.033 | 2.48e-04 |
| Network 1 | Band mass: $\alpha$ | -0.124 | 0.032 | 2.22e-04 |
| Network 1 | Band mass: $\beta$ | -0.074 | 0.033 | 3.12e-02 |
| Network 1 | Band mass: $\gamma$ | 0.150 | 0.045 | 1.42e-03 |
| Network 2 | Network strength ($\lambda^{\text{fMRI}}$) | -0.139 | 0.040 | 9.40e-04 |
| Network 2 | Spectral centroid ($\mu^{\text{spec}}$) | 0.272 | 0.046 | 1.63e-08 |
| Network 2 | Band mass: $\theta$ | -0.155 | 0.031 | 2.29e-06 |
| Network 2 | Band mass: $\alpha$ | -0.112 | 0.033 | 1.08e-03 |
| Network 2 | Band mass: $\beta$ | -0.032 | 0.035 | 4.03e-01 |
| Network 2 | Band mass: $\gamma$ | 0.206 | 0.042 | 2.29e-06 |
| Network 3 | Network strength ($\lambda^{\text{fMRI}}$) | -0.244 | 0.032 | 4.52e-13 |
| Network 3 | Spectral centroid ($\mu^{\text{spec}}$) | 0.241 | 0.046 | 6.26e-07 |
| Network 3 | Band mass: $\theta$ | -0.140 | 0.032 | 3.72e-05 |
| Network 3 | Band mass: $\alpha$ | -0.110 | 0.033 | 1.42e-03 |
| Network 3 | Band mass: $\beta$ | -0.041 | 0.036 | 2.96e-01 |
| Network 3 | Band mass: $\gamma$ | 0.164 | 0.050 | 1.44e-03 |
| Network 4 | Network strength ($\lambda^{\text{fMRI}}$) | -0.239 | 0.034 | 1.36e-11 |
| Network 4 | Spectral centroid ($\mu^{\text{spec}}$) | 0.288 | 0.039 | 1.46e-12 |
| Network 4 | Band mass: $\theta$ | -0.092 | 0.036 | 1.35e-02 |
| Network 4 | Band mass: $\alpha$ | -0.056 | 0.037 | 1.54e-01 |
| Network 4 | Band mass: $\beta$ | 0.030 | 0.039 | 4.80e-01 |
| Network 4 | Band mass: $\gamma$ | 0.198 | 0.043 | 1.10e-05 |
| Network 5 | Network strength ($\lambda^{\text{fMRI}}$) | -0.132 | 0.036 | 3.87e-04 |
| Network 5 | Spectral centroid ($\mu^{\text{spec}}$) | 0.296 | 0.034 | 2.41e-17 |
| Network 5 | Band mass: $\theta$ | -0.137 | 0.033 | 6.92e-05 |
| Network 5 | Band mass: $\alpha$ | -0.095 | 0.034 | 7.65e-03 |
| Network 5 | Band mass: $\beta$ | 0.012 | 0.037 | 7.78e-01 |
| Network 5 | Band mass: $\gamma$ | 0.211 | 0.044 | 5.02e-06 |
| Network 6 | Network strength ($\lambda^{\text{fMRI}}$) | -0.228 | 0.037 | 1.89e-09 |
| Network 6 | Spectral centroid ($\mu^{\text{spec}}$) | 0.298 | 0.037 | 4.03e-15 |
| Network 6 | Band mass: $\theta$ | -0.140 | 0.033 | 4.44e-05 |
| Network 6 | Band mass: $\alpha$ | -0.115 | 0.033 | 7.99e-04 |
| Network 6 | Band mass: $\beta$ | -0.065 | 0.034 | 6.93e-02 |
| Network 6 | Band mass: $\gamma$ | 0.133 | 0.039 | 1.15e-03 |
| Network 7 | Network strength ($\lambda^{\text{fMRI}}$) | -0.261 | 0.031 | 3.72e-16 |
| Network 7 | Spectral centroid ($\mu^{\text{spec}}$) | 0.301 | 0.033 | 5.28e-18 |
| Network 7 | Band mass: $\theta$ | -0.146 | 0.032 | 1.09e-05 |
| Network 7 | Band mass: $\alpha$ | -0.113 | 0.032 | 7.88e-04 |
| Network 7 | Band mass: $\beta$ | -0.024 | 0.035 | 5.18e-01 |
| Network 7 | Band mass: $\gamma$ | 0.218 | 0.043 | 1.69e-06 |
| Network 8 | Network strength ($\lambda^{\text{fMRI}}$) | -0.292 | 0.031 | 7.45e-19 |
| Network 8 | Spectral centroid ($\mu^{\text{spec}}$) | 0.317 | 0.033 | 1.02e-19 |
| Network 8 | Band mass: $\theta$ | -0.129 | 0.034 | 3.00e-04 |
| Network 8 | Band mass: $\alpha$ | -0.088 | 0.035 | 1.63e-02 |
| Network 8 | Band mass: $\beta$ | 0.010 | 0.037 | 8.18e-01 |
| Network 8 | Band mass: $\gamma$ | 0.211 | 0.039 | 2.24e-07 |
| Network 9 | Network strength ($\lambda^{\text{fMRI}}$) | -0.203 | 0.031 | 2.15e-10 |
| Network 9 | Spectral centroid ($\mu^{\text{spec}}$) | 0.258 | 0.038 | 7.45e-11 |
| Network 9 | Band mass: $\theta$ | -0.127 | 0.033 | 2.56e-04 |
| Network 9 | Band mass: $\alpha$ | -0.110 | 0.033 | 1.26e-03 |
| Network 9 | Band mass: $\beta$ | -0.036 | 0.034 | 3.27e-01 |
| Network 9 | Band mass: $\gamma$ | 0.226 | 0.039 | 3.73e-08 |
| Network 10 | Network strength ($\lambda^{\text{fMRI}}$) | -0.280 | 0.033 | 2.04e-16 |
| Network 10 | Spectral centroid ($\mu^{\text{spec}}$) | 0.295 | 0.033 | 1.08e-17 |
| Network 10 | Band mass: $\theta$ | -0.134 | 0.033 | 8.17e-05 |
| Network 10 | Band mass: $\alpha$ | -0.107 | 0.033 | 1.70e-03 |
| Network 10 | Band mass: $\beta$ | -0.050 | 0.034 | 1.69e-01 |
| Network 10 | Band mass: $\gamma$ | 0.190 | 0.041 | 7.88e-06 |
| Network 11 | Network strength ($\lambda^{\text{fMRI}}$) | -0.230 | 0.037 | 2.14e-09 |
| Network 11 | Spectral centroid ($\mu^{\text{spec}}$) | 0.282 | 0.039 | 4.92e-12 |
| Network 11 | Band mass: $\theta$ | -0.136 | 0.032 | 5.74e-05 |
| Network 11 | Band mass: $\alpha$ | -0.103 | 0.033 | 2.38e-03 |
| Network 11 | Band mass: $\beta$ | -0.011 | 0.035 | 7.82e-01 |
| Network 11 | Band mass: $\gamma$ | 0.193 | 0.045 | 3.78e-05 |
| Network 12 | Network strength ($\lambda^{\text{fMRI}}$) | -0.071 | 0.044 | 1.27e-01 |
| Network 12 | Spectral centroid ($\mu^{\text{spec}}$) | 0.264 | 0.045 | 1.37e-08 |
| Network 12 | Band mass: $\theta$ | -0.150 | 0.033 | 1.48e-05 |
| Network 12 | Band mass: $\alpha$ | -0.114 | 0.034 | 1.15e-03 |
| Network 12 | Band mass: $\beta$ | -0.034 | 0.035 | 3.75e-01 |
| Network 12 | Band mass: $\gamma$ | 0.174 | 0.041 | 4.82e-05 |

**Table S4:**
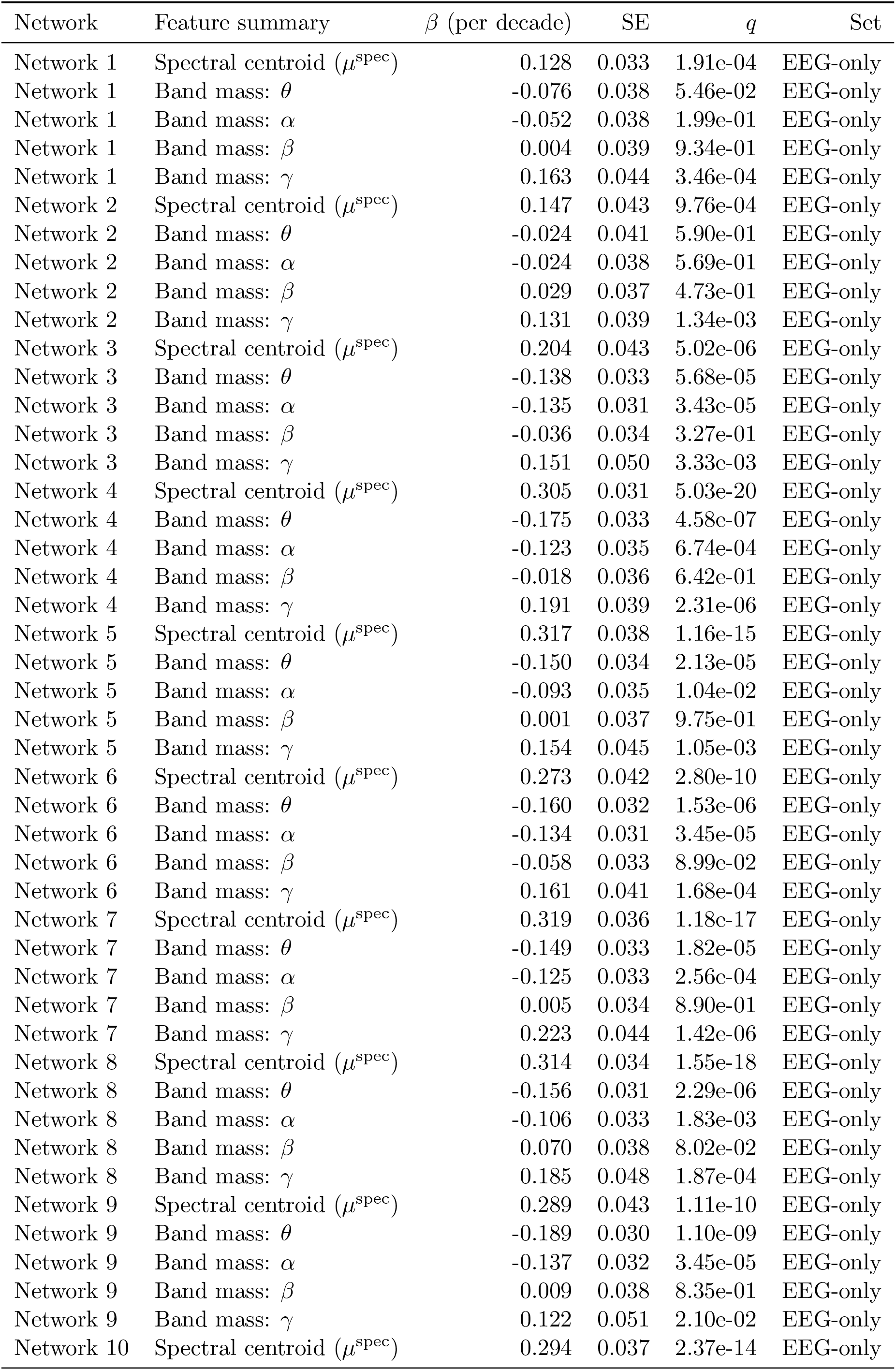
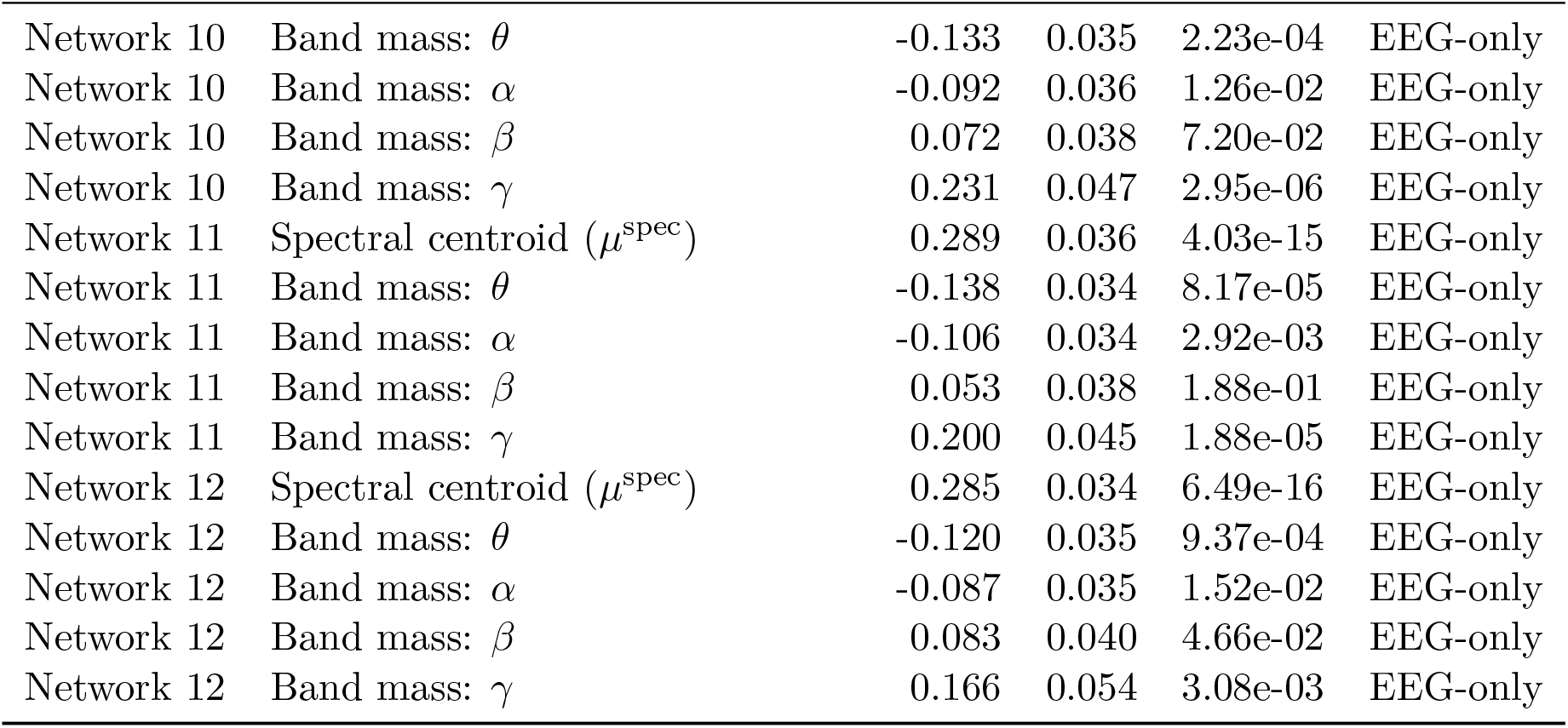
Age associations for EEG-only baseline features (per decade; SD units), adjusting for sex (male). Within-set BH-FDR *q*-values are reported. Rows are ordered by network ID (energy order) and feature type.

| Network | Feature summary | $\beta$ (per decade) | SE | $q$ | Set |
| --- | --- | --- | --- | --- | --- |
| Network 1 | Spectral centroid ($\mu^{\text{spec}}$) | 0.128 | 0.033 | 1.91e-04 | EEG-only |
| Network 1 | Band mass: $\theta$ | -0.076 | 0.038 | 5.46e-02 | EEG-only |
| Network 1 | Band mass: $\alpha$ | -0.052 | 0.038 | 1.99e-01 | EEG-only |
| Network 1 | Band mass: $\beta$ | 0.004 | 0.039 | 9.34e-01 | EEG-only |
| Network 1 | Band mass: $\gamma$ | 0.163 | 0.044 | 3.46e-04 | EEG-only |
| Network 2 | Spectral centroid ($\mu^{\text{spec}}$) | 0.147 | 0.043 | 9.76e-04 | EEG-only |
| Network 2 | Band mass: $\theta$ | -0.024 | 0.041 | 5.90e-01 | EEG-only |
| Network 2 | Band mass: $\alpha$ | -0.024 | 0.038 | 5.69e-01 | EEG-only |
| Network 2 | Band mass: $\beta$ | 0.029 | 0.037 | 4.73e-01 | EEG-only |
| Network 2 | Band mass: $\gamma$ | 0.131 | 0.039 | 1.34e-03 | EEG-only |
| Network 3 | Spectral centroid ($\mu^{\text{spec}}$) | 0.204 | 0.043 | 5.02e-06 | EEG-only |
| Network 3 | Band mass: $\theta$ | -0.138 | 0.033 | 5.68e-05 | EEG-only |
| Network 3 | Band mass: $\alpha$ | -0.135 | 0.031 | 3.43e-05 | EEG-only |
| Network 3 | Band mass: $\beta$ | -0.036 | 0.034 | 3.27e-01 | EEG-only |
| Network 3 | Band mass: $\gamma$ | 0.151 | 0.050 | 3.33e-03 | EEG-only |
| Network 4 | Spectral centroid ($\mu^{\text{spec}}$) | 0.305 | 0.031 | 5.03e-20 | EEG-only |
| Network 4 | Band mass: $\theta$ | -0.175 | 0.033 | 4.58e-07 | EEG-only |
| Network 4 | Band mass: $\alpha$ | -0.123 | 0.035 | 6.74e-04 | EEG-only |
| Network 4 | Band mass: $\beta$ | -0.018 | 0.036 | 6.42e-01 | EEG-only |
| Network 4 | Band mass: $\gamma$ | 0.191 | 0.039 | 2.31e-06 | EEG-only |
| Network 5 | Spectral centroid ($\mu^{\text{spec}}$) | 0.317 | 0.038 | 1.16e-15 | EEG-only |
| Network 5 | Band mass: $\theta$ | -0.150 | 0.034 | 2.13e-05 | EEG-only |
| Network 5 | Band mass: $\alpha$ | -0.093 | 0.035 | 1.04e-02 | EEG-only |
| Network 5 | Band mass: $\beta$ | 0.001 | 0.037 | 9.75e-01 | EEG-only |
| Network 5 | Band mass: $\gamma$ | 0.154 | 0.045 | 1.05e-03 | EEG-only |
| Network 6 | Spectral centroid ($\mu^{\text{spec}}$) | 0.273 | 0.042 | 2.80e-10 | EEG-only |
| Network 6 | Band mass: $\theta$ | -0.160 | 0.032 | 1.53e-06 | EEG-only |
| Network 6 | Band mass: $\alpha$ | -0.134 | 0.031 | 3.45e-05 | EEG-only |
| Network 6 | Band mass: $\beta$ | -0.058 | 0.033 | 8.99e-02 | EEG-only |
| Network 6 | Band mass: $\gamma$ | 0.161 | 0.041 | 1.68e-04 | EEG-only |
| Network 7 | Spectral centroid ($\mu^{\text{spec}}$) | 0.319 | 0.036 | 1.18e-17 | EEG-only |
| Network 7 | Band mass: $\theta$ | -0.149 | 0.033 | 1.82e-05 | EEG-only |
| Network 7 | Band mass: $\alpha$ | -0.125 | 0.033 | 2.56e-04 | EEG-only |
| Network 7 | Band mass: $\beta$ | 0.005 | 0.034 | 8.90e-01 | EEG-only |
| Network 7 | Band mass: $\gamma$ | 0.223 | 0.044 | 1.42e-06 | EEG-only |
| Network 8 | Spectral centroid ($\mu^{\text{spec}}$) | 0.314 | 0.034 | 1.55e-18 | EEG-only |
| Network 8 | Band mass: $\theta$ | -0.156 | 0.031 | 2.29e-06 | EEG-only |
| Network 8 | Band mass: $\alpha$ | -0.106 | 0.033 | 1.83e-03 | EEG-only |
| Network 8 | Band mass: $\beta$ | 0.070 | 0.038 | 8.02e-02 | EEG-only |
| Network 8 | Band mass: $\gamma$ | 0.185 | 0.048 | 1.87e-04 | EEG-only |
| Network 9 | Spectral centroid ($\mu^{\text{spec}}$) | 0.289 | 0.043 | 1.11e-10 | EEG-only |
| Network 9 | Band mass: $\theta$ | -0.189 | 0.030 | 1.10e-09 | EEG-only |
| Network 9 | Band mass: $\alpha$ | -0.137 | 0.032 | 3.45e-05 | EEG-only |
| Network 9 | Band mass: $\beta$ | 0.009 | 0.038 | 8.35e-01 | EEG-only |
| Network 9 | Band mass: $\gamma$ | 0.122 | 0.051 | 2.10e-02 | EEG-only |
| Network 10 | Spectral centroid ($\mu^{\text{spec}}$) | 0.294 | 0.037 | 2.37e-14 | EEG-only |
| Network 10 | Band mass: $\theta$ | -0.133 | 0.035 | 2.23e-04 | EEG-only |
| Network 10 | Band mass: $\alpha$ | -0.092 | 0.036 | 1.26e-02 | EEG-only |
| Network 10 | Band mass: $\beta$ | 0.072 | 0.038 | 7.20e-02 | EEG-only |
| Network 10 | Band mass: $\gamma$ | 0.231 | 0.047 | 2.95e-06 | EEG-only |
| Network 11 | Spectral centroid ($\mu^{\text{spec}}$) | 0.289 | 0.036 | 4.03e-15 | EEG-only |
| Network 11 | Band mass: $\theta$ | -0.138 | 0.034 | 8.17e-05 | EEG-only |
| Network 11 | Band mass: $\alpha$ | -0.106 | 0.034 | 2.92e-03 | EEG-only |
| Network 11 | Band mass: $\beta$ | 0.053 | 0.038 | 1.88e-01 | EEG-only |
| Network 11 | Band mass: $\gamma$ | 0.200 | 0.045 | 1.88e-05 | EEG-only |
| Network 12 | Spectral centroid ($\mu^{\text{spec}}$) | 0.285 | 0.034 | 6.49e-16 | EEG-only |
| Network 12 | Band mass: $\theta$ | -0.120 | 0.035 | 9.37e-04 | EEG-only |
| Network 12 | Band mass: $\alpha$ | -0.087 | 0.035 | 1.52e-02 | EEG-only |
| Network 12 | Band mass: $\beta$ | 0.083 | 0.040 | 4.66e-02 | EEG-only |
| Network 12 | Band mass: $\gamma$ | 0.166 | 0.054 | 3.08e-03 | EEG-only |

**Table S5:** Age associations for fMRI-only baseline features (per decade; SD units), adjusting for sex (male). Within-set BH-FDR *q*-values are reported. Rows are ordered by network ID (energy order).

| Network | Feature summary | $\beta$ (per decade) | SE | $q$ | Set |
| --- | --- | --- | --- | --- | --- |
| Network 1 | Network strength ($\lambda^{\text{fMRI}}$) | -0.135 | 0.040 | 1.20e-03 | FMRI-only |
| Network 2 | Network strength ($\lambda^{\text{fMRI}}$) | -0.132 | 0.036 | 3.84e-04 | FMRI-only |
| Network 3 | Network strength ($\lambda^{\text{fMRI}}$) | -0.259 | 0.031 | 1.14e-15 | FMRI-only |
| Network 4 | Network strength ($\lambda^{\text{fMRI}}$) | -0.287 | 0.032 | 7.93e-18 | FMRI-only |
| Network 5 | Network strength ($\lambda^{\text{fMRI}}$) | -0.261 | 0.032 | 2.25e-15 | FMRI-only |
| Network 6 | Network strength ($\lambda^{\text{fMRI}}$) | 0.017 | 0.043 | 7.27e-01 | FMRI-only |
| Network 7 | Network strength ($\lambda^{\text{fMRI}}$) | -0.229 | 0.036 | 7.72e-10 | FMRI-only |
| Network 8 | Network strength ($\lambda^{\text{fMRI}}$) | -0.160 | 0.034 | 6.66e-06 | FMRI-only |
| Network 9 | Network strength ($\lambda^{\text{fMRI}}$) | -0.210 | 0.036 | 2.54e-08 | FMRI-only |
| Network 10 | Network strength ($\lambda^{\text{fMRI}}$) | -0.220 | 0.039 | 6.39e-08 | FMRI-only |
| Network 11 | Network strength ($\lambda^{\text{fMRI}}$) | -0.283 | 0.030 | 9.58e-20 | FMRI-only |
| Network 12 | Network strength ($\lambda^{\text{fMRI}}$) | -0.236 | 0.036 | 3.44e-10 | FMRI-only |

**Table S6:** Age associations for all fused features derived from the Top-*K* energy-screened networks (*K* = 5), adjusting for sex and education. Each row corresponds to a univariate regression of *z*(feature) on age mid (years) plus covariates, with robust standard errors and BH-FDR correction. Effects are reported per decade (10-year change) in SD units of the feature. Rows are ordered by network ID (energy order) and feature type.

| Network | Feature summary | $\beta$ (per decade) | SE | $q$ |
| --- | --- | --- | --- | --- |
| Network 1 | Network strength ($\lambda^{\text{fMRI}}$) | -0.171 | 0.041 | 1.04e-04 |
| Network 1 | Spectral centroid ($\mu^{\text{spec}}$) | 0.198 | 0.048 | 1.04e-04 |
| Network 1 | Band mass: $\theta$ | -0.116 | 0.037 | 3.57e-03 |
| Network 1 | Band mass: $\alpha$ | -0.110 | 0.038 | 6.51e-03 |
| Network 1 | Band mass: $\beta$ | -0.060 | 0.041 | 1.70e-01 |
| Network 1 | Band mass: $\gamma$ | 0.147 | 0.060 | 2.19e-02 |
| Network 2 | Network strength ($\lambda^{\text{fMRI}}$) | -0.144 | 0.057 | 1.84e-02 |
| Network 2 | Spectral centroid ($\mu^{\text{spec}}$) | 0.268 | 0.036 | 3.79e-13 |
| Network 2 | Band mass: $\theta$ | -0.129 | 0.038 | 1.54e-03 |
| Network 2 | Band mass: $\alpha$ | -0.080 | 0.043 | 8.55e-02 |
| Network 2 | Band mass: $\beta$ | 0.004 | 0.050 | 9.45e-01 |
| Network 2 | Band mass: $\gamma$ | 0.219 | 0.055 | 2.00e-04 |
| Network 3 | Network strength ($\lambda^{\text{fMRI}}$) | -0.299 | 0.038 | 4.74e-14 |
| Network 3 | Spectral centroid ($\mu^{\text{spec}}$) | 0.243 | 0.042 | 3.58e-08 |
| Network 3 | Band mass: $\theta$ | -0.118 | 0.036 | 2.50e-03 |
| Network 3 | Band mass: $\alpha$ | -0.086 | 0.039 | 3.86e-02 |
| Network 3 | Band mass: $\beta$ | -0.008 | 0.046 | 8.85e-01 |
| Network 3 | Band mass: $\gamma$ | 0.207 | 0.077 | 1.24e-02 |
| Network 4 | Network strength ($\lambda^{\text{fMRI}}$) | -0.242 | 0.039 | 2.58e-09 |
| Network 4 | Spectral centroid ($\mu^{\text{spec}}$) | 0.288 | 0.041 | 2.69e-11 |
| Network 4 | Band mass: $\theta$ | -0.091 | 0.046 | 6.36e-02 |
| Network 4 | Band mass: $\alpha$ | -0.054 | 0.047 | 3.14e-01 |
| Network 4 | Band mass: $\beta$ | 0.029 | 0.051 | 6.25e-01 |
| Network 4 | Band mass: $\gamma$ | 0.185 | 0.046 | 1.77e-04 |
| Network 5 | Network strength ($\lambda^{\text{fMRI}}$) | -0.162 | 0.046 | 1.08e-03 |
| Network 5 | Spectral centroid ($\mu^{\text{spec}}$) | 0.316 | 0.040 | 4.74e-14 |
| Network 5 | Band mass: $\theta$ | -0.128 | 0.040 | 2.96e-03 |
| Network 5 | Band mass: $\alpha$ | -0.077 | 0.046 | 1.23e-01 |
| Network 5 | Band mass: $\beta$ | 0.036 | 0.055 | 5.77e-01 |
| Network 5 | Band mass: $\gamma$ | 0.220 | 0.062 | 9.46e-04 |

### S8.3. Additional network visualizations (networks 6–12)

#### Surface maps for all networks

To complement the main-text visualizations of the highest-energy networks (Fig. 6), we provide surface renderings for the remaining fused networks in the selected rank-*R* = 12 model. Figure S8 shows networks *r* = 6, …, 12 (ordered by median total factor energy), displayed on `fsaverage5` with lateral and medial views for both hemispheres. Visualization conventions match the main text: each network map is displayed up to an arbitrary global sign (fixed only for consistent red/blue polarity across panels) and shown with the same mild thresholding and color scaling conventions.

#### Spectral profiles and subject-level summaries (networks 6–12)

Figure S9 reports spectral profiles and subject-level score distributions for networks *r* = 6, …, 12, analogous to the main-text summary for the Top-5 networks (Fig. 7). Specifically, we show the median and interquartile range of the fitted subject-specific EEG weight curves 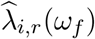 across the modeling frequency grid, alongside distributions of the derived subject-level summaries (network strength from fMRI and spectral centroid from EEG).

#### Age trends beyond the Top-5 networks

Figure S10 provides age scatterplots for networks *r* = 6, …, 12 using the same summaries as in the main text (Fig. 8). These plots serve as a qualitative check that age-related variation is not confined to the Top-5 networks and help contextualize the full association tables reported in Table S3.

**Figure S8:**
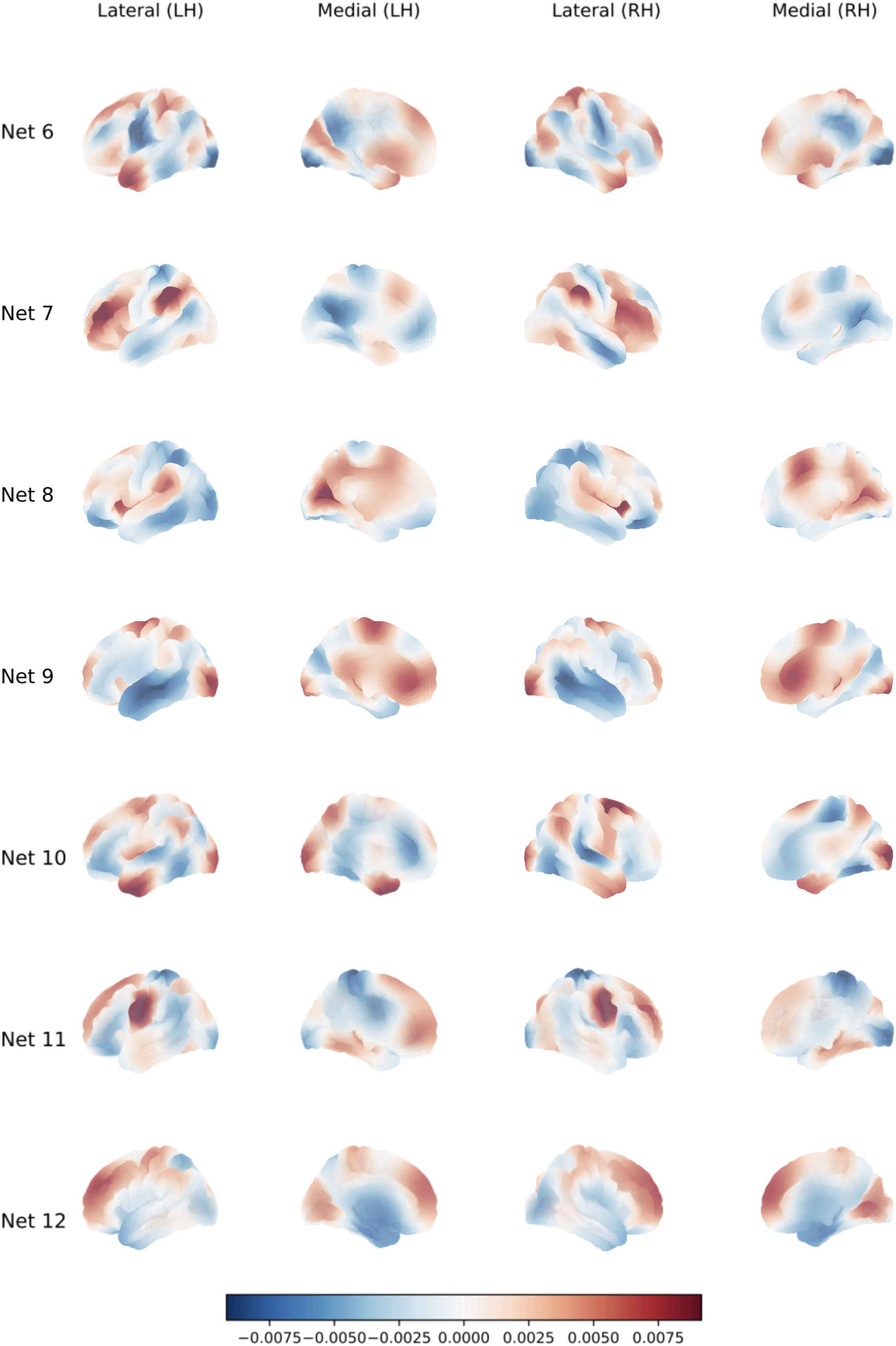
MPI–LEMON: surface visualization of fused spatial networks 6–12 (rank-*R* = 12), displayed on `fsaverage5`. Each row corresponds to one network (*r* = 6, …, 12; ordered by median total factor energy), and columns show lateral and medial views for left and right hemispheres.

**Figure S9:**
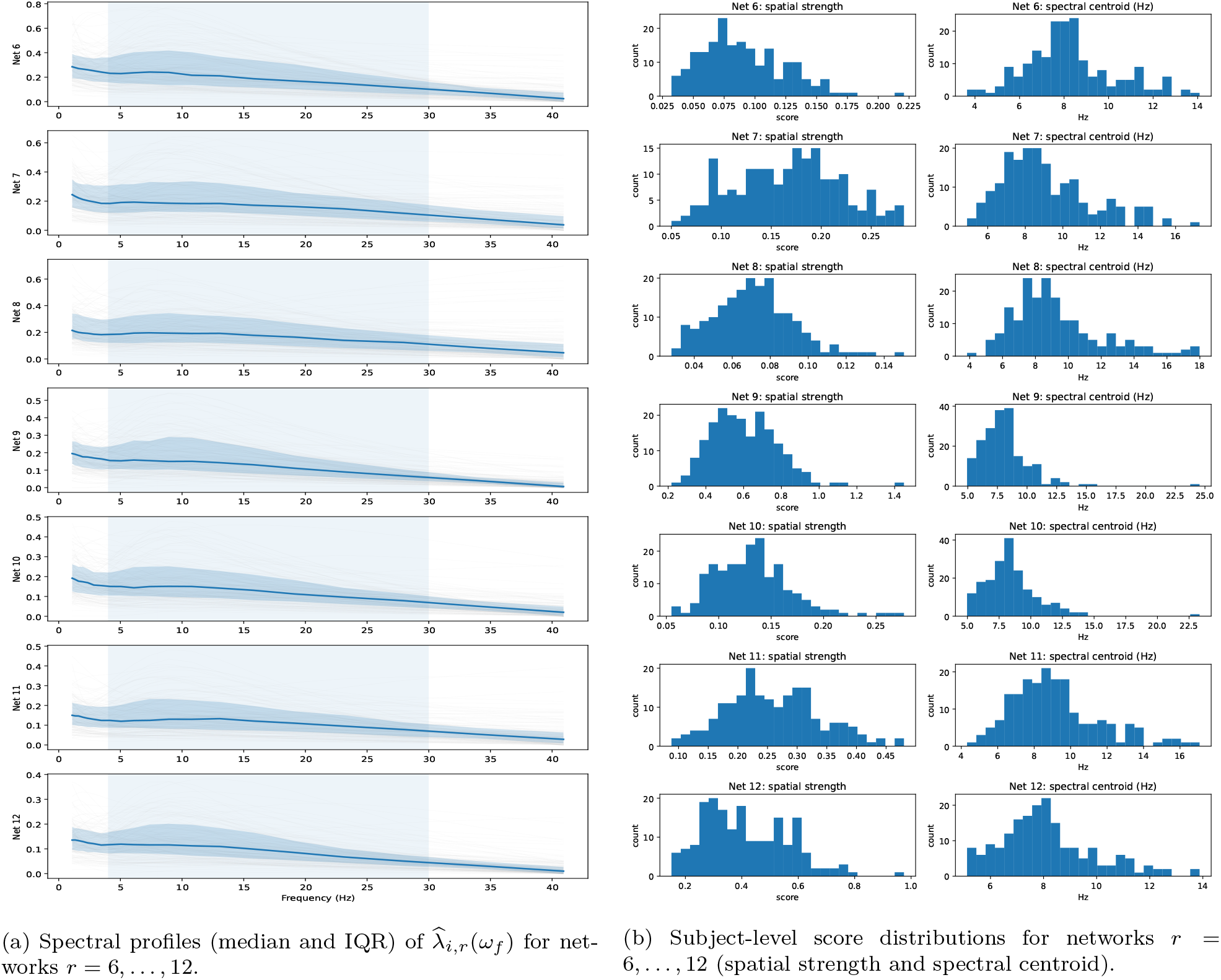
MPI–LEMON: spectral and subject-level summaries for fused networks 6–12 (rank-*R* = 12). **Left:** distribution of subject-specific EEG weight curves 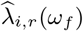 over the modeling frequency grid, shown as the median across subjects with an interquartile range (25th–75th percentile) band. **Right:** histograms of subject-level summaries for the same networks, including the fMRI-derived network-strength score (one scalar per subject and network) and the EEG-derived spectral centroid (Hz-weighted center of mass of 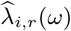.

**Figure S10:**
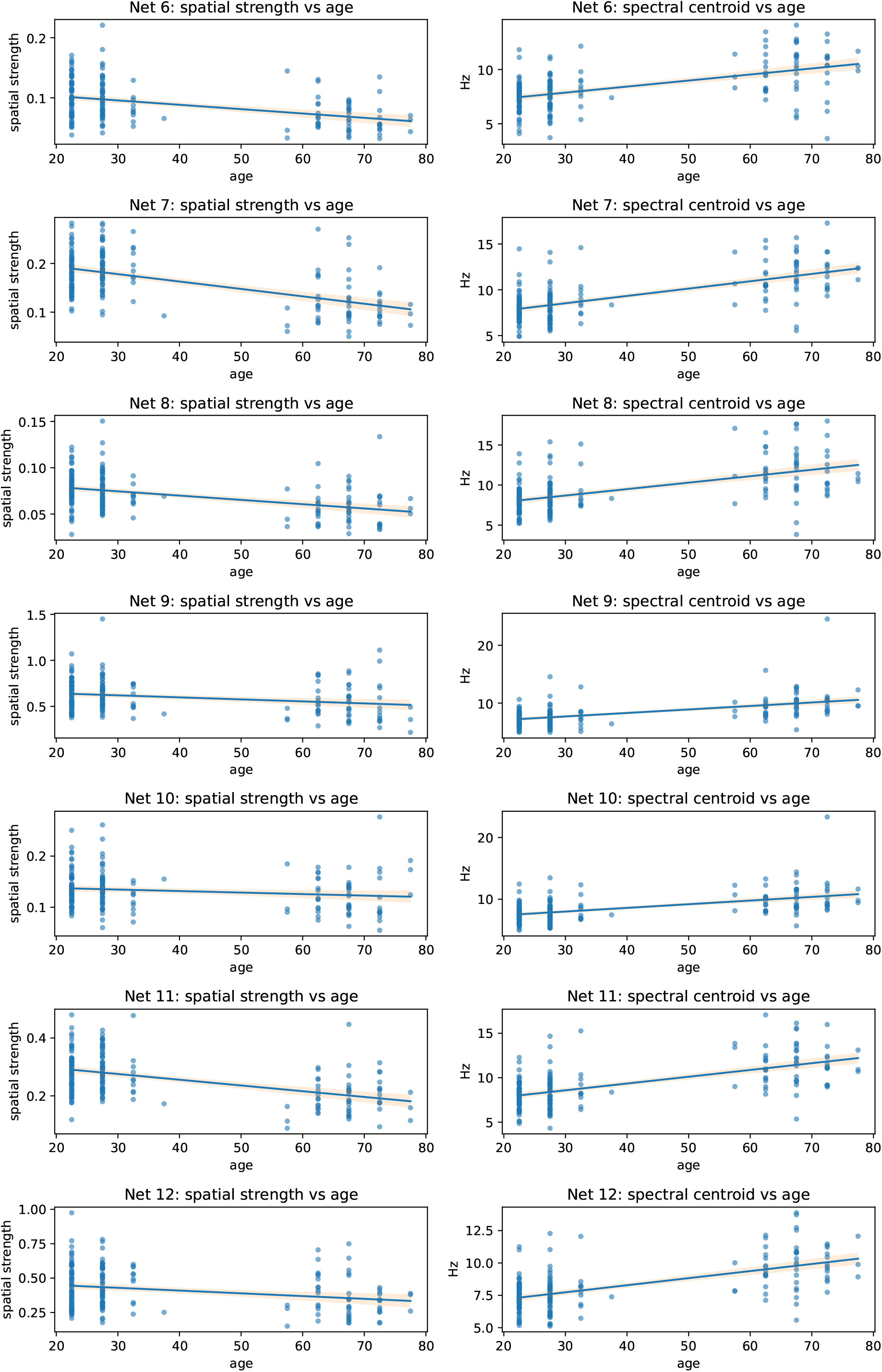
MPI–LEMON: age relationships for fused network summaries for networks 6–12 (rank-*R* = 12). For each network, we show scatter plots of subject age (midpoint of age bin) versus (left) fMRI-derived network strength and (right) EEG-derived spectral centroid, with linear fit lines (and confidence bands when shown). These plots provide a qualitative complement to the univariate age-association screening by illustrating the direction and variability of age trends beyond the Top-5 networks emphasized in the main text.

## Footnotes

1 The exact list is provided in `canonical_59.txt` and reproduced here: Fp1, Fp2, F7, F3, Fz, F4, F8, FC5, FC1, FC2, FC6, C3, Cz, C4, T8, CP5, CP1, CP2, CP6, AFz, P7, P3, Pz, P4, P8, PO9, O1, Oz, O2, PO10, AF7, AF3, AF4, AF8, F5, F1, F2, F6, FT7, FC3, FC4, FT8, C5, C1, C2, C6, CP3, CPz, CP4, TP8, P5, P1, P2, P6, PO7, PO3, POz, PO4, PO8.

